# A new archaeal virus that suppresses the transcription of host immunity genes

**DOI:** 10.1101/2024.02.12.579488

**Authors:** Israela Turgeman-Grott, Noam Golan, Uri Neri, Doron Naki, Neta Altman-Price, Kim Eizenshtein, Deepak K. Choudhary, Rachel Levy, Yarden Shalev, Himani Himani, Leah Reshef, Uri Gophna

**Affiliations:** The Shmunis School of Biomedicine and Cancer Research, Faculty of Life Sciences, Tel Aviv University, Tel Aviv, Israel; The Avinoam Adam department of natural sciences, The Open University of Israel, Raanana, Israel

## Abstract

In some extreme environments, archaeal cells have been shown to have chronic viral infections, and such infections are well-tolerated by the hosts and may potentially protect against more lethal infections by lytic viruses. We have discovered that a natural *Haloferax* strain (48N), which is closely related to the model organism *Haloferax volcanii,* is chronically-infected by a lemon-shaped virus, which we could purify from the medium. The chronic infection by this virus, which we named LSV-48N, is never cleared, despite the multiple defense systems of the host that include CRISPR-Cas, and two CBASS systems. Curing 48N of its virus by genetic engineering, led to radical changes in the gene expression profile of 48N and a dramatic improvement in its growth rate. Remarkably, the cured 48N is the fastest-growing haloarchaeon reported to date, with a generation time of approximately 1 hour at 45°C compared to the typical 2.5 hours of *H. volcanii* or its infected isogen, and faster than any known haloarchaeon. The virus subverts host defenses by reducing their transcription and interfering with the CRISPR spacer acquisition machinery. Our results suggest that the slow growth of many halophilic archaea could be due to the effects of proviruses within their genomes that consume resources and alter the gene expression of their hosts.

## Introduction

A major challenge in microbiology is to understand virus-host interactions that involve chronic infections, where cells maintain their viability yet release virions over extended periods without clearing the infection. Viral infections are expected to lead to a major loss of fitness for the infected strains, and consequently to their demise, yet in some environments such infections can be paradoxically quite common ^1,2^. Archaeal viruses represent a true frontier for virology research, showing an enormous diversity of morphology, egress mechanisms ^3^, and molecular biology. Ecologically, the impact of archaeal viruses on their hosts, in some ecosystems such as deep marine sediments, may be greater than that of bacteriophages ^4^, yet relatively few of them have been cultivated ^2,5–7^, compared to bacteriophages or eukaryotic viruses. Chronic infections appear to be common in archaea, and the viruses that cause such infections are often challenging to isolate since they do not form plaques on soft agar plates. Past studies of such viruses have thus relied on metagenomics and single-cell genomics to identify and quantify host-virus interactions in archaea-dominated ecosystems. These approaches have led to major discoveries, including the observation that in some extreme environments >60% of cells are infected with viruses and most of them contain two or more virus types ^1^. This has led to the hypothesis that in some natural environments lysogeny (where a virus is completely dormant) or a relatively mild chronic infection is not just tolerable, but may actually be beneficial when it offers protection to cells against more virulent viruses ^1,2^. Moreover, it has also been shown in thermo-acidophilic archaea that chronic infection can be maintained when infected cells produced a toxin that kills off competing non-infected strains, as shown ^8^.

In haloarchaea, chronic viral infections are common ^9–11^, yet unlike lytic infection ^12^, there was no good model system to study chronic infection until very recently. The newly established system of *Haloferax volcanii* pleomorphic virus 1 ^9^, uses a virus isolated from an environmental sample, whose original host remains unknown, to successfully infect the lab strain of *H. volcanii* (DS2), the best studied archaeal model organism ^13^. This virus has a relatively minor effect on host growth, in line with the “tolerable infection” scenario.

Many archaea harbor multiple defense systems, including CRISPR-Cas systems, which provide acquired immunity. Chronically infecting viruses must thus avoid triggering such defenses, or subvert them, in order to establish a long-term coexistence with their hosts. In the case of *H. volcanii* pleomorphic virus 1, lack of CRISPR-mediated acquired immunity by DS2 during exponential growth was attributed to a two-fold down-regulation of the acquisition module gene *cas1*, yet no down-regulation was observed during stationary phase, despite sustained virus production ^9^.

Here we study the interaction of a chronically infecting lemon-shaped virus and its host the halophilic archaeon *Haloferax volcanii* 48N (henceforth 48N), which is closely related to the model organism *H. volcanii* (97.91% average nucleotide identity in coding genes). By using a “gene therapy” approach in which the provirus was deleted from the genome we were able to cure 48N of its virus, and compare phenotypes and gene expression of the cured and infected strains. We observed that under chronic infection, growth is dramatically delayed and anti-viral defense systems are transcriptionally repressed, and the CRISPR-Cas-based acquired immunity is disrupted.

## Results and Discussion

### Haloferax volcanii DS2 acquires CRISPR spacers against a region within the 48N genome that encodes a provirus

48N is a strain originally isolated from the tidal pools of Atlit for which a draft genome is available ^14,15^. We performed a mating experiment between a uracil auxotroph derivative of 48N (UG613, supplementary Table T3) that carries the pWL102 plasmid conferring resistance to mevinolin and a tryptophan auxotroph derivative of *Haloferax volcanii* DS2 (UG469). This assay, designed to obtain mostly colonies of UG469 that received pWL102 by mating, was conducted by applying stationary cultures of the two strains onto a filter on a rich medium and then transferring them to selectable plates containing mevinolin and lacking uracil. We then tested the spacer acquisition profile of the CRISPR-Cas system in the UG469 mating products and observed that spacer acquisition was highly skewed: the vast majority of spacers matched just one region in the 48N genome (Fig. 1A; Supplementary Table T1). A closer examination of that genome region showed that it contained ORFs encoding two homologs of bacteriophage-associated integrases that are typical to temperate phages. The same region also had matches in the CRISPR arrays of two *Haloferax* strains isolated in the same year from the same site as 48N: *Haloferax* strains 24N and 47N (Table 1), which did not contain the putative virus locus in their genomes. Most notably, in both of these strains, the leader proximal spacer, which in type I systems is the one most recently acquired, matched the suspected provirus (Table 1). The pattern of acquisition resembled that previously observed in bacteria following infection with dsDNA phages ^16^, i.e. a strong enrichment for a specific part of the viral genome, potentially representing the free end of the injected DNA (Fig. 1A). Taken together these observations led us to hypothesize that spacers were acquired from a virus that chronically infected 48N and was able to deliver its DNA into strain UG469, and probably some environmental strains of *Haloferax,* but failed to establish an infection in those strains.

**Figure 1.**
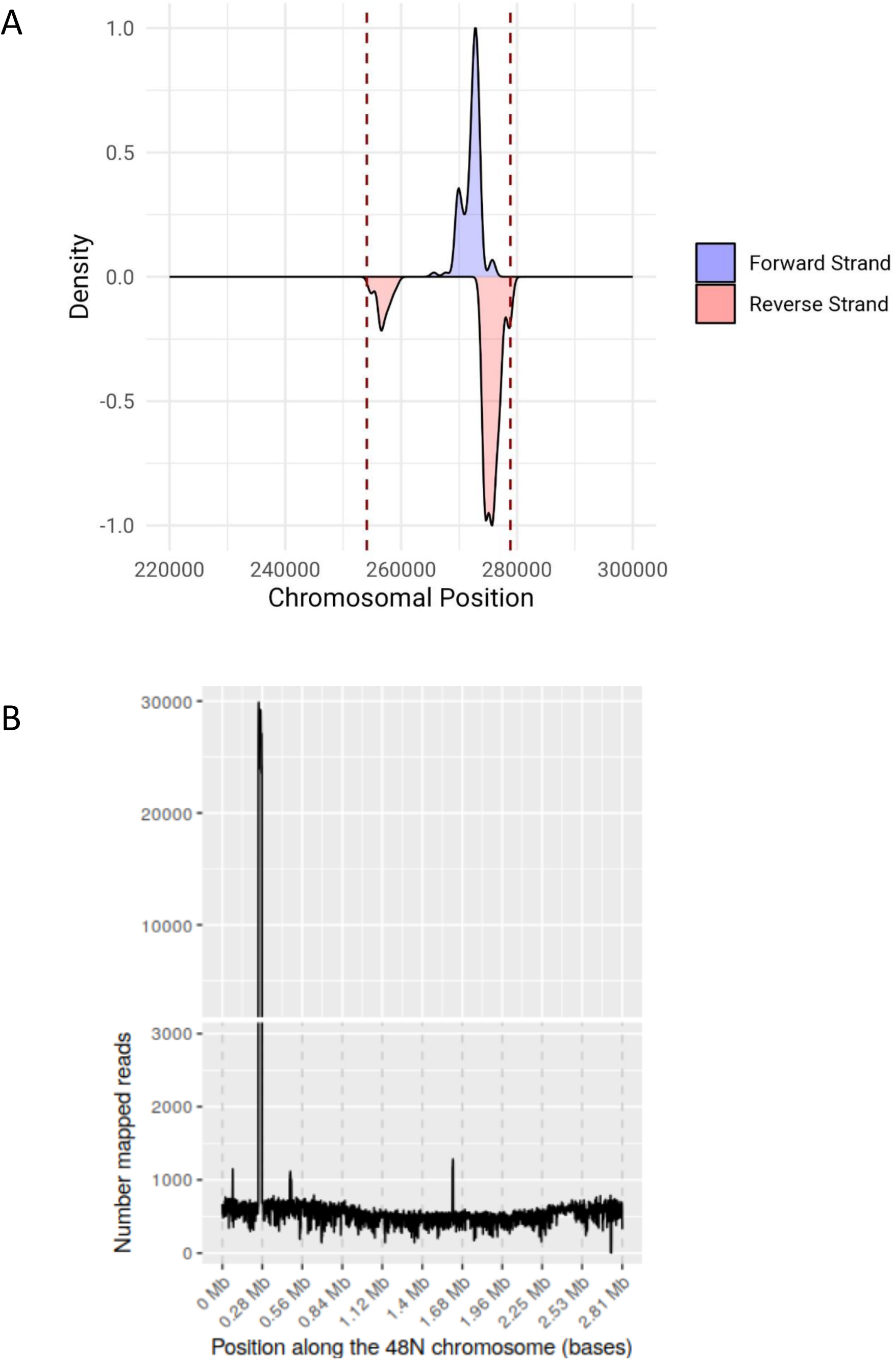
Genomics-based evidence for the existence of a virus in 48N. A. CRISPR spacer acquisition against the 48N chromosome by *H. volcanii* during mating on filters is almost exclusively against one locus in the genome. B. Coverage plot of Illumina sequence reads that map to the 48N chromosome. DNA was extracted from cells grown to high cell density over 3 days (stationary culture).

**Table 1.**
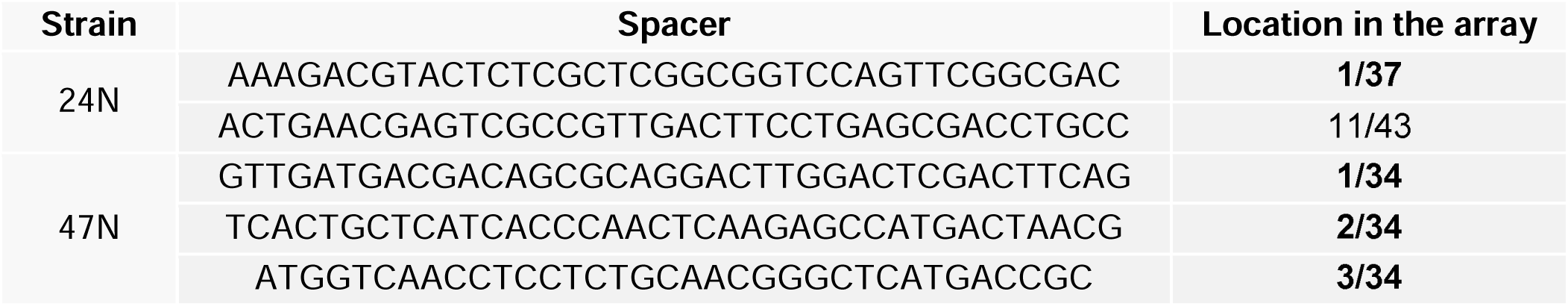
Spacers matching the provirus locus from 48N in genomes of natural isolates obtained from the same site and season. Leader-proximal spacer locations are highlighted in bold type)

To test whether the suspected genomic region indeed originated from a replicating mobile element, we first obtained a fully assembled genome of 48N main chromosome (for which only a draft was available) along with its four natural plasmids, by combining the previous Illumina reads with newly generated Pacific BioScience SMRT sequencing data (see Methods). We then mapped the raw Illumina reads to the genome and obtained coverage scores of different areas of the genome. This analysis showed that coverage of the suspected provirus locus was over 20-fold higher than the genome average (Figure 1B). Notably, examination of the assembly graph showed that the pro-virus element exists both as an integrated provirus and in a circular form. In addition to the CRISPR-targeted proviral region, three other genomic regions within the 48N main chromosome had sequence coverage of about 2.5-3.5 fold - higher than the mean coverage, indicating that additional replicating elements are present in the 48N culture (Fig. 1B).

### 48N encodes a chronically infecting lemon-shaped virus

To test whether the replicating element we identified is indeed a virus, we concentrated the supernatant of a 48N culture and submitted it to transmission electron microscopy (TEM). TEM images revealed the presence of predominantly lemon-shaped virus-like particles about 65-90 nm long and 35-50 nm wide (Fig. 2 A-C), similar in morphology to lemon|(spindle)-shaped viruses previously observed in other archaeal genera such as *Haloarcula* ^17,18^ and *Sulfolobus/Saccharolobus* ^19–21^, despite the lack of sequence similarity of any of the ORFs encoded by the provirus to any known capsid protein. Both intracellular viruses and a fraction of the secreted ones appeared attached to fibrillar structures (blue dashed arrows, Fig. 2C and 2E). The supernatant also contained additional spherical virus-like particles about 75-90 nm in diameter (Fig. 2D). When we extracted DNA from the supernatant and submitted it to Illumina sequencing and de-novo genome assembly (see Methods), we observed: numerous small (>5kbp) contigs of relatively low coverage depth (averaging between 0.95 and 4.82 fold coverage) mapping to the host chromosome and plasmids, and two prominent contigs, 24kbp and 16kbp long, of higher coverage depth (7403.05 and 17.29 average fold, respectively (Supplementary Table T2), The 24kbp contig corresponded to a circular replicon. The 16kbp contig encoded two putative proteins that showed similarity to proteins conserved in haloarchaeal pleomorphic viruses (one matching ORF7 of *Halogeometricum* sp. pleomorphic virus 1 ^22,23^, and the other ORF6 of that virus with a closer match to *Haloferax* pleomorphic virus 1 ^9^. Additionally, the 16kbp contig encoded a tyrosine recombinase/integrase typical to archaeal proviruses and temperate bacteriophages that have double-stranded DNA genomes. This implies that the 16kbp contig likely belongs to a pleomorphic virus, while the 24kbp genome corresponds to a lemon-shaped virus (henceforth LSV-48N). LSV-48N virions were also observed within 48N cells (Fig. 2E).

**Figure 2.**
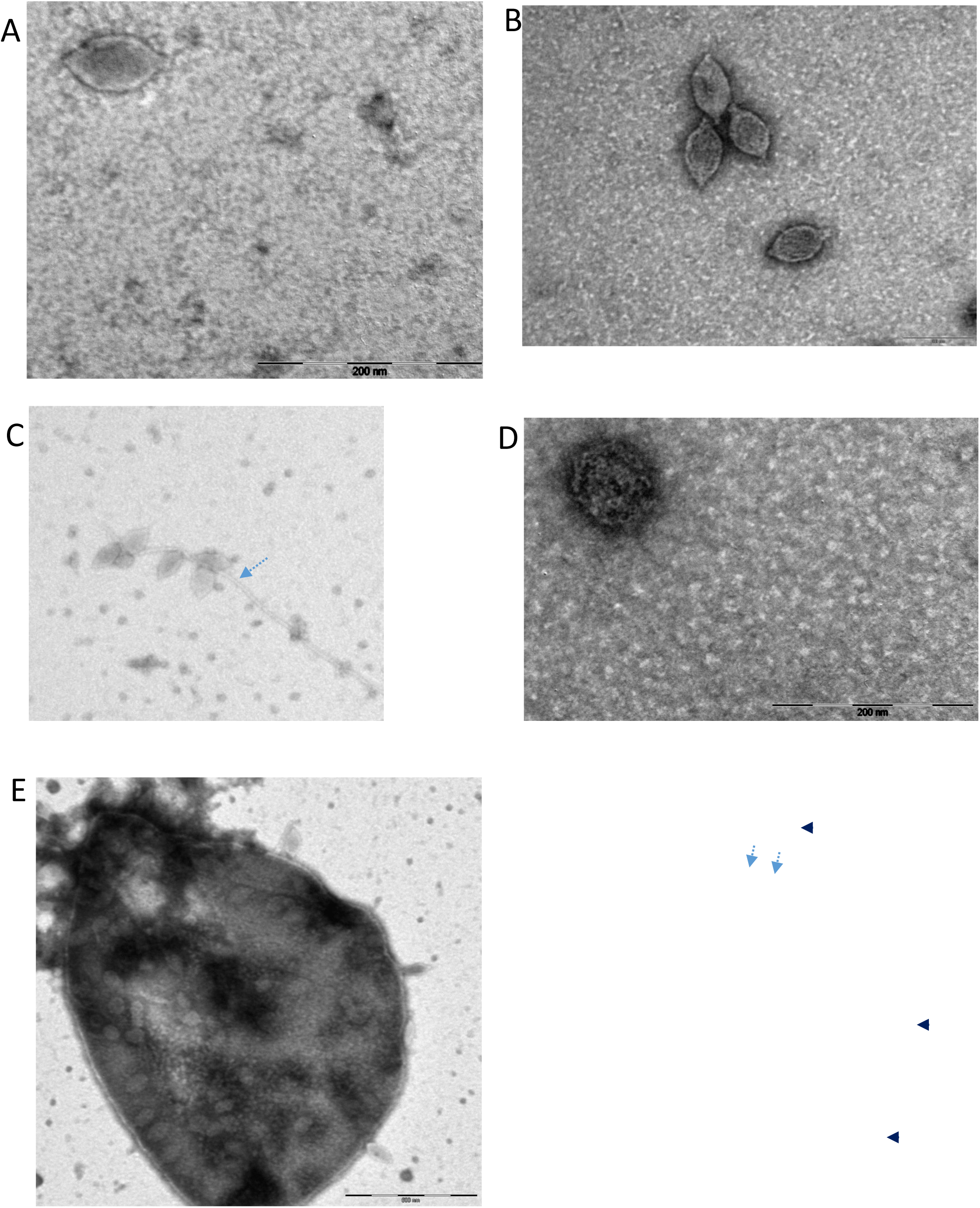
Viruses infecting *Haloferax* Atlit 48N. A-C. Transmission electron micrographs of lemon-shaped viruses released to the supernatant. D. A pleomorphic virus present in the same supernatant preparation. E. Lemon-shaped viruses outside (black triangles) and within cells. Dashed blue arrows mark fibrillar structures that some of the viruses adhered to.

### LSV-48N is non-lethal and is continuously shed during growth

We used outward-facing primers that can amplify a circularized genome or primers that amplify the edges of the integrated form of the virus (Supplementary Table T4) for PCR-based analysis of DNA extracted from 48N cultures. This analysis indicated that both circular and integrated forms are present under standard, lower-salt and low-phosphate conditions (Supplementary Figure S1). In order to determine at what stages during 48N growth curve are viruses produced and secreted, we used droplet digital PCR (ddPCR) to quantify viral genome copies in DNA extracted from the supernatant as well as from the cells at different time points when growing 48N in rich liquid medium (see Methods). Results showed that during early exponential growth (OD of 0.1) there is maximal accumulation of viral genome copies within cells, yet the maximal release of viruses to the medium occurs during late exponential growth-early stationary phase (OD of 0.7-1). (Supplementary Figure S2). When the viability of the cells was tested using fluorescent microscopy and the LIVE/DEAD™ BacLight™ kit, there were very few dead cells present (roughly 1.5%). Furthermore, 48N consistently grew faster than *H. volcanii* lab strain (Fig. 3A), indicating that chronic infection with LSV-48N is well-tolerated by the host.

**Figure 3.**
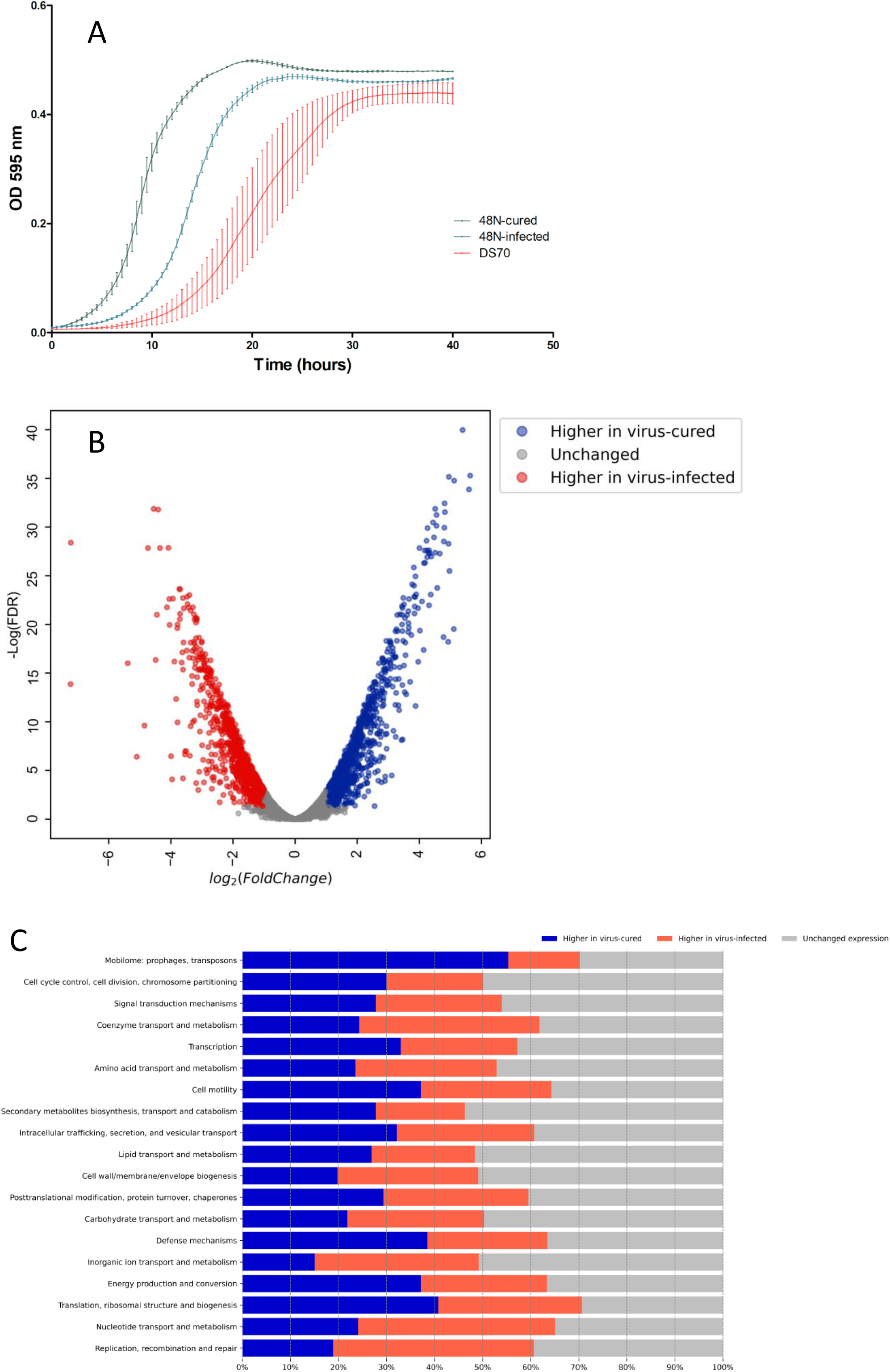
Effects of virus curing on gene expression and growth of the host. A. growth curves of 48N strains (UG461 and its virus-cured derivative) and the *H. volcanii* lab strain. Samples were grown over-night in YPC rich medium at 45°C to the mid-log phase and then diluted with a fresh medium to OD_595nm_ of 0.1. The growth curves were carried out in microplate reader at 45°C with continuous shaking while measurement were taken every 30 minutes at a wavelength of 595nm. B. A volcano plot showing differentially abundant transcripts in cultures of 48N and virus-cured 48N, based on RNA-Seq. C. Gene categories affected by infection with LSV-48N.

### LSV-48N is a representative of a new family of viral elements in Haloferax

Since LSV-48N showed no significant sequence similarity to known haloarchaeal viruses, we used sequence comparisons of six-frame translations of LSV-48N sequences to compare with available haloarchaeal genomes to identify related integrated proviruses. We detected several related proviruses in other strains and species of *Haloferax* that had notable similarities in gene organization and sequence to LSV-48N (Fig. 4). Given that nearly all LSV-48N genes had homologs in these proviruses we conclude that LSV-48N is the first studied representative of a new family of viruses able to integrate into the genomes of *Haloferax* species.

**Figure 4.**
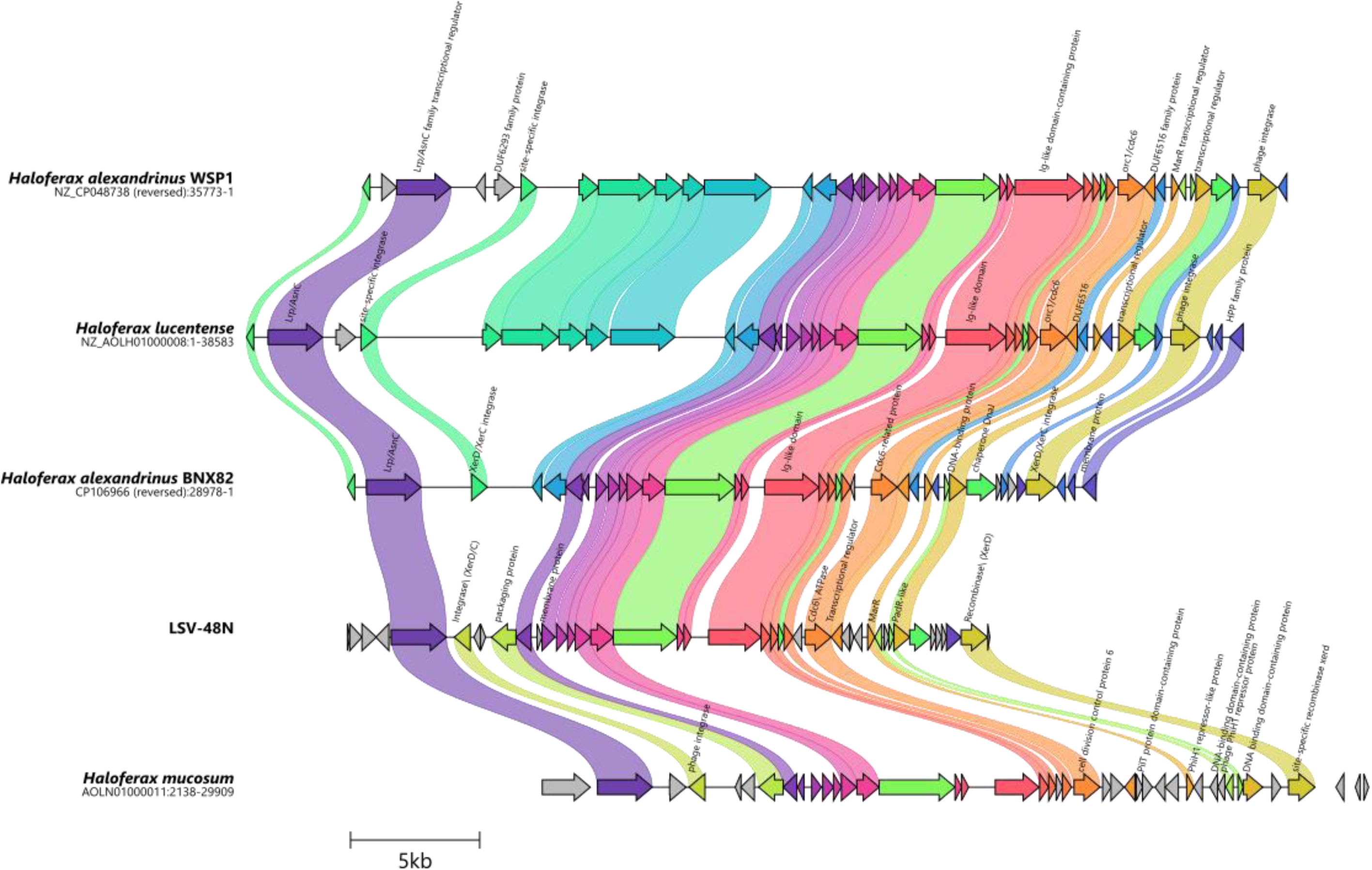
Comparative synteny plots of putative *Haloferax* proviruses. Gene cluster plot was generated with clinker ^46^. Each color represents a gene family and linked protein-coding genes have over 50% pair-wise sequence identity.

### LSV-48N profoundly affects the growth and physiology of its host

Even when grown for 400 generations in rich medium, no colonies that spontaneously lost LSV-48N (tested by virus-specific colony PCR) could be detected. We therefore deleted the provirus from the genome using a two-step strategy, beginning from UG613. First, we deleted the gene for the putative phage-like integrase from UG613 using the pop-in/pop-out approach. Then, we deleted the remaining part of the viral genome using the same method and confirmed the clean deletion via whole genome sequencing. As expected, TEM images showed lemon-shaped viruses in the supernatant of the infected 48N/l/*pyrE2* strain but not in the virus-cured strain (cured 48N), while the spherical virus-like particles were still present.

Chronic viral infection often results in a considerable growth burden and a major shift of the host gene expression (Alarcón-Schumacher*, et al.*, 2022). Indeed, when we compared the growth of the wild-type infected 48N to its cured 48N, we observed a major growth improvement, with a growth rate over two-fold higher, the fastest to our knowledge ever observed for a haloarchaeon ^24^ at 45LJLJL (Fig.3A, Supplementary video V1). We then extracted RNA from both naturally-infected and cured strains and compared their transcriptomes using RNA-seq. As can be expected from such striking differences in growth, RNA-seq analysis revealed an immense difference in the transcription pattern of 48N, with almost 60% of the genes in cured 48N exhibiting a significant difference in their expression (Fig. 3B) that spanned all functional categories (Fig. 3C). The large increase in generation time in the virus-infected strain indicates that the virus is consuming much of the cellular resources and potentially modulating host metabolism. The RNA-seq results showed that not only were viral genes encoding membrane proteins very highly transcribed (Supplementary Fig. S3), but the expression of many host genes which are associated with different functions, including metabolism, were unexpectedly altered. Thus, the two cytochrome bd-type quinol oxidase subunits that play a critical role in respiration in haloarchaea were more than 10-fold higher in the infected strain, implying that respiration may be increased during infection. In contrast, many host metabolic genes were strongly downregulated, such as the archaeal-type H+ ATPase subunits, nearly all of which had mRNA levels decrease of more than 15 fold.

### LSV downregulates the defense systems of its host

Based on genomic analysis using the PADLOC server ^25^, the 48N genome contains multiple potential anti-viral defenses encoded on its natural plasmids. Among those, CRISPR-Cas and the two CBASS systems should be able to prevent chronic viral infection: CRISPR-Cas can generally acquire spacers against replicating selfish elements and then target their DNA for degradation, while CBASS systems cause abortive-infection by recognizing viral infection and killing the host ^26^. However, a comparison of the expression levels of defense genes in 48N and cured-48N showed that these defense systems, and additional ones, such as Hachiman, were strongly downregulated in the infected strain (Table 2). Notably, the genes encoding the cyclases of the CBASS systems, required for their activation in the presence of viral infection had much lower levels in 48N compared to cured 48N (8.29 and 14.6 fold lower in CBASS type 1 and 2, respectively), as were the cas2 and cas4, genes required for new spacer acquisition (17.19 and 4.68 lower, respectively). Thus, LSV-48N infection downregulates cellular defenses, either directly or indirectly at the level of transcription, sufficiently to allow chronic and sustained infection.

**Table 2.** Anti-viral immunity genes whose expression is significantly higher in the cured strain vs. its infected isogen.

| Predicted protein encoded by the gene | Level in cured (RPKM) | Level in Infected (RPKM) | Fold increase (cured) |
| --- | --- | --- | --- |
| **CRISPR- associated protein Cas7** | 205.31 | 53.85 | 3.81 |
| **CRISPR- associated protein Cas5** | 146.69 | 25.36 | 5.79 |
| **CRISPR-associated protein Cas4** | 95.30 | 20.38 | 4.68 |
| **CRISPR-associated protein Cas2** | 126.33 | 7.35 | 17.19 |
| **CBASS_1_Cyclase** | 45.85 | 5.53 | 8.29 |
| **CBASS_1_Effector** | 66.68 | 23.89 | 2.79 |
| **CBASS_2 Cyclase** | 343.63 | 23.56 | 14.58 |
| **DUF1837 domain-containing protein / Hachiman A1** | 347.90 | 13.25 | 26.25 |
| **DEAD/DEAH box helicase / Hachiman B1** | 222.52 | 30.53 | 7.29 |

To further investigate why 48N does not acquire CRISPR immunity against LSV-48N, we performed a spacer acquisition assay on the 48N CRISPR arrays using amplicon sequencing. We observed extremely low spacer acquisition levels and no spacers that originated from the genome of LSV-48N. Furthermore, 63% of newly acquired spacers did not have exact or near exact matches in any 48N replicon. A closer inspection of the newly acquired spacers revealed that many of them clearly resembled 48N genomic sequences but had more than five substitutions, Supplementary Figure S4). Thus, it appears that either LSV-48N or another selfish genetic element within the 48N genome encodes a factor that interferes with the spacer acquisition process of the 48N CRISPR-Cas system.

## Conclusions

The lemon(spindle) viral morphotype is unique to the domain archaea and is the most abundant particle shape in several hyper-saline ecosystems ^27,28^. Surprisingly, LSV-48N is unrelated to previously isolated haloarchaeal lemon-shaped viruses ^17,18^, but rather shares several characteristics with fuselloviruses of Thermoproteota (such as SSV1 and SSV2), despite lack of significant sequence similarity, having a circular double stranded genome, causing no cell lysis, and existing both as circular plasmid-like form and as an integrated provirus ^21,29^. Furthermore, infection by SSV1 was shown to dampen the induction of CRISPR-Cas systems in *Saccharolobus solfataricus* similar to what we show for LSV-48N ^30^. However, unlike SSV1 that has a “harmonic coexistence” with its host, with only minor effects on host gene expression (unless induced by UV), and little effect on growth ^21,29^, LSV-48N causes drastic changes in both host gene expression and growth.

Extreme environments that are often dominated by archaea tend to have lower diversity and lower cell density than bacteria-dominated hyper-diverse ecosystems such as soil or the mammalian gut. Consequently, host-virus interactions of extremophilic archaea may differ greatly from phage-bacteria interactions, and it has been suggested that when chronic infection protects against lethal infection, it can actually be beneficial. The viruses in such environments, in turn, may benefit from a chronic lifestyle when the host cell density is low, making long-term exploitation of the host and its progeny much more desirable ^31^. Here we examine the physiology and gene expression of a chronically infected extremophile strain. Importantly, because it was isolated directly from an evaporation pond where other strains of the same species abound, it can be assumed to be relatively fit, despite being infected. Furthermore, it grows faster than DS2 in the lab on standard media, and outcompetes it quickly in head-to-head growth experiments (data not shown). Nonetheless, the seemingly mild burden of chronic infection is in reality not mild at all - curing the virus dramatically increased the growth rate as well as the expression of anti-viral defense genes. While it may seem paradoxical that such a burden is sustained in a natural habitat, one should keep in mind that in many ecosystems that are continuously limited for resources, such as available carbon and nitrogen, and are not subject to feast or famine fluctuations, the maximal growth rate may not be the key trait determining ecological fitness. Furthermore, the two strains that were isolated in the same sampling round with 48N (24N and 47N) that were not infected by LSV-48N and had CRISPR spacers matching its genome, had similar growth rates to that of 48N (generation times on rich medium of 2.08 and 2.96 h. For 24N and 47N, respectively, vs. 2.18h for 48N). Thus, as observed previously for other extreme environments, hypersaline systems may allow chronic infection with viruses to be tolerated (Munson-McGee*, et al.*, 2018, Wirth & Young, 2020).

## Acknowledgements

The authors thank Shachar Robinzon for performing Haloferax mating experiments, Vered Holdengreber for help with electron microscopy and R. Thane Papke for his help with 48N genomics. This work was funded by a European Research Council grant AdG 787514.

## Methods

### Culture conditions

Haloferax strains were routinely grown aerobically at 45°C in either Hv-YPC (rich medium) or in Hv-Ca / Hv-Ca+ (minimal medium) as described in ^32^. When required, uracil was added at a concentration of 50 µg/ml. For pop-out selection medium, 5-fluoroorotic acid (5-FOA) was added to a concentration of 50 µg/ml.

### Droplet Digital PCR (ddPCR) analysis on circular LSV-48N

The number of viral particles in 1ml of 48N culture was inferred from ddPCR experiments (Bio-Rad ddPCR device according to manufacture protocol). 1ml from cultures at different growth stages (OD600nm: 0.1, 0.2, 0.3, 0.5, 0.7, 1.0) were analyzed. In order to measure the amount of secreted viral particles in the media, DNA was extracted from 200µl of the supernatant using Quick-DNA_viral_kit (Zymo Research) following centrifugation at 12,000 rpm. For “in cells” viral particles quantification, 1ml from the cell culture from each growth stage was vortexed and washed twice with Hv-Ca broth to remove cell adhesed viruses. The cells were then diluted and lysed (suspended pellet cells from 1ml with 100 µl ST buffer [1 M NaCl, 20 mM Tris.HCl] and 100 µl lysis buffer [100 mM EDTA pH 8.0, 0.2% SDS]) before viral DNA was extracted from lysed cells (Quick-DNA viral kit (Zymo Research)). Viral DNA extractions were diluted to final concentration of 0.2 pg/µl before added to the ddPCR reaction mix. Primers used are: IT24, IT25 for the circular form of the virus and: Is660 Is661 for control housekeeping gene. 48N cells were grown to OD600nm of: 0.1, 0.2, 0.3, 0.5, 0.7, 1.0. in YPC. CFU counts was were made for each culture in order to deduce the average number of archaeal cells in 1 ml.

### LSV-48N particles imaging

LSV-48N particles were visualized using a negative staining approach: Samples were adsorbed on formvar/carbon coated grids and stained with 2% aqueous uranyl acetate. Samples were examined using JEM 1400plus transmission electron microscope (Jeol, Japan).

### Growth curves

To compare the growth of the 48N strains, each sample was grown over-night in appropriate media at 45°C to the mid-log phase and then diluted to a fresh medium to OD600nm of 0.1. The growth curves were carried out in 96-well plates at 45°C with continuous shaking, using the Biotek ELX808IU-PC microplate reader. Optical density was measured every 30 minutes at a wavelength of 595nm.

### LSV-48N circular / integrated form detection by PCR

Integrated and circulated form of LSV-48N inferred using PCR analysis with specific PCR oligonucleotides. The integrated form was inferred when a PCR product of 1000 bp was visible after gel electrophoresis using IT556 and IT557 for upstream integration site and IT558 and IT559 for downstream integration site. One primer corresponds to the genome and the other one to the virus in the integration site. The circular form was obtained using IT560 and IT561 The primers matched the ends of the virus and the 1000 bp product is only obtained if the virus closes in a circle. PCR was performed using KAPA 2G ready mix by Roche.

### Live/Dead staining

To assess the viability of 48N and virus-cured 48N a Live/Dead staining experiment was performed utilizing the LIVE/DEAD BacLight Bacterial Viability Kit (Thermo-Fisher). Mixed Cultures of 48N and 48N cured were grown overnight in YPC medium to late logarithmic growth in multiple biological replicates. Subsequently, the cultures were diluted to an optical density (OD) of 0.1 and approximately 50 µl of resuspended cultures were aliquoted into 1 ml microfuge tubes. Subsequently, 10 µl of Propidium iodide dye and 10 µl of SYTO9 dye were added separately from stock solutions. The microfuge tubes were covered with aluminum foil to prevent exposure to light and minimize dye degradation. The treated cultures were then ncubated in the dark for 15-20 minutes to facilitate cellular uptake of the stains. Following the incubation period, 10 µl of the treated culture was dispensed onto a glass slide, and was visualized using a confocal microscope. Multiple images were acquired at magnifications of 20x and 40x to visualize the stained cells and assess their viability.

### Mating experiments

Liquid cultures of both parental strains were grown to an O.D600 of ∼1.8. The parental strains were then mixed in 1:1 ratio and applied to a nitrocellulose 0.45µm filters using a Swinnex 25mm filter holder. The filter with the mating products was transferred to a rich medium plate (Hv-YPC) for 24 hours for phenotypic expression. The cells were then re-suspended and washed in Hv-Ca broth before plated on Hv-Ca media containing mevinolin and lacking uracil. After 14 days in 45LC incubator, we were able to detect colonies that grew on the selectable plates. In each biological replicate, about 200 colonies were picked, and their DNA was extracted using the spooling protocol ^33^. The DNA samples were used to identify newly acquired CRISPR spacers (see below).

### New CRISPR spacers detection

Extracted DNA was used as PCR template for either *H. volcanii* or 48N CRISPR arrays using specific primers amplifying the region between the leader and the third spacer of each array (see primers list in Supplementary Table T2). When analyzed by agarose gel electrophoresis, PCR products 70 bp higher than the main PCR fragment represent potential new spacer-repeat acquisitions. We extracted from the gel the region that corresponded to amplicons of increased length (potentially having the new spacer sequences), isolated the DNA from the gel and amplified the fragment through another cycle of PCR. New acquisition events could then be detected via the presence of a higher band in the gel. PCR products were then sent for further processing and Illumina amplicon sequencing (240,000-290,000 reads per array per biological repeat) at the Center for Genomic Research, University of Illinois, USA. Briefly, the elongated PCR product was enriched using Ampure beads size-selection; sample specific barcodes and Illumina adaptors were added by PCR; and the resulting products were purified, pooled, and paired-ends sequenced on a MiSeq Illumina platform. Notably, even after these consecutive steps of size selection many reads still represented amplicons derived from no acquisition amplifications. Two *H. volcanii* arrays (C and D) were processed from five biological replicates in the vol-48N mating experiment; 3 48N arrays (A, B, C) were processed from 4 biological replicates from the 48N experiment (48N culture grown in YPC medium).

### Initial spacer data processing

As an initial quality control step, raw reads in which the amplification primers were detected either zero or multiple times were removed. Reads were then quality-filtered (Q>20) and merged using PEAR (paired-end read merger ^34^), yielding a minimum of 198000 high-quality sequences (median sample depth: 350000). Biological replicates within each array were then converted to a single multi-FASTA file using QIIME’s multiple_split_libraries_fastq.py script 13, followed by de-replication, abundance sorting and clustering (95% threshold) using VSEARCH 14. Pairwise identity at the clustering step was defined as [matching columns]/ [alignment length] (set in VSERACH as –iddef 1), which ensures terminal gaps are not ignored and sequences of different length do not cluster together. The length distribution of the clustered sequences is similar to that of the raw sequences; the arrays show a peak at 250 bp (size of the original fragment, containing 3 repeats), with progressively smaller peaks at 320 bp (corresponding to one new acquisition) and 390 bp (2 new acquisitions). The raw reads were then mapped back to the clusters, to create a table presenting the abundance of each cluster in each biological repeat.

### Spacer extraction

The centroid sequence of each cluster was used for extraction of acquired new spacer sequences. To identify true acquisition events while excluding DNA rearrangement events (which may also result in an extended PCR product), a custom-made R script based on the ‘Biostrings’ and ‘tools’ R packages was used to count the number of repeats in each centroid sequence, allowing up to 2 mismatches per repeat. Fragments containing more than 3 repeats were tagged as putative acquisitions, and the sequence between the 2 repeats closest to the leader end was extracted. “False positives”, which are the result of rearrangement of spacers within or between arrays rather than canonical acquisition, were eliminated by screening the extracted putative spacers against all original CRISPR arrays, allowing up to 5 mismatches per spacer; spacers with matches in existing CRISPR arrays were excluded from further analysis.

### Mapping new spacers to genomic location

In order to establish the protospacer location for each new acquisition, we used the blastn-short program 15 at an E-value of .001 against a reference database containing *H. volcanii* and 48N, including their natural plasmids and any additional plasmid present in the experiment (pWL102 in the mating experiment). Blast results were refined using custom made-R scripts based on the ‘stringr’ package. In brief, no more than 2 mismatches between the spacer and the protospacer were allowed; in cases of multiple matches, the location with the highest score was preserved (while conserving abundance information). Furthermore, spacers that were aligned only partially were removed when the alignment length was shorter than the total spacer length by three or more bases.

### Generating ⍰pyrE2 48N strain

Strain 48N was verified for its sensitivity to 5FOA (150 μg/ml) in YPC medium and its ability to grow on CA medium. Knock out of the pyrE2 gene was done with pGB68 ^35^ by pop-in and then pop-out as described in the reference. Pop-in transformants were selected on YPC plates containing Novobiocin (0.3µg/ml) and individual colonies were then transferred into liquid YPC medium with no novobiocin and shaken in 45°C for 48 hours after which they were streaked on YPC plates supplemented with 5FOA. Resistant colonies were picked and streaked on CA plates and YPC plates. Notably, half of the colonies retained their ability to grow on CA whileL resistant to 5FOA. To obtain true knockout colonies, 3 cycles of 5FOA selection were performed. The genotype of the pop-in colonies, as well as knockout colonies was confirmed by PCR using the primers:IT748, IT749.

### Virus-cured 48N strain construction

Construction of the 48 cured strain was performed according to the pop-in/pop-out protocol described in (Allers et al. 2004; ^35^. In this method, the upstream and downstream flanking regions of the sequence to be deleted are amplified by PCR and cloned together into the ‘suicide plasmid’ pTA131, that cannot replicate autonomously and carries the *pyrE2* selectable genetic marker. The plasmids are then transformed (Transformation of 48N was carried out using the polyethylene glycol 600 method as described in (Allers et al. 2004; Cline et al. 1989) into the 48N Δ*pyrE2* mutant, and the transformants, in which the plasmids have been integrated into the chromosome, are selected for on plates that lack uracil (’pop-in’). Upon counter-selection on plates containing uracil and 5-fluoroorotic acid (5-FOA), the only cells that survive are those in which the integrated plasmids have been excised by spontaneous intra-chromosomal homologues recombination (’pop-out’), either restoring the WT gene or resulting in allele exchange. The generation of the LSV-48N-cured strain was performed in two steps. We first deleted the viral integrase gene, so that viral genomes present in the circular form will not be able to re-integrate after the deletion has occurred. We used the Δ-integrase strain UG595 and inserted it with pIS76 plasmid to delete the entire virus from the chromosome. The ‘pop-out’ strains were screened using pairs of primers IT1 and IT4 located upstream and downstream to the LSV-48N integration site and the sequence was validated by sequencing.

### Preprocessing of Illumina and PacBio sequencing data

The Illumina raw reads used for the previous, publicly available, assembly of *Haloferax volcanii* spp. 48N were fetched. Reads mapping to PhiX174 (a common library preparation spike-in) were identified using bowtie2 ^36^. Subsequently, unmapped reads were filtered using Trimmomatic (version 0.36) (Bolger et al. 2014) using the command ‘adapters/NexteraPE-PE.fa:2:30:10 LEADING:3 TRAILING:3 SLIDINGWINDOW:4:15 MINLEN:36‘. PacificBio Science (“long”) reads were filtered using bbduk.sh from the bbtools suite (Bushnell, 2014) with the following flags: minlen=100 maq=8, i.e. reads were retained only if they were of a minimal length of 100 nucleotides and had a minimal average quality of 8.

### Assembly of 48N and LSV-48N genomes

LSV-48N processed reads were assembled using SPAdes (version 3.11.0). Manual examination of the assembly graph, together with PCR amplification, were used to verify the genome terminals and the circular topology.

The hybrid assembly of Haloferax volcanii spp. 48N was performed using Unicycler (version 0.4.4, Wick et al. 2017) with default options, supplying pilon for genome polishing (Walker et al. 2014). The assembly graph was examined and the topology of the virus form was manually resolved. Mapping of the raw reads and manual examination of the coverage suggested both integrated and circular forms of the virus were present in the original culture. For simplicity, the resolved form used for coverage (depth) calculation was set as the circular form (i.e. as a stand alone replicon as a separate contig) in order to report the virus coverage separately from that of the host chromosome.

Coverage reports (including estimated depth) where generated using bbmap.sh from the bbtools suite (Bushnell, 2014), via the ‘covstats’ output. For these estimation, only the Illumina reads were provided as a query and the resolved 48N genome as reference.

Finally, the hybrid assembly of *Haloferax volcanii* spp. 48N completeness was estimated at 99.57% with contamination of 0.13% (CheckM version 1.0.18, ^37^).

### Genome structural and functional annotations

For gene prediction, the assembled genomes were analyzed using genemark ^38^. Predicted protein sequences were then searched (using hmmsearch ^39,40^, mmseqs ^41^ and psi-BLAST ^42^) against several public protein domain databases (Pfam, PDB, CDD and arCOGs^43^). The functional/descriptive label of profile matches with good alignment statistics was cloned onto the query protein. Predicted proteins without sufficiently reliable match were manually examined further using web-hhpred ^44^and CollabFold2 ^45^.

### Live cell imaging

The experimental setup utilized a microfluidics chip, crafted from polydimethylsiloxane (SylgardTM 184 Silicone Elastomer Kit, Dowsil), along with its curing agent. This chip was cast on a SU-8 mold (designed by Klayout, F-Si) with the following specifications: Eight parallel flow channels (200 μm wide, 22 μm deep), divided into two smaller channels (100 μm wide, 22 μm deep). The latter comprised 144 equally spaced chambers, featuring varying dimensions (10, 20, 40, 60, 80, 100 μm wide, 60 μm deep, 0.83 μm height) on their lateral sides. To secure the microfluidics chip, a round cover glass (diameter = 47 mm, # 1.5; Menzel-GlaLJser, Braunschweig, Germany) was employed and affixed using an oxygen plasma cleaner (PDC-32G-2 Plasma Cleaner, Harrick Plasma, New York, USA). The archaea cultivated within this microfluidic system underwent observation and monitoring through a Nikon Ti inverted microscope equipped with a PFS (Perfect Focus System), an Andor clara CCD camera (Andor Tech., Belfast, UK), and an OKOlab Microscope Incubation System (OKOlabs GmbH, Italy). The microscope setup included a motorized stage, automated stage controllers (Nikon), shutters (BD, USA), and a Plan Apochromat Lambda D 100X Oil immersion objective (Nikon), all controlled by the NIS Elements software.

The experimental protocol involved growing virus-infected 48N and virus-cured 48N, in YPC medium until they reached the stationary phase. Subsequently, each strain was individually introduced into a separate main channel and allowed to grow in YPC medium at 37°C for a duration of 20 hours. Following this incubation period, chambers containing these strains were selected, and a time lapse of 19.5 hours was recorded at 100X magnification, with a timer interval of 5 minutes between each frame. Upon completion of the experiment, data were collected and analyzed using Python scripts.

### RNA-Seq

Total RNA was extracted from 4 replicates of virus-cured 48N, and 3 replicates of wt 48N, using the RNeasy kit (Qiagen). Extracted RNA was shipped in dry ice to the DNA Services facility within the Research Resources Center (RRC), University of Illinois at Chicago.

Libraries were prepared with NuGen’s Universal Prokaryotic kit, including a rRNA depletion step, and sequenced (2×150bp) on a NovaSeq6000 SP. Raw sequencing data was processed using BBtools tool suite (sourceforge.net/projects/bbmap/,our full pipeline uploaded to https://github.com/UriNeri/LSV). In brief, bbduk was used for quality filtration, adaptor trimming, removal of known sequencing artifacts such as phiX, and exclusion of reads mapping to the rRNA operon, which could not be entirely eliminated at the rRNA depletion step. BBtools sequence expression analyzer ‘seal’ was then used to map cleaned data to 48N coding sequences and calculate RPKM values. A single outlier replicate of virus-cured 48N was discarded at this stage, leaving 3 wt vs. 3 virus-cured replicates for differential gene expression analysis using R package EdgeR. Graphical representations were generated utilizing the Bioinfokit and Matplotlib packages in Python.

### Data and code availability

Complete assemblies for Haloferax 48N sp. Atlit were deposited to NCBI genomes, accession numbers CP137689-CP137693; this is a revision of the previous draft assembly, deposited in NCBI (PRJNA431124). Complete sequences of the virus LSV-48N and the plasmid pWL102 are available from GenBank, accession numbers PP151251 and PP297526, respectively. Raw sequence data from RNASeq, and from CRISPR spacer acquisition assays, are found in NCBIs SRA (Sequence Read Archive), project accession numbers PRJNA1072013 and PRJNA1072009, respectively. Custom code and scripts generated or used in this work are fully and freely available under the MIT open source license via https://github.com/UriNeri/HLSV1

