## Supplemental Table 1 for "A new archaeal virus that suppresses the transcription of host immunity genes"

**A.** **Number of unique spacer matches**

| | 48N chromosome | pWL-102 | *H. volcanii* chromosome | phv1 | phv3 | phv4 | p48N_2 | Multiple hits |
|---|---|---|---|---|---|---|---|---|
| Replicate 1 | 47 | 5 | 18 | 2 | 8 | 6 | 1 | 78 |
| Replicate 2 | 178 | 18 | 44 | 11 | 27 | 26 | 4 | 206 |
| Replicate 3 | 201 | 14 | 25 | 1 | 25 | 10 | 2 | 128 |
| Replicate 4 | 276 | 30 | 56 | 5 | 42 | 21 | 2 | 247 |
| Replicate 5 | 107 | 5 | 34 | 1 | 13 | 14 | 4 | 92 |

**B.** **Total number of unique spacer matches**

| | 48N chromosome | pWL-102 | *H. volcanii* chromosome | phv1 | phv3 | phv4 | p48N_2 | Multiple hits |
|---|---|---|---|---|---|---|---|---|
| Replicate 1 | 406 | 5 | 18 | 2 | 8 | 6 | 1 | 82 |
| Replicate 2 | 6067 | 18 | 45 | 11 | 27 | 26 | 4 | 229 |
| Replicate 3 | 7303 | 14 | 26 | 1 | 26 | 10 | 2 | 157 |
| Replicate 4 | 6110 | 32 | 58 | 5 | 45 | 21 | 2 | 542 |
| Replicate 5 | 1119 | 8 | 36 | 1 | 13 | 14 | 5 | 250 |
