## Supplemental Table 2 for "A new archaeal virus that suppresses the transcription of host immunity genes"

**Supplementary Table T2 – Sequence coverage statistics of 48N supernatant,  entries corresponding to the LSV-48N genome are highlighted by green background.**

| #ID | Avg_fold | Length | Ref_GC | Covered_percent | Covered_bases | Plus_reads | Minus_reads | Read_GC | Median_fold |
|---|---|---|---|---|---|---|---|---|---|
| NODE_1_length_24864_cov_2599.177754 | 7403.051 | 24864 | 0.6059 | 100.00 | 24864 | 602812 | 603150 | 0.6062 | 7690 |
| NODE_2_length_16315_cov_6.742949 | 17.2951 | 16315 | 0.5479 | 100.00 | 16315 | 984 | 983 | 0.5576 | 17 |
| NODE_3_length_1714_cov_1.740379 | 4.8279 | 1714 | 0.6831 | 99.53 | 1706 | 30 | 28 | 0.6837 | 5 |
| NODE_4_length_1356_cov_1.606724 | 4.6342 | 1356 | 0.6336 | 96.61 | 1310 | 21 | 23 | 0.6311 | 5 |
| NODE_5_length_1281_cov_1.758306 | 4.4887 | 1281 | 0.662 | 100.00 | 1281 | 20 | 21 | 0.6588 | 4 |
| NODE_6_length_1280_cov_1.158770 | 2.5852 | 1280 | 0.7226 | 97.73 | 1251 | 10 | 12 | 0.7196 | 2 |
| NODE_7_length_1272_cov_7.209205 | 16.6399 | 1272 | 0.6148 | 97.17 | 1236 | 69 | 75 | 0.6124 | 18 |
| NODE_8_length_941_cov_1.671296 | 4.2869 | 941 | 0.7216 | 99.04 | 932 | 15 | 12 | 0.713 | 4 |
| NODE_9_length_866_cov_1.468948 | 3.6859 | 866 | 0.6986 | 100.00 | 866 | 12 | 9 | 0.7132 | 4 |
| NODE_10_length_714_cov_1.053375 | 2.2003 | 714 | 0.7311 | 100.00 | 714 | 7 | 4 | 0.7407 | 2 |
| NODE_11_length_704_cov_1.551834 | 4.2031 | 704 | 0.6861 | 100.00 | 704 | 10 | 10 | 0.6929 | 4 |
| NODE_12_length_694_cov_0.771475 | 1.7305 | 694 | 0.6873 | 100.00 | 694 | 4 | 4 | 0.6886 | 2 |
| NODE_13_length_650_cov_1.670157 | 3.2262 | 650 | 0.6662 | 100.00 | 650 | 7 | 7 | 0.6581 | 3 |
| NODE_14_length_646_cov_0.980668 | 2.1409 | 646 | 0.6687 | 100.00 | 646 | 5 | 5 | 0.6616 | 2 |
| NODE_15_length_621_cov_0.926471 | 1.8357 | 621 | 0.7053 | 100.00 | 621 | 4 | 4 | 0.7088 | 2 |
| NODE_16_length_616_cov_0.988868 | 1.9529 | 616 | 0.668 | 83.93 | 517 | 4 | 4 | 0.6719 | 2 |
| NODE_17_length_613_cov_0.910448 | 1.9086 | 613 | 0.6395 | 100.00 | 613 | 4 | 4 | 0.6156 | 2 |
| NODE_18_length_605_cov_0.875000 | 2.0248 | 605 | 0.6777 | 100.00 | 605 | 5 | 4 | 0.6955 | 2 |
| NODE_19_length_603_cov_1.127376 | 2.4892 | 603 | 0.6683 | 100.00 | 603 | 6 | 4 | 0.6396 | 2 |
| NODE_20_length_592_cov_1.034951 | 1.9747 | 592 | 0.7331 | 100.00 | 592 | 3 | 5 | 0.7399 | 2 |
| NODE_21_length_591_cov_0.655642 | 2.0372 | 591 | 0.6277 | 100.00 | 591 | 3 | 5 | 0.6279 | 2 |
| NODE_22_length_564_cov_1.509240 | 3.1259 | 564 | 0.6489 | 100.00 | 564 | 6 | 6 | 0.6469 | 3 |
| NODE_23_length_560_cov_0.863354 | 1.6089 | 560 | 0.66 | 98.21 | 550 | 3 | 3 | 0.657 | 2 |
| NODE_24_length_548_cov_1.095541 | 2.1551 | 548 | 0.6971 | 100.00 | 548 | 4 | 4 | 0.6913 | 2 |
| NODE_25_length_525_cov_0.767857 | 1.4343 | 525 | 0.7124 | 100.00 | 525 | 2 | 3 | 0.7158 | 1 |
| NODE_26_length_520_cov_0.697517 | 1.4404 | 520 | 0.5577 | 100.00 | 520 | 3 | 2 | 0.5607 | 1 |
| NODE_27_length_510_cov_1.214781 | 2.4882 | 510 | 0.6706 | 100.00 | 510 | 4 | 5 | 0.6675 | 2 |
| NODE_28_length_508_cov_0.721578 | 1.5217 | 508 | 0.6732 | 100.00 | 508 | 3 | 3 | 0.6843 | 1 |
| NODE_29_length_503_cov_1.600939 | 3.2863 | 503 | 0.6998 | 100.00 | 503 | 6 | 5 | 0.7078 | 3 |
| NODE_30_length_496_cov_0.737470 | 1.7681 | 496 | 0.7661 | 100.00 | 496 | 4 | 3 | 0.7685 | 2 |
| NODE_31_length_495_cov_1.081340 | 2.6101 | 495 | 0.6909 | 100.00 | 495 | 4 | 5 | 0.6888 | 3 |
| NODE_32_length_495_cov_0.937799 | 1.8222 | 495 | 0.5697 | 100.00 | 495 | 3 | 3 | 0.5698 | 2 |
| NODE_33_length_494_cov_0.829736 | 1.913 | 494 | 0.6579 | 100.00 | 494 | 3 | 4 | 0.6741 | 1 |
| NODE_34_length_492_cov_1.192771 | 2.3171 | 492 | 0.628 | 100.00 | 492 | 4 | 4 | 0.6219 | 2 |
| NODE_35_length_489_cov_0.983010 | 2.1104 | 489 | 0.7014 | 100.00 | 489 | 4 | 3 | 0.7074 | 2 |
| NODE_36_length_488_cov_0.871046 | 1.5389 | 488 | 0.7316 | 100.00 | 488 | 3 | 2 | 0.7324 | 2 |
| NODE_37_length_485_cov_0.899510 | 1.5505 | 485 | 0.6577 | 100.00 | 485 | 3 | 2 | 0.6596 | 1 |

| | | | | | | | | | |
|---|---|---|---|---|---|---|---|---|---|
| NODE_38_length_477_cov_1.447500 | 2.8302 | 477 | 0.6562 | 100.00 | 477 | 4 | 5 | 0.6585 | 3 |
| NODE_39_length_477_cov_0.635000 | 1.5765 | 477 | 0.6834 | 100.00 | 477 | 3 | 2 | 0.6809 | 1 |
| NODE_40_length_475_cov_0.670854 | 1.2674 | 475 | 0.6316 | 100.00 | 475 | 2 | 2 | 0.6379 | 1 |
| NODE_41_length_470_cov_0.674300 | 1.2809 | 470 | 0.7426 | 100.00 | 470 | 2 | 2 | 0.7409 | 1 |
| NODE_42_length_469_cov_0.742347 | 1.597 | 469 | 0.6823 | 100.00 | 469 | 2 | 3 | 0.6889 | 1 |
| NODE_43_length_465_cov_0.976804 | 3.0946 | 465 | 0.5226 | 100.00 | 465 | 5 | 6 | 0.516 | 3 |
| NODE_44_length_454_cov_0.960212 | 1.9846 | 454 | 0.5617 | 100.00 | 454 | 3 | 3 | 0.5538 | 2 |
| NODE_45_length_447_cov_0.764865 | 1.9172 | 447 | 0.6264 | 100.00 | 447 | 3 | 3 | 0.6112 | 2 |
| NODE_46_length_441_cov_1.013736 | 1.7098 | 441 | 0.6349 | 100.00 | 441 | 3 | 2 | 0.6393 | 2 |
| NODE_47_length_440_cov_0.966942 | 1.8705 | 440 | 0.5864 | 100.00 | 440 | 3 | 4 | 0.5735 | 2 |
| NODE_48_length_436_cov_1.222841 | 2.0711 | 436 | 0.6858 | 100.00 | 436 | 3 | 3 | 0.6955 | 2 |
| NODE_49_length_435_cov_1.773743 | 3.1126 | 435 | 0.7195 | 100.00 | 435 | 4 | 5 | 0.7142 | 3 |
| NODE_50_length_435_cov_0.818436 | 1.731 | 435 | 0.6943 | 100.00 | 435 | 2 | 3 | 0.6972 | 2 |
| NODE_51_length_434_cov_1.126050 | 2.0783 | 434 | 0.659 | 100.00 | 434 | 3 | 3 | 0.6541 | 2 |
| NODE_52_length_430_cov_1.220963 | 2.6512 | 430 | 0.5674 | 100.00 | 430 | 4 | 4 | 0.5763 | 2 |
| NODE_53_length_427_cov_0.765714 | 1.4052 | 427 | 0.6183 | 100.00 | 427 | 2 | 2 | 0.6183 | 1 |
| NODE_54_length_425_cov_1.747126 | 3.5412 | 425 | 0.6706 | 100.00 | 425 | 5 | 5 | 0.687 | 3 |
| NODE_55_length_425_cov_1.051724 | 2.0447 | 425 | 0.6518 | 100.00 | 425 | 2 | 4 | 0.6733 | 2 |
| NODE_56_length_420_cov_0.825073 | 1.4262 | 420 | 0.7143 | 100.00 | 420 | 2 | 2 | 0.7195 | 1 |
| NODE_57_length_418_cov_0.542522 | 1.3923 | 418 | 0.6675 | 100.00 | 418 | 2 | 2 | 0.6817 | 1 |
| NODE_58_length_415_cov_0.863905 | 1.4482 | 415 | 0.6434 | 100.00 | 415 | 2 | 2 | 0.6456 | 1 |
| NODE_59_length_414_cov_0.982196 | 2.1691 | 414 | 0.6039 | 100.00 | 414 | 3 | 3 | 0.6047 | 2 |
| NODE_60_length_412_cov_1.638806 | 3.284 | 412 | 0.6917 | 91.75 | 378 | 6 | 3 | 0.6807 | 4 |
| NODE_61_length_412_cov_1.576119 | 3.2985 | 412 | 0.716 | 100.00 | 412 | 6 | 4 | 0.7079 | 4 |
| NODE_62_length_410_cov_1.855856 | 3.5707 | 410 | 0.5878 | 100.00 | 410 | 5 | 6 | 0.5563 | 4 |
| NODE_63_length_409_cov_1.307229 | 2.934 | 409 | 0.5721 | 100.00 | 409 | 4 | 4 | 0.58 | 3 |
| NODE_64_length_407_cov_0.881818 | 1.4767 | 407 | 0.6364 | 100.00 | 407 | 2 | 2 | 0.629 | 1 |
| NODE_65_length_405_cov_1.954268 | 4.2765 | 405 | 0.5605 | 100.00 | 405 | 5 | 7 | 0.5486 | 4 |
| NODE_66_length_405_cov_1.381098 | 3.4123 | 405 | 0.7556 | 100.00 | 405 | 6 | 4 | 0.7489 | 3 |
| NODE_67_length_402_cov_1.335385 | 2.2338 | 402 | 0.6791 | 100.00 | 402 | 3 | 3 | 0.6793 | 2 |
| NODE_68_length_401_cov_0.953704 | 1.7382 | 401 | 0.6409 | 100.00 | 401 | 2 | 3 | 0.6414 | 2 |
| NODE_69_length_401_cov_0.743827 | 1.9451 | 401 | 0.7282 | 100.00 | 401 | 3 | 3 | 0.724 | 2 |
| NODE_70_length_401_cov_0.731481 | 2.6135 | 401 | 0.5586 | 100.00 | 401 | 4 | 3 | 0.5624 | 3 |
| NODE_71_length_400_cov_1.349845 | 2.2475 | 400 | 0.595 | 100.00 | 400 | 3 | 3 | 0.5818 | 2 |
| NODE_72_length_398_cov_1.146417 | 1.892 | 398 | 0.7136 | 100.00 | 398 | 2 | 3 | 0.7251 | 2 |
| NODE_73_length_398_cov_0.688474 | 1.1357 | 398 | 0.7412 | 100.00 | 398 | 1 | 2 | 0.7588 | 1 |
| NODE_74_length_394_cov_1.605678 | 3.2538 | 394 | 0.6497 | 100.00 | 394 | 5 | 5 | 0.6763 | 2 |
| NODE_75_length_394_cov_1.186120 | 2.2893 | 394 | 0.6218 | 100.00 | 394 | 3 | 3 | 0.6308 | 2 |
| NODE_76_length_394_cov_0.911672 | 1.8883 | 394 | 0.6142 | 100.00 | 394 | 3 | 2 | 0.6282 | 2 |

| | | | | | | | | | |
|---|---|---|---|---|---|---|---|---|---|
| NODE_77_length_394_cov_0.861199 | 1.9391 | 394 | 0.6878 | 100.00 | 394 | 2 | 4 | 0.7094 | 1 |
| NODE_78_length_393_cov_0.930380 | 1.5318 | 393 | 0.6005 | 100.00 | 393 | 2 | 2 | 0.5997 | 1 |
| NODE_79_length_392_cov_1.469841 | 2.5587 | 392 | 0.4949 | 99.23 | 389 | 4 | 3 | 0.4865 | 1 |
| NODE_80_length_392_cov_1.387302 | 3.0893 | 392 | 0.6199 | 100.00 | 392 | 4 | 4 | 0.606 | 3 |
| NODE_81_length_391_cov_0.990446 | 2.3043 | 391 | 0.6573 | 100.00 | 391 | 3 | 3 | 0.6593 | 3 |
| NODE_82_length_390_cov_0.939297 | 1.5436 | 390 | 0.6846 | 100.00 | 390 | 1 | 3 | 0.6744 | 1 |
| NODE_83_length_389_cov_0.708333 | 1.162 | 389 | 0.6401 | 100.00 | 389 | 2 | 1 | 0.6239 | 1 |
| NODE_84_length_384_cov_0.866450 | 1.5651 | 384 | 0.6276 | 100.00 | 384 | 2 | 2 | 0.6306 | 2 |
| NODE_85_length_382_cov_1.409836 | 2.356 | 382 | 0.6597 | 100.00 | 382 | 3 | 3 | 0.66 | 2 |
| NODE_86_length_382_cov_1.055738 | 2.5628 | 382 | 0.6623 | 100.00 | 382 | 4 | 5 | 0.665 | 2 |
| NODE_87_length_381_cov_0.861842 | 1.5774 | 381 | 0.6693 | 100.00 | 381 | 2 | 2 | 0.6589 | 1 |
| NODE_88_length_380_cov_0.966997 | 1.5816 | 380 | 0.6474 | 100.00 | 380 | 2 | 2 | 0.6456 | 1 |
| NODE_89_length_380_cov_0.858086 | 1.5842 | 380 | 0.6816 | 100.00 | 380 | 2 | 2 | 0.6927 | 1 |
| NODE_90_length_379_cov_1.370861 | 2.6517 | 379 | 0.6939 | 100.00 | 379 | 4 | 3 | 0.6985 | 3 |
| NODE_91_length_378_cov_0.677741 | 1.1931 | 378 | 0.6825 | 100.00 | 378 | 1 | 2 | 0.6608 | 1 |
| NODE_92_length_376_cov_1.722408 | 2.8032 | 376 | 0.734 | 100.00 | 376 | 3 | 4 | 0.7287 | 2 |
| NODE_93_length_376_cov_1.451505 | 2.3989 | 376 | 0.7074 | 100.00 | 376 | 2 | 4 | 0.7065 | 2 |
| NODE_94_length_376_cov_0.909699 | 1.7713 | 376 | 0.7074 | 100.00 | 376 | 2 | 3 | 0.7057 | 2 |
| NODE_95_length_376_cov_0.846154 | 1.6011 | 376 | 0.6941 | 98.94 | 372 | 2 | 2 | 0.6977 | 1 |
| NODE_96_length_375_cov_1.221477 | 2 | 375 | 0.704 | 100.00 | 375 | 3 | 2 | 0.7 | 2 |
| NODE_97_length_374_cov_1.020202 | 2.0053 | 374 | 0.7005 | 100.00 | 374 | 4 | 1 | 0.7133 | 2 |
| NODE_98_length_373_cov_1.344595 | 2.6247 | 373 | 0.6568 | 100.00 | 373 | 4 | 3 | 0.6394 | 3 |
| NODE_99_length_373_cov_1.216216 | 2.0241 | 373 | 0.7078 | 100.00 | 373 | 2 | 3 | 0.698 | 2 |
| NODE_100_length_372_cov_0.738983 | 1.207 | 372 | 0.6935 | 100.00 | 372 | 2 | 1 | 0.6889 | 1 |
| NODE_101_length_370_cov_0.993174 | 1.6189 | 370 | 0.7324 | 100.00 | 370 | 2 | 2 | 0.7412 | 2 |
| NODE_102_length_370_cov_0.631399 | 1.2162 | 370 | 0.7081 | 100.00 | 370 | 2 | 1 | 0.7111 | 1 |
| NODE_103_length_369_cov_0.753425 | 2.0407 | 369 | 0.7398 | 100.00 | 369 | 3 | 2 | 0.7387 | 2 |
| NODE_104_length_368_cov_0.862543 | 1.6332 | 368 | 0.6793 | 100.00 | 368 | 1 | 3 | 0.6905 | 2 |
| NODE_105_length_368_cov_0.756014 | 1.2255 | 368 | 0.6793 | 100.00 | 368 | 2 | 1 | 0.694 | 1 |
| NODE_106_length_368_cov_0.666667 | 1.8505 | 368 | 0.7011 | 100.00 | 368 | 3 | 2 | 0.6981 | 1 |
| NODE_107_length_367_cov_0.734483 | 1.2262 | 367 | 0.673 | 100.00 | 367 | 2 | 1 | 0.6711 | 1 |
| NODE_108_length_366_cov_0.892734 | 1.9262 | 366 | 0.6093 | 100.00 | 366 | 3 | 3 | 0.6043 | 2 |
| NODE_109_length_366_cov_0.813149 | 1.6448 | 366 | 0.6667 | 100.00 | 366 | 1 | 3 | 0.6661 | 1 |
| NODE_110_length_366_cov_0.726644 | 1.235 | 366 | 0.7077 | 100.00 | 366 | 2 | 1 | 0.7168 | 1 |
| NODE_111_length_364_cov_0.759582 | 1.6511 | 364 | 0.5962 | 100.00 | 364 | 2 | 2 | 0.5907 | 2 |
| NODE_112_length_364_cov_0.742160 | 1.2418 | 364 | 0.5934 | 100.00 | 364 | 2 | 1 | 0.5841 | 1 |
| NODE_113_length_363_cov_1.013986 | 1.6501 | 363 | 0.6804 | 100.00 | 363 | 2 | 2 | 0.6912 | 1 |
| NODE_114_length_363_cov_0.807692 | 2.0606 | 363 | 0.6639 | 100.00 | 363 | 3 | 2 | 0.6484 | 2 |
| NODE_115_length_361_cov_1.605634 | 3.3435 | 361 | 0.615 | 100.00 | 361 | 5 | 3 | 0.6181 | 3 |

| | | | | | | | | | |
|---|---|---|---|---|---|---|---|---|---|
| NODE_116_length_360_cov_1.162544 | 2.5028 | 360 | 0.6861 | 100.00 | 360 | 1 | 5 | 0.6704 | 3 |
| NODE_117_length_360_cov_0.780919 | 1.2556 | 360 | 0.6917 | 100.00 | 360 | 2 | 1 | 0.6991 | 1 |
| NODE_118_length_358_cov_1.206406 | 2.0978 | 358 | 0.6453 | 100.00 | 358 | 3 | 2 | 0.6418 | 2 |
| NODE_119_length_355_cov_1.575540 | 2.5408 | 355 | 0.6592 | 100.00 | 355 | 3 | 3 | 0.6796 | 1 |
| NODE_120_length_355_cov_0.967626 | 2.5042 | 355 | 0.6225 | 100.00 | 355 | 3 | 3 | 0.5969 | 2 |
| NODE_121_length_352_cov_0.923636 | 1.7074 | 352 | 0.7244 | 100.00 | 352 | 2 | 2 | 0.7238 | 2 |
| NODE_122_length_352_cov_0.843636 | 1.7102 | 352 | 0.6534 | 100.00 | 352 | 2 | 2 | 0.6661 | 1 |
| NODE_123_length_350_cov_2.076923 | 3.4429 | 350 | 0.6829 | 100.00 | 350 | 4 | 4 | 0.6722 | 3 |
| NODE_124_length_350_cov_1.446886 | 2.5771 | 350 | 0.6629 | 100.00 | 350 | 4 | 2 | 0.6552 | 3 |
| NODE_125_length_350_cov_1.175824 | 2.1571 | 350 | 0.6829 | 100.00 | 350 | 3 | 2 | 0.6887 | 2 |
| NODE_126_length_350_cov_0.805861 | 1.2886 | 350 | 0.6914 | 100.00 | 350 | 2 | 1 | 0.694 | 1 |
| NODE_127_length_349_cov_0.661765 | 1.6705 | 349 | 0.6504 | 100.00 | 349 | 1 | 3 | 0.6561 | 2 |
| NODE_128_length_348_cov_0.929889 | 1.727 | 348 | 0.7155 | 100.00 | 348 | 2 | 2 | 0.7221 | 1 |
| NODE_129_length_347_cov_1.059259 | 1.7291 | 347 | 0.4813 | 100.00 | 347 | 2 | 2 | 0.4833 | 2 |
| NODE_130_length_347_cov_0.988889 | 1.7291 | 347 | 0.6167 | 100.00 | 347 | 2 | 2 | 0.615 | 2 |
| NODE_131_length_347_cov_0.833333 | 2.245 | 347 | 0.536 | 100.00 | 347 | 3 | 3 | 0.5366 | 2 |
| NODE_132_length_346_cov_2.167286 | 3.474 | 346 | 0.578 | 100.00 | 346 | 5 | 3 | 0.6007 | 3 |
| NODE_133_length_346_cov_0.684015 | 1.1994 | 346 | 0.685 | 100.00 | 346 | 2 | 1 | 0.6578 | 1 |
| NODE_134_length_345_cov_1.563433 | 2.6145 | 345 | 0.658 | 100.00 | 345 | 3 | 3 | 0.6574 | 2 |
| NODE_135_length_345_cov_1.272388 | 2.4116 | 345 | 0.5565 | 100.00 | 345 | 3 | 3 | 0.5493 | 3 |
| NODE_136_length_345_cov_1.033582 | 2.1797 | 345 | 0.5681 | 100.00 | 345 | 2 | 3 | 0.5705 | 2 |
| NODE_137_length_345_cov_0.865672 | 1.6 | 345 | 0.658 | 100.00 | 345 | 2 | 2 | 0.6558 | 1 |
| NODE_138_length_344_cov_0.846442 | 1.7355 | 344 | 0.7355 | 100.00 | 344 | 2 | 2 | 0.7353 | 1 |
| NODE_139_length_344_cov_0.805243 | 1.7326 | 344 | 0.7209 | 100.00 | 344 | 2 | 2 | 0.7102 | 2 |
| NODE_140_length_342_cov_1.098113 | 1.7515 | 342 | 0.652 | 100.00 | 342 | 2 | 2 | 0.6483 | 2 |
| NODE_141_length_341_cov_0.837121 | 1.3255 | 341 | 0.6217 | 100.00 | 341 | 2 | 1 | 0.6128 | 1 |
| NODE_142_length_341_cov_0.837121 | 1.3255 | 341 | 0.6481 | 100.00 | 341 | 2 | 1 | 0.6527 | 1 |
| NODE_143_length_340_cov_0.688213 | 1.3265 | 340 | 0.7029 | 100.00 | 340 | 1 | 2 | 0.7051 | 1 |
| NODE_144_length_339_cov_0.671756 | 2.1652 | 339 | 0.6903 | 100.00 | 339 | 2 | 3 | 0.6853 | 2 |
| NODE_145_length_338_cov_0.842912 | 1.3343 | 338 | 0.7101 | 100.00 | 338 | 2 | 1 | 0.7051 | 1 |
| NODE_146_length_337_cov_1.111538 | 3.0148 | 337 | 0.4955 | 100.00 | 337 | 6 | 1 | 0.5234 | 3 |
| NODE_147_length_336_cov_1.606178 | 3.1012 | 336 | 0.5685 | 100.00 | 336 | 3 | 4 | 0.5751 | 3 |
| NODE_148_length_336_cov_0.938224 | 1.7917 | 336 | 0.6548 | 100.00 | 336 | 2 | 2 | 0.6545 | 2 |
| NODE_149_length_336_cov_0.907336 | 1.7887 | 336 | 0.7143 | 100.00 | 336 | 2 | 2 | 0.7238 | 2 |
| NODE_150_length_336_cov_0.845560 | 1.6845 | 336 | 0.6935 | 100.00 | 336 | 2 | 2 | 0.6766 | 2 |
| NODE_151_length_335_cov_1.670543 | 2.6716 | 335 | 0.5224 | 100.00 | 335 | 3 | 3 | 0.5084 | 3 |
| NODE_152_length_335_cov_1.131783 | 1.791 | 335 | 0.6657 | 100.00 | 335 | 1 | 3 | 0.6683 | 2 |
| NODE_153_length_335_cov_0.848837 | 1.3433 | 335 | 0.6627 | 100.00 | 335 | 2 | 1 | 0.6644 | 1 |
| NODE_154_length_335_cov_0.848837 | 1.3433 | 335 | 0.6537 | 100.00 | 335 | 2 | 1 | 0.6444 | 1 |

| | | | | | | | | | |
|---|---|---|---|---|---|---|---|---|---|
| NODE_155_length_333_cov_1.062500 | 2.8228 | 333 | 0.7538 | 100.00 | 333 | 4 | 4 | 0.7463 | 2 |
| NODE_156_length_333_cov_0.851562 | 2.1411 | 333 | 0.7207 | 100.00 | 333 | 3 | 2 | 0.7093 | 2 |
| NODE_157_length_333_cov_0.675781 | 1.4444 | 333 | 0.6757 | 100.00 | 333 | 2 | 2 | 0.6694 | 1 |
| NODE_158_length_332_cov_1.435294 | 2.2651 | 332 | 0.6205 | 100.00 | 332 | 2 | 3 | 0.6303 | 2 |
| NODE_159_length_331_cov_0.858268 | 1.3595 | 331 | 0.4804 | 100.00 | 331 | 1 | 2 | 0.4956 | 1 |
| NODE_160_length_330_cov_0.632411 | 1.7636 | 330 | 0.6636 | 100.00 | 330 | 0 | 4 | 0.6611 | 2 |
| NODE_161_length_329_cov_1.154762 | 1.8237 | 329 | 0.6687 | 100.00 | 329 | 2 | 2 | 0.655 | 2 |
| NODE_162_length_329_cov_0.972222 | 1.6839 | 329 | 0.6474 | 100.00 | 329 | 1 | 3 | 0.6625 | 2 |
| NODE_163_length_329_cov_0.861111 | 1.3617 | 329 | 0.696 | 100.00 | 329 | 2 | 1 | 0.6896 | 1 |
| NODE_164_length_328_cov_1.167331 | 2.189 | 328 | 0.5122 | 100.00 | 328 | 2 | 3 | 0.5167 | 2 |
| NODE_165_length_328_cov_1.155378 | 2.689 | 328 | 0.6707 | 100.00 | 328 | 3 | 3 | 0.656 | 3 |
| NODE_166_length_328_cov_0.880478 | 2.125 | 328 | 0.5427 | 100.00 | 328 | 2 | 4 | 0.5366 | 2 |
| NODE_167_length_328_cov_0.840637 | 2.0091 | 328 | 0.6372 | 100.00 | 328 | 3 | 2 | 0.6404 | 2 |
| NODE_168_length_327_cov_1.080000 | 1.7706 | 327 | 0.7187 | 100.00 | 327 | 2 | 2 | 0.7237 | 2 |
| NODE_169_length_327_cov_0.952000 | 1.6697 | 327 | 0.7187 | 100.00 | 327 | 2 | 2 | 0.7198 | 1 |
| NODE_170_length_327_cov_0.788000 | 1.3792 | 327 | 0.6177 | 100.00 | 327 | 2 | 1 | 0.604 | 1 |
| NODE_171_length_327_cov_0.680000 | 1.737 | 327 | 0.7003 | 100.00 | 327 | 2 | 3 | 0.6901 | 2 |
| NODE_172_length_326_cov_1.265060 | 3.2423 | 326 | 0.638 | 100.00 | 326 | 4 | 3 | 0.6594 | 2 |
| NODE_173_length_326_cov_0.851406 | 1.3896 | 326 | 0.6963 | 100.00 | 326 | 2 | 1 | 0.6998 | 1 |
| NODE_174_length_325_cov_0.887097 | 1.3877 | 325 | 0.6338 | 100.00 | 325 | 1 | 2 | 0.6098 | 1 |
| NODE_175_length_324_cov_0.874494 | 1.6451 | 324 | 0.679 | 100.00 | 324 | 2 | 2 | 0.6754 | 1 |
| NODE_176_length_323_cov_1.227642 | 2.6904 | 323 | 0.6223 | 100.00 | 323 | 3 | 4 | 0.6078 | 3 |
| NODE_177_length_323_cov_0.890244 | 1.8452 | 323 | 0.6656 | 100.00 | 323 | 3 | 2 | 0.6678 | 1 |
| NODE_178_length_322_cov_0.848980 | 2.3323 | 322 | 0.646 | 100.00 | 322 | 3 | 3 | 0.6298 | 2 |
| NODE_179_length_322_cov_0.848980 | 1.7329 | 322 | 0.736 | 100.00 | 322 | 1 | 3 | 0.7383 | 2 |
| NODE_180_length_322_cov_0.742857 | 2.1863 | 322 | 0.6522 | 100.00 | 322 | 2 | 3 | 0.6264 | 2 |
| NODE_181_length_320_cov_0.901235 | 1.4094 | 320 | 0.7188 | 100.00 | 320 | 1 | 2 | 0.7073 | 1 |
| NODE_182_length_318_cov_1.199170 | 1.8836 | 318 | 0.5912 | 100.00 | 318 | 3 | 1 | 0.5967 | 2 |
| NODE_183_length_317_cov_0.916667 | 1.4227 | 317 | 0.6972 | 100.00 | 317 | 2 | 1 | 0.6829 | 1 |
| NODE_184_length_316_cov_1.230126 | 2.2373 | 316 | 0.6772 | 100.00 | 316 | 3 | 3 | 0.6832 | 2 |
| NODE_185_length_315_cov_1.852941 | 2.8667 | 315 | 0.6159 | 100.00 | 315 | 3 | 3 | 0.6058 | 2 |
| NODE_186_length_315_cov_1.121849 | 1.9016 | 315 | 0.7492 | 100.00 | 315 | 2 | 2 | 0.7479 | 2 |
| NODE_187_length_315_cov_0.882353 | 1.4317 | 315 | 0.6571 | 100.00 | 315 | 2 | 1 | 0.6519 | 1 |
| NODE_188_length_315_cov_0.857143 | 1.673 | 315 | 0.6984 | 100.00 | 315 | 2 | 2 | 0.7078 | 2 |
| NODE_189_length_315_cov_0.668067 | 1.4349 | 315 | 0.7175 | 100.00 | 315 | 1 | 2 | 0.7212 | 1 |
| NODE_190_length_315_cov_0.609244 | 0.9556 | 315 | 0.6413 | 82.54 | 260 | 1 | 1 | 0.6578 | 1 |
| NODE_191_length_314_cov_1.033755 | 1.9045 | 314 | 0.6242 | 100.00 | 314 | 2 | 2 | 0.6171 | 2 |
| NODE_192_length_313_cov_1.521186 | 2.869 | 313 | 0.6741 | 100.00 | 313 | 4 | 2 | 0.667 | 3 |
| NODE_193_length_313_cov_1.432203 | 2.3962 | 313 | 0.655 | 100.00 | 313 | 4 | 1 | 0.6493 | 2 |

| | | | | | | | | | |
|---|---|---|---|---|---|---|---|---|---|
| NODE_194_length_313_cov_1.224576 | 1.9073 | 313 | 0.508 | 100.00 | 313 | 2 | 2 | 0.4975 | 2 |
| NODE_195_length_313_cov_1.207627 | 2.3898 | 313 | 0.6422 | 100.00 | 313 | 2 | 3 | 0.6613 | 2 |
| NODE_196_length_313_cov_1.131356 | 2.3962 | 313 | 0.722 | 100.00 | 313 | 2 | 3 | 0.7333 | 2 |
| NODE_197_length_313_cov_0.927966 | 1.4409 | 313 | 0.623 | 100.00 | 313 | 1 | 2 | 0.615 | 1 |
| NODE_198_length_313_cov_0.872881 | 1.9169 | 313 | 0.6454 | 100.00 | 313 | 1 | 3 | 0.6433 | 2 |
| NODE_199_length_312_cov_1.089362 | 1.9327 | 312 | 0.6699 | 100.00 | 312 | 1 | 3 | 0.66 | 1 |
| NODE_200_length_312_cov_0.931915 | 1.4423 | 312 | 0.6154 | 100.00 | 312 | 1 | 2 | 0.6098 | 1 |
| NODE_201_length_312_cov_0.676596 | 1.4519 | 312 | 0.6058 | 100.00 | 312 | 2 | 1 | 0.6159 | 1 |
| NODE_202_length_311_cov_1.363248 | 2.4051 | 311 | 0.6688 | 100.00 | 311 | 2 | 3 | 0.6751 | 2 |
| NODE_203_length_311_cov_0.940171 | 1.9325 | 311 | 0.7138 | 100.00 | 311 | 3 | 1 | 0.7148 | 2 |
| NODE_204_length_310_cov_1.184549 | 2.4194 | 310 | 0.5839 | 100.00 | 310 | 2 | 3 | 0.5827 | 2 |
| NODE_205_length_310_cov_0.862661 | 1.9032 | 310 | 0.7355 | 100.00 | 310 | 2 | 2 | 0.7463 | 2 |
| NODE_206_length_309_cov_0.991379 | 1.7443 | 309 | 0.7087 | 100.00 | 309 | 2 | 2 | 0.718 | 2 |
| NODE_207_length_309_cov_0.948276 | 1.9482 | 309 | 0.7023 | 100.00 | 309 | 1 | 3 | 0.7076 | 2 |
| NODE_208_length_308_cov_1.116883 | 2.8929 | 308 | 0.5487 | 100.00 | 308 | 4 | 5 | 0.5163 | 2 |
| NODE_209_length_308_cov_0.969697 | 2.6786 | 308 | 0.5649 | 100.00 | 308 | 4 | 2 | 0.5516 | 3 |
| NODE_210_length_307_cov_1.269565 | 1.9609 | 307 | 0.6482 | 100.00 | 307 | 2 | 2 | 0.6645 | 2 |
| NODE_211_length_306_cov_0.768559 | 1.8954 | 306 | 0.7288 | 100.00 | 306 | 3 | 1 | 0.7438 | 2 |
| NODE_212_length_306_cov_0.633188 | 1.4673 | 306 | 0.6569 | 100.00 | 306 | 2 | 1 | 0.6667 | 1 |
| NODE_213_length_305_cov_1.311404 | 2.318 | 305 | 0.6689 | 100.00 | 305 | 3 | 2 | 0.6719 | 2 |
| NODE_214_length_305_cov_0.956140 | 1.4721 | 305 | 0.718 | 100.00 | 305 | 1 | 2 | 0.7127 | 1 |
| NODE_215_length_305_cov_0.951754 | 1.977 | 305 | 0.6852 | 100.00 | 305 | 1 | 3 | 0.6965 | 2 |
| NODE_216_length_304_cov_2.114537 | 3.8717 | 304 | 0.5691 | 100.00 | 304 | 5 | 4 | 0.5777 | 4 |
| NODE_217_length_304_cov_1.167401 | 1.9836 | 304 | 0.7007 | 100.00 | 304 | 3 | 1 | 0.7065 | 2 |
| NODE_218_length_304_cov_0.762115 | 1.4803 | 304 | 0.5461 | 100.00 | 304 | 1 | 2 | 0.5444 | 1 |
| NODE_219_length_303_cov_0.946903 | 1.9736 | 303 | 0.6304 | 100.00 | 303 | 3 | 1 | 0.6156 | 2 |
| NODE_220_length_302_cov_1.568889 | 2.4801 | 302 | 0.745 | 100.00 | 302 | 4 | 1 | 0.761 | 2 |
| NODE_221_length_301_cov_0.982143 | 1.5017 | 301 | 0.6844 | 100.00 | 301 | 2 | 1 | 0.6858 | 2 |
| NODE_222_length_301_cov_0.968750 | 1.4983 | 301 | 0.6379 | 100.00 | 301 | 2 | 1 | 0.6452 | 1 |
| NODE_223_length_301_cov_0.892857 | 1.9934 | 301 | 0.7043 | 100.00 | 301 | 2 | 2 | 0.6955 | 2 |
| NODE_224_length_300_cov_0.739910 | 1.4967 | 300 | 0.6733 | 100.00 | 300 | 2 | 1 | 0.6622 | 1 |
| NODE_225_length_300_cov_0.632287 | 1.82 | 300 | 0.6867 | 100.00 | 300 | 3 | 1 | 0.6489 | 2 |
| NODE_226_length_299_cov_1.648649 | 2.5117 | 299 | 0.6923 | 100.00 | 299 | 2 | 3 | 0.6924 | 3 |
| NODE_227_length_298_cov_1.348416 | 2.2953 | 298 | 0.6644 | 100.00 | 298 | 3 | 2 | 0.6711 | 2 |
| NODE_228_length_298_cov_1.067873 | 2.0168 | 298 | 0.7081 | 100.00 | 298 | 0 | 4 | 0.7027 | 2 |
| NODE_229_length_298_cov_0.986425 | 2.0168 | 298 | 0.5671 | 100.00 | 298 | 2 | 2 | 0.5674 | 2 |
| NODE_230_length_297_cov_1.381818 | 2.5084 | 297 | 0.4512 | 100.00 | 297 | 2 | 4 | 0.4456 | 2 |
| NODE_231_length_296_cov_1.068493 | 1.8311 | 296 | 0.4223 | 100.00 | 296 | 2 | 2 | 0.4059 | 1 |
| NODE_232_length_296_cov_1.009132 | 1.527 | 296 | 0.7432 | 100.00 | 296 | 2 | 1 | 0.7412 | 2 |

| | | | | | | | | | |
|---|---|---|---|---|---|---|---|---|---|
| NODE_233_length_296_cov_0.981735 | 1.9426 | 296 | 0.6791 | 100.00 | 296 | 3 | 1 | 0.6872 | 2 |
| NODE_234_length_296_cov_0.808219 | 1.5203 | 296 | 0.5338 | 100.00 | 296 | 2 | 1 | 0.5222 | 2 |
| NODE_235_length_295_cov_1.165138 | 1.9051 | 295 | 0.6949 | 100.00 | 295 | 3 | 1 | 0.7072 | 2 |
| NODE_236_length_294_cov_1.649770 | 3.0306 | 294 | 0.6497 | 100.00 | 294 | 4 | 2 | 0.6588 | 3 |
| NODE_237_length_294_cov_0.976959 | 2.1633 | 294 | 0.6565 | 100.00 | 294 | 3 | 2 | 0.6658 | 2 |
| NODE_238_length_294_cov_0.857143 | 1.5272 | 294 | 0.6429 | 100.00 | 294 | 1 | 2 | 0.6481 | 2 |
| NODE_239_length_294_cov_0.857143 | 1.5408 | 294 | 0.6871 | 100.00 | 294 | 2 | 1 | 0.6887 | 2 |
| NODE_240_length_294_cov_0.820276 | 1.5374 | 294 | 0.6565 | 100.00 | 294 | 3 | 0 | 0.6659 | 2 |
| NODE_241_length_293_cov_1.722222 | 3.3618 | 293 | 0.587 | 100.00 | 293 | 4 | 4 | 0.6376 | 4 |
| NODE_242_length_293_cov_1.657407 | 2.5666 | 293 | 0.7065 | 100.00 | 293 | 2 | 3 | 0.7154 | 3 |
| NODE_243_length_293_cov_0.925926 | 1.5427 | 293 | 0.6314 | 100.00 | 293 | 2 | 1 | 0.6394 | 2 |
| NODE_244_length_293_cov_0.884259 | 2.0444 | 293 | 0.5495 | 100.00 | 293 | 2 | 2 | 0.5543 | 2 |
| NODE_245_length_291_cov_1.023364 | 1.5464 | 291 | 0.6598 | 100.00 | 291 | 1 | 2 | 0.6689 | 2 |
| NODE_246_length_291_cov_0.897196 | 1.457 | 291 | 0.732 | 100.00 | 291 | 1 | 2 | 0.7206 | 1 |
| NODE_247_length_291_cov_0.803738 | 1.6838 | 291 | 0.6048 | 100.00 | 291 | 3 | 2 | 0.625 | 2 |
| NODE_248_length_290_cov_1.380282 | 2.5966 | 290 | 0.6069 | 100.00 | 290 | 2 | 3 | 0.5963 | 3 |
| NODE_249_length_290_cov_1.037559 | 1.5586 | 290 | 0.6655 | 100.00 | 290 | 2 | 1 | 0.6681 | 2 |
| NODE_250_length_290_cov_1.032864 | 1.5552 | 290 | 0.7138 | 100.00 | 290 | 2 | 1 | 0.7073 | 2 |
| NODE_251_length_290_cov_0.957746 | 1.5 | 290 | 0.6724 | 100.00 | 290 | 2 | 1 | 0.6563 | 1 |
| NODE_252_length_290_cov_0.868545 | 1.5552 | 290 | 0.7414 | 100.00 | 290 | 2 | 1 | 0.7384 | 2 |
| NODE_253_length_289_cov_1.547170 | 2.4775 | 289 | 0.6747 | 100.00 | 289 | 3 | 2 | 0.669 | 2 |
| NODE_254_length_289_cov_1.523585 | 3.1246 | 289 | 0.6609 | 100.00 | 289 | 3 | 3 | 0.6645 | 3 |
| NODE_255_length_289_cov_1.216981 | 2.7509 | 289 | 0.7197 | 100.00 | 289 | 4 | 3 | 0.7283 | 2 |
| NODE_256_length_289_cov_1.198113 | 2.6055 | 289 | 0.7128 | 100.00 | 289 | 2 | 3 | 0.7185 | 3 |
| NODE_257_length_289_cov_1.037736 | 1.5606 | 289 | 0.692 | 100.00 | 289 | 2 | 1 | 0.694 | 2 |
| NODE_258_length_289_cov_0.919811 | 1.5606 | 289 | 0.6471 | 100.00 | 289 | 1 | 2 | 0.6585 | 2 |
| NODE_259_length_289_cov_0.872642 | 1.5675 | 289 | 0.699 | 100.00 | 289 | 1 | 2 | 0.7174 | 2 |
| NODE_260_length_289_cov_0.693396 | 1.5571 | 289 | 0.6228 | 100.00 | 289 | 1 | 2 | 0.6378 | 2 |
| NODE_261_length_288_cov_1.672986 | 2.8333 | 288 | 0.4201 | 100.00 | 288 | 3 | 3 | 0.3946 | 2 |
| NODE_262_length_288_cov_1.289100 | 2.0243 | 288 | 0.691 | 100.00 | 288 | 2 | 2 | 0.693 | 2 |
| NODE_263_length_288_cov_1.184834 | 2.0833 | 288 | 0.7431 | 100.00 | 288 | 3 | 1 | 0.7342 | 2 |
| NODE_264_length_288_cov_1.047393 | 1.5694 | 288 | 0.4444 | 100.00 | 288 | 1 | 2 | 0.4602 | 2 |
| NODE_265_length_288_cov_0.890995 | 1.5625 | 288 | 0.7257 | 100.00 | 288 | 2 | 1 | 0.7089 | 2 |
| NODE_266_length_288_cov_0.597156 | 1.375 | 288 | 0.7326 | 100.00 | 288 | 2 | 1 | 0.7261 | 1 |
| NODE_267_length_287_cov_1.380952 | 2.0906 | 287 | 0.5819 | 100.00 | 287 | 2 | 2 | 0.625 | 2 |
| NODE_268_length_287_cov_1.038095 | 1.5645 | 287 | 0.662 | 100.00 | 287 | 1 | 2 | 0.6615 | 2 |
| NODE_269_length_287_cov_1.014286 | 1.5714 | 287 | 0.6725 | 100.00 | 287 | 2 | 1 | 0.6741 | 2 |
| NODE_270_length_287_cov_0.738095 | 1.9686 | 287 | 0.7108 | 100.00 | 287 | 2 | 2 | 0.6977 | 2 |
| NODE_271_length_286_cov_1.746411 | 2.6294 | 286 | 0.7028 | 100.00 | 286 | 3 | 2 | 0.7088 | 3 |

| | | | | | | | | | |
|---|---|---|---|---|---|---|---|---|---|
| NODE_272_length_286_cov_1.047847 | 1.5734 | 286 | 0.5455 | 100.00 | 286 | 1 | 2 | 0.54 | 2 |
| NODE_273_length_286_cov_1.043062 | 1.5734 | 286 | 0.4755 | 100.00 | 286 | 1 | 2 | 0.4533 | 2 |
| NODE_274_length_286_cov_0.593301 | 1.5664 | 286 | 0.6818 | 100.00 | 286 | 2 | 2 | 0.6942 | 1 |
| NODE_275_length_285_cov_1.408654 | 2.1088 | 285 | 0.5965 | 100.00 | 285 | 2 | 2 | 0.6007 | 2 |
| NODE_276_length_285_cov_1.389423 | 2.1123 | 285 | 0.7123 | 100.00 | 285 | 3 | 1 | 0.7027 | 2 |
| NODE_277_length_285_cov_0.975962 | 1.5825 | 285 | 0.5719 | 100.00 | 285 | 2 | 1 | 0.5686 | 2 |
| NODE_278_length_285_cov_0.927885 | 1.5789 | 285 | 0.4246 | 100.00 | 285 | 2 | 1 | 0.4178 | 2 |
| NODE_279_length_285_cov_0.697115 | 1.3719 | 285 | 0.6211 | 100.00 | 285 | 2 | 1 | 0.6173 | 1 |
| NODE_280_length_284_cov_1.115942 | 2.4437 | 284 | 0.662 | 100.00 | 284 | 3 | 3 | 0.6542 | 2 |
| NODE_281_length_283_cov_1.509709 | 2.4664 | 283 | 0.6184 | 100.00 | 283 | 3 | 2 | 0.6103 | 2 |
| NODE_282_length_283_cov_1.305825 | 2.1272 | 283 | 0.7138 | 100.00 | 283 | 3 | 1 | 0.7076 | 2 |
| NODE_283_length_282_cov_0.956098 | 1.5922 | 282 | 0.6596 | 100.00 | 282 | 1 | 2 | 0.6615 | 2 |
| NODE_284_length_281_cov_0.857843 | 1.7189 | 281 | 0.6548 | 100.00 | 281 | 2 | 2 | 0.6253 | 1 |
| NODE_285_length_281_cov_0.725490 | 1.0747 | 281 | 0.6868 | 100.00 | 281 | 1 | 1 | 0.6921 | 1 |
| NODE_286_length_281_cov_0.710784 | 1.0712 | 281 | 0.7011 | 89.68 | 252 | 1 | 1 | 0.691 | 1 |
| NODE_287_length_280_cov_1.305419 | 2.15 | 280 | 0.4786 | 100.00 | 280 | 2 | 2 | 0.4817 | 2 |
| NODE_288_length_280_cov_0.891626 | 1.6036 | 280 | 0.6821 | 100.00 | 280 | 1 | 2 | 0.6978 | 2 |
| NODE_289_length_280_cov_0.724138 | 1.075 | 280 | 0.65 | 100.00 | 280 | 1 | 1 | 0.6478 | 1 |
| NODE_290_length_280_cov_0.714286 | 2.9714 | 280 | 0.5964 | 100.00 | 280 | 3 | 3 | 0.5979 | 3 |
| NODE_291_length_279_cov_0.975248 | 1.6165 | 279 | 0.7348 | 100.00 | 279 | 1 | 2 | 0.7184 | 2 |
| NODE_292_length_279_cov_0.896040 | 2.0609 | 279 | 0.6057 | 100.00 | 279 | 4 | 1 | 0.5878 | 2 |
| NODE_293_length_279_cov_0.722772 | 1.0753 | 279 | 0.6703 | 100.00 | 279 | 1 | 1 | 0.6667 | 1 |
| NODE_294_length_278_cov_1.805970 | 3.2338 | 278 | 0.6367 | 100.00 | 278 | 3 | 3 | 0.6229 | 4 |
| NODE_295_length_278_cov_1.009950 | 1.6259 | 278 | 0.554 | 100.00 | 278 | 1 | 2 | 0.573 | 2 |
| NODE_296_length_278_cov_0.726368 | 1.0791 | 278 | 0.5791 | 100.00 | 278 | 2 | 0 | 0.5781 | 1 |
| NODE_297_length_278_cov_0.726368 | 1.0791 | 278 | 0.6223 | 100.00 | 278 | 1 | 1 | 0.6267 | 1 |
| NODE_298_length_278_cov_0.726368 | 1.0791 | 278 | 0.723 | 100.00 | 278 | 2 | 0 | 0.7333 | 1 |
| NODE_299_length_277_cov_1.100000 | 1.6282 | 277 | 0.7292 | 100.00 | 277 | 1 | 2 | 0.7428 | 2 |
| NODE_300_length_277_cov_0.955000 | 2.3827 | 277 | 0.4946 | 100.00 | 277 | 3 | 3 | 0.4958 | 3 |
| NODE_301_length_277_cov_0.855000 | 1.6209 | 277 | 0.6173 | 100.00 | 277 | 2 | 1 | 0.612 | 2 |
| NODE_302_length_277_cov_0.735000 | 1.0866 | 277 | 0.6823 | 100.00 | 277 | 2 | 0 | 0.6877 | 1 |
| NODE_303_length_277_cov_0.735000 | 1.0866 | 277 | 0.6895 | 100.00 | 277 | 1 | 1 | 0.6744 | 1 |
| NODE_304_length_277_cov_0.725000 | 1.0794 | 277 | 0.6209 | 100.00 | 277 | 1 | 1 | 0.6221 | 1 |
| NODE_305_length_276_cov_1.090452 | 2.0833 | 276 | 0.6812 | 100.00 | 276 | 2 | 2 | 0.6643 | 2 |
| NODE_306_length_276_cov_1.040201 | 1.6304 | 276 | 0.7355 | 100.00 | 276 | 1 | 2 | 0.7267 | 2 |
| NODE_307_length_276_cov_1.025126 | 2.1812 | 276 | 0.7391 | 100.00 | 276 | 1 | 3 | 0.7313 | 2 |
| NODE_308_length_276_cov_0.879397 | 2.1232 | 276 | 0.6486 | 100.00 | 276 | 4 | 1 | 0.6331 | 2 |
| NODE_309_length_276_cov_0.738693 | 1.6196 | 276 | 0.6377 | 100.00 | 276 | 2 | 2 | 0.6555 | 1 |
| NODE_310_length_276_cov_0.738693 | 1.0906 | 276 | 0.4964 | 100.00 | 276 | 1 | 1 | 0.495 | 1 |

| | | | | | | | | | |
|---|---|---|---|---|---|---|---|---|---|
| NODE_311_length_276_cov_0.738693 | 1.0906 | 276 | 0.7101 | 100.00 | 276 | 1 | 1 | 0.7076 | 1 |
| NODE_312_length_276_cov_0.733668 | 1.087 | 276 | 0.7464 | 100.00 | 276 | 1 | 1 | 0.7467 | 1 |
| NODE_313_length_276_cov_0.728643 | 1.0833 | 276 | 0.6486 | 100.00 | 276 | 1 | 1 | 0.6488 | 1 |
| NODE_314_length_276_cov_0.728643 | 1.0906 | 276 | 0.7283 | 100.00 | 276 | 1 | 1 | 0.7409 | 1 |
| NODE_315_length_276_cov_0.723618 | 2.1087 | 276 | 0.7319 | 100.00 | 276 | 3 | 1 | 0.7525 | 2 |
| NODE_316_length_276_cov_0.723618 | 1.0833 | 276 | 0.7101 | 100.00 | 276 | 1 | 1 | 0.7157 | 1 |
| NODE_317_length_275_cov_1.494949 | 2.1964 | 275 | 0.6073 | 100.00 | 275 | 2 | 2 | 0.6159 | 2 |
| NODE_318_length_275_cov_1.474747 | 2.1818 | 275 | 0.6909 | 89.09 | 245 | 2 | 2 | 0.7033 | 2 |
| NODE_319_length_275_cov_1.146465 | 1.9491 | 275 | 0.6945 | 100.00 | 275 | 2 | 2 | 0.6903 | 1 |
| NODE_320_length_275_cov_0.742424 | 1.0945 | 275 | 0.6364 | 100.00 | 275 | 1 | 1 | 0.6346 | 1 |
| NODE_321_length_275_cov_0.742424 | 1.6436 | 275 | 0.7309 | 100.00 | 275 | 1 | 2 | 0.75 | 2 |
| NODE_322_length_275_cov_0.742424 | 1.0945 | 275 | 0.5673 | 100.00 | 275 | 2 | 0 | 0.5748 | 1 |
| NODE_323_length_275_cov_0.742424 | 1.0945 | 275 | 0.5273 | 100.00 | 275 | 1 | 1 | 0.5249 | 1 |
| NODE_324_length_274_cov_0.741117 | 1.0949 | 274 | 0.4891 | 100.00 | 274 | 1 | 1 | 0.4767 | 1 |
| NODE_325_length_274_cov_0.741117 | 1.0949 | 274 | 0.7153 | 100.00 | 274 | 1 | 1 | 0.7067 | 1 |
| NODE_326_length_274_cov_0.741117 | 1.0985 | 274 | 0.6752 | 100.00 | 274 | 1 | 1 | 0.6744 | 1 |
| NODE_327_length_274_cov_0.741117 | 1.0949 | 274 | 0.6496 | 100.00 | 274 | 1 | 1 | 0.6533 | 1 |
| NODE_328_length_274_cov_0.736041 | 1.0912 | 274 | 0.5693 | 100.00 | 274 | 1 | 1 | 0.5719 | 1 |
| NODE_329_length_273_cov_1.122449 | 1.652 | 273 | 0.6923 | 100.00 | 273 | 1 | 2 | 0.6874 | 2 |
| NODE_330_length_273_cov_0.795918 | 1.6447 | 273 | 0.6447 | 100.00 | 273 | 1 | 2 | 0.637 | 2 |
| NODE_331_length_273_cov_0.755102 | 1.1062 | 273 | 0.6484 | 100.00 | 273 | 1 | 1 | 0.649 | 1 |
| NODE_332_length_273_cov_0.750000 | 1.3919 | 273 | 0.652 | 100.00 | 273 | 1 | 2 | 0.6474 | 1 |
| NODE_333_length_273_cov_0.750000 | 1.1026 | 273 | 0.6484 | 100.00 | 273 | 1 | 1 | 0.6512 | 1 |
| NODE_334_length_273_cov_0.744898 | 1.0989 | 273 | 0.5934 | 100.00 | 273 | 1 | 1 | 0.5833 | 1 |
| NODE_335_length_273_cov_0.734694 | 1.0989 | 273 | 0.641 | 100.00 | 273 | 1 | 1 | 0.64 | 1 |
| NODE_336_length_272_cov_1.297436 | 2.2132 | 272 | 0.6801 | 100.00 | 272 | 3 | 1 | 0.6993 | 2 |
| NODE_337_length_272_cov_0.758974 | 1.1103 | 272 | 0.7096 | 100.00 | 272 | 1 | 1 | 0.7185 | 1 |
| NODE_338_length_272_cov_0.753846 | 1.5441 | 272 | 0.6801 | 100.00 | 272 | 1 | 2 | 0.6452 | 2 |
| NODE_339_length_272_cov_0.748718 | 1.1029 | 272 | 0.5699 | 100.00 | 272 | 1 | 1 | 0.5567 | 1 |
| NODE_340_length_272_cov_0.748718 | 1.1029 | 272 | 0.7353 | 100.00 | 272 | 1 | 1 | 0.7333 | 1 |
| NODE_341_length_272_cov_0.743590 | 1.1066 | 272 | 0.5919 | 100.00 | 272 | 1 | 1 | 0.5914 | 1 |
| NODE_342_length_271_cov_1.505155 | 2.2214 | 271 | 0.6347 | 100.00 | 271 | 2 | 2 | 0.6196 | 2 |
| NODE_343_length_271_cov_1.371134 | 2.2288 | 271 | 0.631 | 100.00 | 271 | 2 | 2 | 0.6308 | 2 |
| NODE_344_length_271_cov_1.139175 | 1.6679 | 271 | 0.69 | 100.00 | 271 | 2 | 1 | 0.6991 | 2 |
| NODE_345_length_271_cov_1.128866 | 1.6605 | 271 | 0.6716 | 100.00 | 271 | 1 | 2 | 0.6689 | 2 |
| NODE_346_length_271_cov_1.041237 | 1.6642 | 271 | 0.5609 | 100.00 | 271 | 1 | 2 | 0.5543 | 2 |
| NODE_347_length_271_cov_0.757732 | 1.1107 | 271 | 0.6384 | 100.00 | 271 | 1 | 1 | 0.6412 | 1 |
| NODE_348_length_271_cov_0.752577 | 1.1107 | 271 | 0.6605 | 100.00 | 271 | 1 | 1 | 0.6678 | 1 |
| NODE_349_length_271_cov_0.752577 | 1.107 | 271 | 0.6827 | 100.00 | 271 | 1 | 1 | 0.68 | 1 |

| | | | | | | | | | |
|---|---|---|---|---|---|---|---|---|---|
| NODE_350_length_270_cov_1.336788 | 2.0963 | 270 | 0.6519 | 100.00 | 270 | 2 | 2 | 0.6519 | 2 |
| NODE_351_length_270_cov_1.000000 | 1.6778 | 270 | 0.7667 | 100.00 | 270 | 2 | 1 | 0.7748 | 2 |
| NODE_352_length_270_cov_0.829016 | 2.0889 | 270 | 0.7185 | 100.00 | 270 | 1 | 3 | 0.7038 | 2 |
| NODE_353_length_270_cov_0.766839 | 1.1185 | 270 | 0.4111 | 100.00 | 270 | 1 | 1 | 0.394 | 1 |
| NODE_354_length_270_cov_0.761658 | 1.1148 | 270 | 0.6852 | 100.00 | 270 | 1 | 1 | 0.6811 | 1 |
| NODE_355_length_270_cov_0.756477 | 1.1111 | 270 | 0.6667 | 100.00 | 270 | 2 | 0 | 0.6667 | 1 |
| NODE_356_length_270_cov_0.756477 | 1.1148 | 270 | 0.6704 | 100.00 | 270 | 1 | 1 | 0.6744 | 1 |
| NODE_357_length_270_cov_0.751295 | 1.1111 | 270 | 0.6556 | 100.00 | 270 | 1 | 1 | 0.6633 | 1 |
| NODE_358_length_270_cov_0.751295 | 1.1074 | 270 | 0.6556 | 100.00 | 270 | 1 | 1 | 0.65 | 1 |
| NODE_359_length_270_cov_0.751295 | 1.1074 | 270 | 0.6926 | 100.00 | 270 | 1 | 1 | 0.6867 | 1 |
| NODE_360_length_269_cov_2.072917 | 4.2677 | 269 | 0.6989 | 100.00 | 269 | 5 | 3 | 0.719 | 5 |
| NODE_361_length_269_cov_1.718750 | 2.7546 | 269 | 0.6059 | 100.00 | 269 | 3 | 2 | 0.5924 | 3 |
| NODE_362_length_269_cov_1.140625 | 1.6766 | 269 | 0.6543 | 100.00 | 269 | 2 | 1 | 0.6497 | 2 |
| NODE_363_length_269_cov_1.140625 | 1.6729 | 269 | 0.6617 | 100.00 | 269 | 2 | 1 | 0.6622 | 2 |
| NODE_364_length_269_cov_1.036458 | 2.3978 | 269 | 0.632 | 100.00 | 269 | 2 | 3 | 0.6403 | 2 |
| NODE_365_length_269_cov_0.770833 | 1.6803 | 269 | 0.6543 | 100.00 | 269 | 2 | 1 | 0.6452 | 2 |
| NODE_366_length_268_cov_1.523560 | 2.2351 | 268 | 0.6493 | 100.00 | 268 | 2 | 2 | 0.6427 | 3 |
| NODE_367_length_268_cov_1.157068 | 1.6866 | 268 | 0.7351 | 100.00 | 268 | 2 | 1 | 0.7367 | 2 |
| NODE_368_length_268_cov_1.141361 | 1.6791 | 268 | 0.6679 | 100.00 | 268 | 1 | 2 | 0.6733 | 2 |
| NODE_369_length_268_cov_1.125654 | 1.6642 | 268 | 0.709 | 100.00 | 268 | 2 | 1 | 0.7063 | 2 |
| NODE_370_length_268_cov_0.968586 | 2.2537 | 268 | 0.694 | 100.00 | 268 | 2 | 2 | 0.702 | 2 |
| NODE_371_length_268_cov_0.774869 | 1.1269 | 268 | 0.6455 | 100.00 | 268 | 1 | 1 | 0.649 | 1 |
| NODE_372_length_268_cov_0.769634 | 1.1231 | 268 | 0.5373 | 100.00 | 268 | 1 | 1 | 0.5515 | 1 |
| NODE_373_length_268_cov_0.769634 | 1.1269 | 268 | 0.6866 | 100.00 | 268 | 1 | 1 | 0.6755 | 1 |
| NODE_374_length_268_cov_0.769634 | 1.1269 | 268 | 0.694 | 100.00 | 268 | 1 | 1 | 0.702 | 1 |
| NODE_375_length_268_cov_0.764398 | 1.1231 | 268 | 0.5784 | 100.00 | 268 | 1 | 1 | 0.588 | 1 |
| NODE_376_length_268_cov_0.759162 | 1.6828 | 268 | 0.7015 | 100.00 | 268 | 1 | 2 | 0.7162 | 2 |
| NODE_377_length_268_cov_0.722513 | 1.1194 | 268 | 0.7015 | 100.00 | 268 | 0 | 2 | 0.7067 | 1 |
| NODE_378_length_267_cov_1.152632 | 1.6854 | 267 | 0.6105 | 100.00 | 267 | 1 | 2 | 0.5907 | 2 |
| NODE_379_length_267_cov_0.778947 | 1.1311 | 267 | 0.7378 | 100.00 | 267 | 1 | 1 | 0.7384 | 1 |
| NODE_380_length_267_cov_0.773684 | 1.1273 | 267 | 0.6592 | 100.00 | 267 | 1 | 1 | 0.6645 | 1 |
| NODE_381_length_267_cov_0.768421 | 1.1236 | 267 | 0.603 | 100.00 | 267 | 1 | 1 | 0.58 | 1 |
| NODE_382_length_267_cov_0.768421 | 1.1236 | 267 | 0.633 | 100.00 | 267 | 1 | 1 | 0.63 | 1 |
| NODE_383_length_267_cov_0.768421 | 1.1236 | 267 | 0.633 | 100.00 | 267 | 1 | 1 | 0.6333 | 1 |
| NODE_384_length_267_cov_0.763158 | 1.6854 | 267 | 0.6442 | 100.00 | 267 | 2 | 1 | 0.6422 | 2 |
| NODE_385_length_267_cov_0.763158 | 1.6891 | 267 | 0.6105 | 100.00 | 267 | 2 | 1 | 0.5876 | 2 |
| NODE_386_length_267_cov_0.647368 | 1.5019 | 267 | 0.6929 | 100.00 | 267 | 1 | 2 | 0.6903 | 2 |
| NODE_387_length_266_cov_1.772487 | 2.8233 | 266 | 0.5113 | 100.00 | 266 | 3 | 2 | 0.5007 | 3 |
| NODE_388_length_266_cov_0.947090 | 2.1842 | 266 | 0.5714 | 100.00 | 266 | 2 | 2 | 0.5399 | 2 |

| | | | | | | | | | |
|---|---|---|---|---|---|---|---|---|---|
| NODE_389_length_266_cov_0.857143 | 1.6241 | 266 | 0.6917 | 100.00 | 266 | 1 | 2 | 0.6807 | 2 |
| NODE_390_length_266_cov_0.783069 | 1.6015 | 266 | 0.6316 | 100.00 | 266 | 2 | 2 | 0.6419 | 1 |
| NODE_391_length_266_cov_0.777778 | 1.703 | 266 | 0.6692 | 100.00 | 266 | 2 | 2 | 0.6799 | 1 |
| NODE_392_length_266_cov_0.777778 | 1.1316 | 266 | 0.6617 | 100.00 | 266 | 1 | 1 | 0.6645 | 1 |
| NODE_393_length_266_cov_0.772487 | 1.1278 | 266 | 0.6316 | 100.00 | 266 | 1 | 1 | 0.6467 | 1 |
| NODE_394_length_265_cov_2.218085 | 3.4 | 265 | 0.4755 | 100.00 | 265 | 3 | 3 | 0.4772 | 4 |
| NODE_395_length_265_cov_1.542553 | 2.9887 | 265 | 0.6906 | 100.00 | 265 | 4 | 2 | 0.6885 | 2 |
| NODE_396_length_265_cov_1.271277 | 2.2 | 265 | 0.6604 | 100.00 | 265 | 1 | 3 | 0.6267 | 2 |
| NODE_397_length_265_cov_1.079787 | 1.7019 | 265 | 0.6679 | 100.00 | 265 | 2 | 1 | 0.6851 | 2 |
| NODE_398_length_265_cov_0.787234 | 1.5698 | 265 | 0.6491 | 100.00 | 265 | 1 | 2 | 0.6504 | 2 |
| NODE_399_length_265_cov_0.781915 | 1.1358 | 265 | 0.6642 | 100.00 | 265 | 1 | 1 | 0.6645 | 1 |
| NODE_400_length_265_cov_0.771277 | 1.1283 | 265 | 0.7358 | 100.00 | 265 | 1 | 1 | 0.7358 | 1 |
| NODE_401_length_265_cov_0.739362 | 1.1321 | 265 | 0.5774 | 100.00 | 265 | 2 | 0 | 0.5667 | 1 |
| NODE_402_length_265_cov_0.734043 | 1.1019 | 265 | 0.7132 | 100.00 | 265 | 1 | 1 | 0.6987 | 1 |
| NODE_403_length_264_cov_1.016043 | 1.7045 | 264 | 0.5455 | 100.00 | 264 | 1 | 2 | 0.5244 | 2 |
| NODE_404_length_264_cov_0.903743 | 1.7045 | 264 | 0.6402 | 100.00 | 264 | 2 | 1 | 0.6244 | 2 |
| NODE_405_length_264_cov_0.786096 | 1.1439 | 264 | 0.6667 | 100.00 | 264 | 1 | 1 | 0.6689 | 1 |
| NODE_406_length_264_cov_0.786096 | 1.1439 | 264 | 0.697 | 100.00 | 264 | 1 | 1 | 0.6954 | 1 |
| NODE_407_length_264_cov_0.780749 | 1.1364 | 264 | 0.6023 | 100.00 | 264 | 1 | 1 | 0.6067 | 1 |
| NODE_408_length_264_cov_0.780749 | 1.1364 | 264 | 0.697 | 100.00 | 264 | 0 | 2 | 0.69 | 1 |
| NODE_409_length_264_cov_0.770053 | 1.1364 | 264 | 0.6061 | 100.00 | 264 | 1 | 1 | 0.6033 | 1 |
| NODE_410_length_264_cov_0.700535 | 1.9886 | 264 | 0.6742 | 100.00 | 264 | 3 | 2 | 0.655 | 2 |
| NODE_411_length_263_cov_1.129032 | 2.5095 | 263 | 0.6198 | 100.00 | 263 | 1 | 4 | 0.6182 | 3 |
| NODE_412_length_263_cov_0.795699 | 1.1483 | 263 | 0.6084 | 100.00 | 263 | 1 | 1 | 0.6126 | 1 |
| NODE_413_length_263_cov_0.795699 | 1.1483 | 263 | 0.6768 | 100.00 | 263 | 1 | 1 | 0.6921 | 1 |
| NODE_414_length_263_cov_0.790323 | 1.1483 | 263 | 0.7034 | 100.00 | 263 | 1 | 1 | 0.7053 | 1 |
| NODE_415_length_263_cov_0.779570 | 1.1407 | 263 | 0.6388 | 100.00 | 263 | 1 | 1 | 0.6367 | 1 |
| NODE_416_length_262_cov_0.989189 | 1.874 | 262 | 0.7137 | 83.97 | 220 | 3 | 1 | 0.6945 | 2 |
| NODE_417_length_262_cov_0.789189 | 1.1489 | 262 | 0.7023 | 100.00 | 262 | 1 | 1 | 0.7143 | 1 |
| NODE_418_length_262_cov_0.783784 | 2.2863 | 262 | 0.4237 | 100.00 | 262 | 2 | 2 | 0.4274 | 2 |
| NODE_419_length_262_cov_0.735135 | 1.1412 | 262 | 0.6603 | 62.21 | 163 | 1 | 1 | 0.6756 | 2 |
| NODE_420_length_261_cov_1.190217 | 1.7241 | 261 | 0.7318 | 100.00 | 261 | 1 | 2 | 0.7444 | 2 |
| NODE_421_length_261_cov_0.902174 | 1.7241 | 261 | 0.6513 | 100.00 | 261 | 2 | 1 | 0.62 | 2 |
| NODE_422_length_261_cov_0.864130 | 2.1111 | 261 | 0.6743 | 100.00 | 261 | 2 | 2 | 0.6806 | 2 |
| NODE_423_length_261_cov_0.804348 | 1.1571 | 261 | 0.7165 | 100.00 | 261 | 1 | 1 | 0.7185 | 1 |
| NODE_424_length_261_cov_0.798913 | 1.1533 | 261 | 0.7203 | 100.00 | 261 | 1 | 1 | 0.7243 | 1 |
| NODE_425_length_261_cov_0.793478 | 1.1494 | 261 | 0.6782 | 100.00 | 261 | 1 | 1 | 0.6744 | 1 |
| NODE_426_length_261_cov_0.793478 | 1.1494 | 261 | 0.4291 | 100.00 | 261 | 1 | 1 | 0.4367 | 1 |
| NODE_427_length_261_cov_0.788043 | 1.1494 | 261 | 0.705 | 100.00 | 261 | 1 | 1 | 0.71 | 1 |

| | | | | | | | | | |
|---|---|---|---|---|---|---|---|---|---|
| NODE_428_length_261_cov_0.788043 | 1.1494 | 261 | 0.5977 | 100.00 | 261 | 1 | 1 | 0.5967 | 1 |
| NODE_429_length_260_cov_1.786885 | 2.7385 | 260 | 0.6808 | 100.00 | 260 | 1 | 4 | 0.6798 | 3 |
| NODE_430_length_260_cov_1.448087 | 2.8885 | 260 | 0.55 | 100.00 | 260 | 3 | 2 | 0.5566 | 3 |
| NODE_431_length_260_cov_0.923497 | 1.7346 | 260 | 0.6808 | 100.00 | 260 | 1 | 2 | 0.6807 | 2 |
| NODE_432_length_260_cov_0.808743 | 1.1615 | 260 | 0.6346 | 100.00 | 260 | 1 | 1 | 0.6325 | 1 |
| NODE_433_length_260_cov_0.803279 | 1.1615 | 260 | 0.6538 | 100.00 | 260 | 1 | 1 | 0.649 | 1 |
| NODE_434_length_260_cov_0.803279 | 1.1577 | 260 | 0.6731 | 100.00 | 260 | 1 | 1 | 0.6645 | 1 |
| NODE_435_length_260_cov_0.803279 | 2.8077 | 260 | 0.5615 | 100.00 | 260 | 3 | 3 | 0.5507 | 2 |
| NODE_436_length_260_cov_0.797814 | 1.1577 | 260 | 0.6231 | 100.00 | 260 | 1 | 1 | 0.6512 | 1 |
| NODE_437_length_260_cov_0.797814 | 1.1577 | 260 | 0.6731 | 100.00 | 260 | 1 | 1 | 0.6678 | 1 |
| NODE_438_length_260_cov_0.792350 | 1.6 | 260 | 0.7308 | 100.00 | 260 | 3 | 0 | 0.719 | 2 |
| NODE_439_length_260_cov_0.792350 | 1.15 | 260 | 0.6346 | 100.00 | 260 | 1 | 1 | 0.6355 | 1 |
| NODE_440_length_260_cov_0.748634 | 1.1538 | 260 | 0.5808 | 100.00 | 260 | 2 | 0 | 0.58 | 1 |
| NODE_441_length_260_cov_0.715847 | 1.1615 | 260 | 0.6923 | 100.00 | 260 | 1 | 1 | 0.6854 | 1 |
| NODE_442_length_260_cov_0.693989 | 1.1538 | 260 | 0.5269 | 100.00 | 260 | 0 | 2 | 0.5233 | 1 |
| NODE_443_length_259_cov_1.170330 | 1.7452 | 259 | 0.7027 | 100.00 | 259 | 1 | 2 | 0.708 | 2 |
| NODE_444_length_259_cov_1.038462 | 1.7452 | 259 | 0.6988 | 100.00 | 259 | 1 | 2 | 0.6947 | 2 |
| NODE_445_length_259_cov_1.000000 | 1.8919 | 259 | 0.7259 | 100.00 | 259 | 2 | 2 | 0.7265 | 2 |
| NODE_446_length_259_cov_0.994505 | 1.7375 | 259 | 0.7181 | 100.00 | 259 | 2 | 1 | 0.7267 | 2 |
| NODE_447_length_259_cov_0.807692 | 1.1622 | 259 | 0.6023 | 100.00 | 259 | 2 | 0 | 0.6179 | 1 |
| NODE_448_length_259_cov_0.807692 | 1.1622 | 259 | 0.6486 | 100.00 | 259 | 1 | 1 | 0.6512 | 1 |
| NODE_449_length_259_cov_0.796703 | 1.1583 | 259 | 0.6911 | 100.00 | 259 | 1 | 1 | 0.6833 | 1 |
| NODE_450_length_258_cov_2.000000 | 2.8992 | 258 | 0.6589 | 100.00 | 258 | 2 | 3 | 0.6707 | 4 |
| NODE_451_length_258_cov_1.624309 | 2.3372 | 258 | 0.6705 | 100.00 | 258 | 3 | 1 | 0.66 | 3 |
| NODE_452_length_258_cov_1.099448 | 1.6705 | 258 | 0.6434 | 100.00 | 258 | 1 | 2 | 0.6386 | 2 |
| NODE_453_length_258_cov_0.812155 | 1.1667 | 258 | 0.6395 | 100.00 | 258 | 1 | 1 | 0.6291 | 1 |
| NODE_454_length_258_cov_0.812155 | 1.1667 | 258 | 0.7054 | 100.00 | 258 | 1 | 1 | 0.7076 | 1 |
| NODE_455_length_258_cov_0.812155 | 1.7519 | 258 | 0.6744 | 100.00 | 258 | 3 | 0 | 0.6593 | 2 |
| NODE_456_length_258_cov_0.806630 | 2.1124 | 258 | 0.6783 | 100.00 | 258 | 2 | 2 | 0.6833 | 2 |
| NODE_457_length_258_cov_0.806630 | 1.1628 | 258 | 0.6628 | 100.00 | 258 | 0 | 2 | 0.6633 | 1 |
| NODE_458_length_258_cov_0.801105 | 1.1589 | 258 | 0.6202 | 100.00 | 258 | 1 | 1 | 0.6221 | 1 |
| NODE_459_length_258_cov_0.734807 | 1.1705 | 258 | 0.6085 | 100.00 | 258 | 1 | 1 | 0.606 | 1 |
| NODE_460_length_257_cov_1.550000 | 2.3424 | 257 | 0.6381 | 100.00 | 257 | 2 | 2 | 0.6213 | 2 |
| NODE_461_length_257_cov_1.372222 | 2.3346 | 257 | 0.6459 | 100.00 | 257 | 3 | 1 | 0.6533 | 3 |
| NODE_462_length_257_cov_1.194444 | 3.2335 | 257 | 0.7549 | 100.00 | 257 | 4 | 2 | 0.7547 | 3 |
| NODE_463_length_257_cov_1.155556 | 1.7471 | 257 | 0.607 | 100.00 | 257 | 2 | 1 | 0.5991 | 2 |
| NODE_464_length_257_cov_1.105556 | 2.1712 | 257 | 0.7315 | 100.00 | 257 | 2 | 2 | 0.7103 | 2 |
| NODE_465_length_257_cov_1.005556 | 1.7588 | 257 | 0.5875 | 100.00 | 257 | 3 | 0 | 0.5774 | 2 |
| NODE_466_length_257_cov_0.905556 | 1.7549 | 257 | 0.7704 | 100.00 | 257 | 2 | 1 | 0.7561 | 2 |

| | | | | | | | | | |
|---|---|---|---|---|---|---|---|---|---|
| NODE_467_length_257_cov_0.877778 | 1.8132 | 257 | 0.6576 | 100.00 | 257 | 2 | 2 | 0.6803 | 2 |
| NODE_468_length_257_cov_0.811111 | 1.1673 | 257 | 0.6381 | 100.00 | 257 | 1 | 1 | 0.6467 | 1 |
| NODE_469_length_257_cov_0.727778 | 1.1089 | 257 | 0.6848 | 100.00 | 257 | 1 | 1 | 0.6811 | 1 |
| NODE_470_length_257_cov_0.716667 | 1.1518 | 257 | 0.6693 | 100.00 | 257 | 1 | 1 | 0.6722 | 1 |
| NODE_471_length_256_cov_1.413408 | 2.4961 | 256 | 0.6875 | 100.00 | 256 | 3 | 2 | 0.6539 | 2 |
| NODE_472_length_256_cov_1.078212 | 1.7578 | 256 | 0.6875 | 100.00 | 256 | 2 | 1 | 0.6792 | 2 |
| NODE_473_length_256_cov_0.821229 | 1.1758 | 256 | 0.7461 | 100.00 | 256 | 1 | 1 | 0.7475 | 1 |
| NODE_474_length_256_cov_0.821229 | 1.1758 | 256 | 0.6641 | 100.00 | 256 | 2 | 0 | 0.6645 | 1 |
| NODE_475_length_256_cov_0.815642 | 1.1758 | 256 | 0.5938 | 100.00 | 256 | 1 | 1 | 0.6047 | 1 |
| NODE_476_length_256_cov_0.810056 | 1.168 | 256 | 0.707 | 100.00 | 256 | 1 | 1 | 0.6933 | 1 |
| NODE_477_length_256_cov_0.810056 | 1.1719 | 256 | 0.6797 | 100.00 | 256 | 1 | 1 | 0.6833 | 1 |
| NODE_478_length_256_cov_0.759777 | 1.1758 | 256 | 0.5547 | 100.00 | 256 | 1 | 1 | 0.5648 | 1 |
| NODE_479_length_255_cov_1.213483 | 2.1137 | 255 | 0.6667 | 100.00 | 255 | 2 | 2 | 0.6698 | 2 |
| NODE_480_length_255_cov_1.140449 | 2.651 | 255 | 0.651 | 100.00 | 255 | 3 | 2 | 0.6368 | 3 |
| NODE_481_length_255_cov_0.960674 | 1.5765 | 255 | 0.6549 | 100.00 | 255 | 2 | 1 | 0.6493 | 1 |
| NODE_482_length_255_cov_0.837079 | 1.7608 | 255 | 0.7373 | 100.00 | 255 | 2 | 1 | 0.745 | 2 |
| NODE_483_length_255_cov_0.820225 | 1.1765 | 255 | 0.6118 | 100.00 | 255 | 1 | 1 | 0.6067 | 1 |
| NODE_484_length_255_cov_0.814607 | 1.1725 | 255 | 0.4314 | 100.00 | 255 | 1 | 1 | 0.4314 | 1 |
| NODE_485_length_255_cov_0.803371 | 1.1686 | 255 | 0.7176 | 100.00 | 255 | 1 | 1 | 0.7114 | 1 |
| NODE_486_length_255_cov_0.764045 | 1.1804 | 255 | 0.5255 | 100.00 | 255 | 1 | 1 | 0.5249 | 1 |
| NODE_487_length_255_cov_0.752809 | 1.1804 | 255 | 0.5294 | 100.00 | 255 | 1 | 1 | 0.5216 | 1 |
| NODE_488_length_254_cov_2.011299 | 3.5512 | 254 | 0.7362 | 100.00 | 254 | 3 | 3 | 0.7373 | 4 |
| NODE_489_length_254_cov_1.649718 | 2.3622 | 254 | 0.7283 | 100.00 | 254 | 1 | 3 | 0.7317 | 2 |
| NODE_490_length_254_cov_1.604520 | 2.3425 | 254 | 0.7323 | 100.00 | 254 | 2 | 2 | 0.7345 | 3 |
| NODE_491_length_254_cov_1.220339 | 1.7598 | 254 | 0.563 | 100.00 | 254 | 2 | 1 | 0.5727 | 2 |
| NODE_492_length_254_cov_1.135593 | 1.7717 | 254 | 0.6693 | 100.00 | 254 | 0 | 3 | 0.6778 | 2 |
| NODE_493_length_254_cov_1.129944 | 1.7717 | 254 | 0.6772 | 100.00 | 254 | 1 | 2 | 0.68 | 2 |
| NODE_494_length_254_cov_0.830508 | 1.185 | 254 | 0.6969 | 100.00 | 254 | 1 | 1 | 0.6977 | 1 |
| NODE_495_length_254_cov_0.824859 | 1.185 | 254 | 0.6614 | 100.00 | 254 | 1 | 1 | 0.6578 | 1 |
| NODE_496_length_253_cov_1.244318 | 1.7787 | 253 | 0.7391 | 100.00 | 253 | 2 | 1 | 0.7244 | 2 |
| NODE_497_length_253_cov_1.153409 | 1.7866 | 253 | 0.664 | 100.00 | 253 | 2 | 1 | 0.6527 | 2 |
| NODE_498_length_253_cov_0.829545 | 1.1858 | 253 | 0.7036 | 100.00 | 253 | 1 | 1 | 0.6967 | 1 |
| NODE_499_length_253_cov_0.829545 | 1.7787 | 253 | 0.6008 | 100.00 | 253 | 1 | 2 | 0.6289 | 2 |
| NODE_500_length_253_cov_0.823864 | 1.1818 | 253 | 0.6087 | 100.00 | 253 | 1 | 1 | 0.6067 | 1 |
| NODE_501_length_252_cov_0.862857 | 1.7579 | 252 | 0.6508 | 100.00 | 252 | 1 | 2 | 0.6386 | 2 |
| NODE_502_length_252_cov_0.834286 | 1.1984 | 252 | 0.631 | 100.00 | 252 | 1 | 1 | 0.6358 | 1 |
| NODE_503_length_252_cov_0.834286 | 1.1905 | 252 | 0.6587 | 100.00 | 252 | 1 | 1 | 0.65 | 1 |
| NODE_504_length_252_cov_0.800000 | 1.1905 | 252 | 0.4444 | 100.00 | 252 | 1 | 1 | 0.4467 | 1 |
| NODE_505_length_252_cov_0.737143 | 1.1944 | 252 | 0.7341 | 100.00 | 252 | 1 | 1 | 0.7409 | 1 |

| | | | | | | | | | |
|---|---|---|---|---|---|---|---|---|---|
| NODE_506_length_251_cov_1.942529 | 4.9841 | 251 | 0.5697 | 100.00 | 251 | 6 | 4 | 0.534 | 6 |
| NODE_507_length_251_cov_1.270115 | 1.8008 | 251 | 0.6454 | 100.00 | 251 | 1 | 2 | 0.6327 | 2 |
| NODE_508_length_251_cov_1.097701 | 1.988 | 251 | 0.6096 | 100.00 | 251 | 2 | 2 | 0.5972 | 1 |
| NODE_509_length_251_cov_0.844828 | 1.1992 | 251 | 0.6135 | 100.00 | 251 | 1 | 1 | 0.6246 | 1 |
| NODE_510_length_251_cov_0.844828 | 1.1992 | 251 | 0.6733 | 100.00 | 251 | 1 | 1 | 0.6711 | 1 |
| NODE_511_length_251_cov_0.718391 | 2.3904 | 251 | 0.753 | 100.00 | 251 | 2 | 2 | 0.7433 | 2 |
| NODE_512_length_251_cov_0.683908 | 1.1952 | 251 | 0.6494 | 100.00 | 251 | 1 | 1 | 0.66 | 1 |
| NODE_513_length_251_cov_0.614943 | 1.0916 | 251 | 0.5817 | 82.07 | 206 | 1 | 1 | 0.5648 | 1 |
| NODE_514_length_250_cov_1.254335 | 1.796 | 250 | 0.652 | 100.00 | 250 | 2 | 1 | 0.6659 | 2 |
| NODE_515_length_250_cov_1.254335 | 2.392 | 250 | 0.708 | 100.00 | 250 | 2 | 2 | 0.709 | 2 |
| NODE_516_length_250_cov_1.236994 | 2.392 | 250 | 0.752 | 100.00 | 250 | 1 | 3 | 0.7262 | 3 |
| NODE_517_length_250_cov_0.849711 | 1.204 | 250 | 0.74 | 100.00 | 250 | 1 | 1 | 0.7475 | 1 |
| NODE_518_length_250_cov_0.803468 | 1.684 | 250 | 0.724 | 100.00 | 250 | 2 | 2 | 0.7173 | 2 |
| NODE_519_length_250_cov_0.716763 | 1.112 | 250 | 0.68 | 100.00 | 250 | 0 | 2 | 0.691 | 1 |
| NODE_520_length_250_cov_0.710983 | 1.196 | 250 | 0.696 | 100.00 | 250 | 1 | 1 | 0.7023 | 1 |
| NODE_521_length_250_cov_0.682081 | 1.192 | 250 | 0.676 | 100.00 | 250 | 1 | 1 | 0.6867 | 1 |
| NODE_522_length_249_cov_0.970930 | 2.1767 | 249 | 0.6225 | 100.00 | 249 | 2 | 2 | 0.6256 | 2 |
| NODE_523_length_249_cov_0.848837 | 1.2048 | 249 | 0.4659 | 100.00 | 249 | 1 | 1 | 0.4733 | 1 |
| NODE_524_length_249_cov_0.848837 | 1.2048 | 249 | 0.6827 | 100.00 | 249 | 1 | 1 | 0.6767 | 1 |
| NODE_525_length_249_cov_0.848837 | 1.2048 | 249 | 0.6225 | 100.00 | 249 | 1 | 1 | 0.6167 | 1 |
| NODE_526_length_249_cov_0.848837 | 1.2088 | 249 | 0.6225 | 100.00 | 249 | 1 | 1 | 0.6379 | 1 |
| NODE_527_length_249_cov_0.750000 | 2.2851 | 249 | 0.6948 | 100.00 | 249 | 2 | 4 | 0.6432 | 2 |
| NODE_528_length_248_cov_0.941520 | 1.8105 | 248 | 0.7137 | 100.00 | 248 | 1 | 2 | 0.7016 | 2 |
| NODE_529_length_248_cov_0.923977 | 1.9476 | 248 | 0.6008 | 100.00 | 248 | 3 | 2 | 0.5942 | 2 |
| NODE_530_length_248_cov_0.859649 | 1.2137 | 248 | 0.7339 | 100.00 | 248 | 1 | 1 | 0.7409 | 1 |
| NODE_531_length_248_cov_0.859649 | 1.2137 | 248 | 0.6129 | 100.00 | 248 | 1 | 1 | 0.598 | 1 |
| NODE_532_length_248_cov_0.859649 | 1.2137 | 248 | 0.6331 | 100.00 | 248 | 1 | 1 | 0.6246 | 1 |
| NODE_533_length_248_cov_0.853801 | 1.7984 | 248 | 0.629 | 100.00 | 248 | 1 | 2 | 0.6173 | 2 |
| NODE_534_length_248_cov_0.842105 | 1.2056 | 248 | 0.6492 | 100.00 | 248 | 1 | 1 | 0.6622 | 1 |
| NODE_535_length_248_cov_0.836257 | 1.1976 | 248 | 0.7137 | 100.00 | 248 | 1 | 1 | 0.7104 | 1 |
| NODE_536_length_248_cov_0.795322 | 1.7177 | 248 | 0.7258 | 100.00 | 248 | 2 | 1 | 0.7416 | 2 |
| NODE_537_length_248_cov_0.695906 | 2.4194 | 248 | 0.6976 | 100.00 | 248 | 1 | 3 | 0.715 | 3 |
| NODE_538_length_248_cov_0.672515 | 1.2137 | 248 | 0.7177 | 100.00 | 248 | 1 | 1 | 0.7309 | 1 |
| NODE_539_length_247_cov_2.147059 | 3.6032 | 247 | 0.5263 | 100.00 | 247 | 4 | 3 | 0.5258 | 4 |
| NODE_540_length_247_cov_1.723529 | 2.7733 | 247 | 0.7126 | 100.00 | 247 | 3 | 2 | 0.7007 | 3 |
| NODE_541_length_247_cov_1.288235 | 1.8219 | 247 | 0.668 | 100.00 | 247 | 2 | 1 | 0.6778 | 2 |
| NODE_542_length_247_cov_0.935294 | 1.8219 | 247 | 0.6721 | 100.00 | 247 | 0 | 3 | 0.6578 | 2 |
| NODE_543_length_247_cov_0.870588 | 1.2227 | 247 | 0.6518 | 100.00 | 247 | 1 | 1 | 0.6623 | 1 |
| NODE_544_length_247_cov_0.858824 | 1.7247 | 247 | 0.6356 | 100.00 | 247 | 2 | 2 | 0.6244 | 1 |

| | | | | | | | | | |
|---|---|---|---|---|---|---|---|---|---|
| NODE_545_length_247_cov_0.858824 | 1.8259 | 247 | 0.664 | 100.00 | 247 | 1 | 2 | 0.6652 | 2 |
| NODE_546_length_247_cov_0.852941 | 1.2105 | 247 | 0.7085 | 100.00 | 247 | 1 | 1 | 0.7124 | 1 |
| NODE_547_length_247_cov_0.805882 | 1.2146 | 247 | 0.6923 | 100.00 | 247 | 2 | 0 | 0.7 | 1 |
| NODE_548_length_247_cov_0.711765 | 1.2146 | 247 | 0.7045 | 100.00 | 247 | 1 | 1 | 0.6987 | 1 |
| NODE_549_length_246_cov_2.609467 | 3.6748 | 246 | 0.6382 | 100.00 | 246 | 3 | 3 | 0.6294 | 5 |
| NODE_550_length_246_cov_1.715976 | 2.4309 | 246 | 0.6382 | 100.00 | 246 | 2 | 2 | 0.6455 | 2 |
| NODE_551_length_246_cov_1.568047 | 2.9675 | 246 | 0.5122 | 100.00 | 246 | 3 | 3 | 0.5096 | 2 |
| NODE_552_length_246_cov_1.106509 | 3.4512 | 246 | 0.561 | 100.00 | 246 | 4 | 3 | 0.5441 | 4 |
| NODE_553_length_246_cov_1.076923 | 1.6789 | 246 | 0.7154 | 100.00 | 246 | 2 | 1 | 0.7264 | 2 |
| NODE_554_length_246_cov_0.923077 | 1.8211 | 246 | 0.6463 | 100.00 | 246 | 2 | 1 | 0.6169 | 2 |
| NODE_555_length_246_cov_0.875740 | 1.2276 | 246 | 0.6463 | 100.00 | 246 | 1 | 1 | 0.6358 | 1 |
| NODE_556_length_246_cov_0.869822 | 1.2236 | 246 | 0.7276 | 100.00 | 246 | 1 | 1 | 0.7276 | 1 |
| NODE_557_length_246_cov_0.863905 | 1.2195 | 246 | 0.687 | 100.00 | 246 | 1 | 1 | 0.6833 | 1 |
| NODE_558_length_246_cov_0.857988 | 1.2154 | 246 | 0.687 | 100.00 | 246 | 1 | 1 | 0.68 | 1 |
| NODE_559_length_246_cov_0.798817 | 1.2195 | 246 | 0.6179 | 100.00 | 246 | 1 | 1 | 0.6367 | 1 |
| NODE_560_length_246_cov_0.686391 | 1.2154 | 246 | 0.6504 | 100.00 | 246 | 1 | 1 | 0.6789 | 1 |
| NODE_561_length_246_cov_0.674556 | 1.565 | 246 | 0.6829 | 100.00 | 246 | 2 | 1 | 0.6831 | 1 |
| NODE_562_length_245_cov_1.601190 | 2.3551 | 245 | 0.6531 | 100.00 | 245 | 2 | 2 | 0.6638 | 3 |
| NODE_563_length_245_cov_0.875000 | 1.2286 | 245 | 0.7184 | 100.00 | 245 | 1 | 1 | 0.7176 | 1 |
| NODE_564_length_245_cov_0.875000 | 1.2286 | 245 | 0.7388 | 100.00 | 245 | 1 | 1 | 0.7442 | 1 |
| NODE_565_length_245_cov_0.875000 | 1.2286 | 245 | 0.6898 | 100.00 | 245 | 1 | 1 | 0.6744 | 1 |
| NODE_566_length_245_cov_0.869048 | 1.2245 | 245 | 0.6694 | 100.00 | 245 | 1 | 1 | 0.6667 | 1 |
| NODE_567_length_245_cov_0.755952 | 3.4571 | 245 | 0.5633 | 100.00 | 245 | 4 | 5 | 0.5177 | 3 |
| NODE_568_length_244_cov_1.305389 | 1.8484 | 244 | 0.6639 | 100.00 | 244 | 2 | 1 | 0.663 | 2 |
| NODE_569_length_244_cov_1.000000 | 1.9467 | 244 | 0.5492 | 100.00 | 244 | 2 | 2 | 0.5315 | 1 |
| NODE_570_length_244_cov_0.880240 | 1.2336 | 244 | 0.5738 | 100.00 | 244 | 1 | 1 | 0.5615 | 1 |
| NODE_571_length_244_cov_0.880240 | 1.8484 | 244 | 0.6844 | 100.00 | 244 | 2 | 2 | 0.7029 | 1 |
| NODE_572_length_244_cov_0.868263 | 1.2295 | 244 | 0.6885 | 100.00 | 244 | 1 | 1 | 0.6867 | 1 |
| NODE_573_length_244_cov_0.856287 | 1.2213 | 244 | 0.709 | 100.00 | 244 | 0 | 2 | 0.702 | 1 |
| NODE_574_length_243_cov_1.319277 | 1.856 | 243 | 0.6872 | 100.00 | 243 | 1 | 2 | 0.6918 | 2 |
| NODE_575_length_243_cov_1.234940 | 1.8519 | 243 | 0.5103 | 100.00 | 243 | 2 | 1 | 0.5 | 2 |
| NODE_576_length_243_cov_0.879518 | 1.2387 | 243 | 0.7078 | 100.00 | 243 | 1 | 1 | 0.7143 | 1 |
| NODE_577_length_243_cov_0.873494 | 1.2305 | 243 | 0.6626 | 100.00 | 243 | 2 | 0 | 0.6622 | 1 |
| NODE_578_length_243_cov_0.867470 | 1.2346 | 243 | 0.6337 | 100.00 | 243 | 1 | 1 | 0.6167 | 1 |
| NODE_579_length_243_cov_0.825301 | 1.8189 | 243 | 0.6831 | 100.00 | 243 | 2 | 1 | 0.694 | 2 |
| NODE_580_length_243_cov_0.759036 | 1.2387 | 243 | 0.6872 | 100.00 | 243 | 1 | 1 | 0.6811 | 1 |
| NODE_581_length_242_cov_2.515152 | 4.6736 | 242 | 0.6942 | 100.00 | 242 | 4 | 4 | 0.7064 | 4 |
| NODE_582_length_242_cov_1.824242 | 3.0992 | 242 | 0.6612 | 100.00 | 242 | 2 | 3 | 0.6853 | 4 |
| NODE_583_length_242_cov_0.896970 | 1.2479 | 242 | 0.657 | 100.00 | 242 | 1 | 1 | 0.649 | 1 |

| | | | | | | | | | |
|---|---|---|---|---|---|---|---|---|---|
| NODE_584_length_242_cov_0.890909 | 1.2438 | 242 | 0.686 | 100.00 | 242 | 1 | 1 | 0.6711 | 1 |
| NODE_585_length_242_cov_0.890909 | 1.2438 | 242 | 0.6653 | 100.00 | 242 | 2 | 0 | 0.6578 | 1 |
| NODE_586_length_242_cov_0.884848 | 1.2397 | 242 | 0.6901 | 100.00 | 242 | 1 | 1 | 0.6854 | 1 |
| NODE_587_length_242_cov_0.884848 | 1.2397 | 242 | 0.5496 | 100.00 | 242 | 1 | 1 | 0.5767 | 1 |
| NODE_588_length_242_cov_0.884848 | 1.2397 | 242 | 0.6653 | 100.00 | 242 | 1 | 1 | 0.67 | 1 |
| NODE_589_length_242_cov_0.884848 | 1.2397 | 242 | 0.6777 | 100.00 | 242 | 1 | 1 | 0.6733 | 1 |
| NODE_590_length_242_cov_0.878788 | 1.2397 | 242 | 0.7355 | 100.00 | 242 | 1 | 1 | 0.7467 | 1 |
| NODE_591_length_242_cov_0.878788 | 1.2355 | 242 | 0.7107 | 100.00 | 242 | 1 | 1 | 0.6957 | 1 |
| NODE_592_length_242_cov_0.878788 | 1.2355 | 242 | 0.6405 | 100.00 | 242 | 1 | 1 | 0.6421 | 1 |
| NODE_593_length_242_cov_0.769697 | 1.2438 | 242 | 0.6529 | 100.00 | 242 | 1 | 1 | 0.6545 | 1 |
| NODE_594_length_241_cov_2.402439 | 4.278 | 241 | 0.6846 | 100.00 | 241 | 4 | 3 | 0.6825 | 4 |
| NODE_595_length_241_cov_1.792683 | 2.4979 | 241 | 0.7054 | 100.00 | 241 | 2 | 2 | 0.7043 | 2 |
| NODE_596_length_241_cov_1.774390 | 2.4855 | 241 | 0.527 | 100.00 | 241 | 2 | 2 | 0.5426 | 3 |
| NODE_597_length_241_cov_1.067073 | 1.8631 | 241 | 0.7261 | 100.00 | 241 | 1 | 2 | 0.7327 | 2 |
| NODE_598_length_241_cov_0.896341 | 1.2531 | 241 | 0.7469 | 100.00 | 241 | 1 | 1 | 0.7483 | 1 |
| NODE_599_length_241_cov_0.890244 | 1.2448 | 241 | 0.668 | 100.00 | 241 | 1 | 1 | 0.6567 | 1 |
| NODE_600_length_241_cov_0.859756 | 1.249 | 241 | 0.4689 | 100.00 | 241 | 0 | 2 | 0.4784 | 1 |
| NODE_601_length_241_cov_0.658537 | 1.0871 | 241 | 0.668 | 100.00 | 241 | 1 | 1 | 0.6523 | 1 |
| NODE_602_length_240_cov_2.693252 | 3.7583 | 240 | 0.725 | 100.00 | 240 | 2 | 4 | 0.7184 | 3 |
| NODE_603_length_240_cov_2.312883 | 4.075 | 240 | 0.5125 | 100.00 | 240 | 4 | 4 | 0.5297 | 4 |
| NODE_604_length_240_cov_1.773006 | 2.8417 | 240 | 0.7083 | 100.00 | 240 | 2 | 3 | 0.6946 | 3 |
| NODE_605_length_240_cov_1.361963 | 1.8875 | 240 | 0.7125 | 100.00 | 240 | 1 | 2 | 0.7086 | 2 |
| NODE_606_length_240_cov_1.012270 | 2.5083 | 240 | 0.6917 | 100.00 | 240 | 3 | 1 | 0.6927 | 3 |
| NODE_607_length_240_cov_0.895706 | 1.25 | 240 | 0.6625 | 100.00 | 240 | 1 | 1 | 0.6633 | 1 |
| NODE_608_length_240_cov_0.889571 | 1.875 | 240 | 0.7167 | 100.00 | 240 | 1 | 2 | 0.7244 | 2 |
| NODE_609_length_240_cov_0.815951 | 1.25 | 240 | 0.6417 | 100.00 | 240 | 1 | 1 | 0.6333 | 1 |
| NODE_610_length_240_cov_0.674847 | 1.1 | 240 | 0.6 | 100.00 | 240 | 1 | 1 | 0.6146 | 1 |
| NODE_611_length_239_cov_1.351852 | 1.887 | 239 | 0.6653 | 100.00 | 239 | 1 | 2 | 0.6652 | 2 |
| NODE_612_length_239_cov_1.296296 | 2.431 | 239 | 0.6402 | 98.33 | 235 | 3 | 1 | 0.6556 | 2 |
| NODE_613_length_239_cov_0.913580 | 1.2636 | 239 | 0.6067 | 100.00 | 239 | 1 | 1 | 0.6225 | 1 |
| NODE_614_length_239_cov_0.907407 | 1.2594 | 239 | 0.6695 | 100.00 | 239 | 1 | 1 | 0.6877 | 1 |
| NODE_615_length_239_cov_0.901235 | 1.2552 | 239 | 0.6192 | 100.00 | 239 | 1 | 1 | 0.6133 | 1 |
| NODE_616_length_239_cov_0.901235 | 1.2552 | 239 | 0.4226 | 100.00 | 239 | 1 | 1 | 0.4367 | 1 |
| NODE_617_length_239_cov_0.820988 | 1.2008 | 239 | 0.6862 | 100.00 | 239 | 1 | 1 | 0.6867 | 1 |
| NODE_618_length_239_cov_0.759259 | 1.7866 | 239 | 0.59 | 100.00 | 239 | 2 | 2 | 0.5765 | 1 |
| NODE_619_length_239_cov_0.703704 | 1.2594 | 239 | 0.6695 | 100.00 | 239 | 1 | 1 | 0.6645 | 1 |
| NODE_620_length_238_cov_1.658385 | 2.521 | 238 | 0.6429 | 100.00 | 238 | 2 | 2 | 0.6467 | 2 |
| NODE_621_length_238_cov_1.000000 | 3.7101 | 238 | 0.6723 | 100.00 | 238 | 3 | 3 | 0.6618 | 4 |
| NODE_622_length_238_cov_0.913043 | 1.2647 | 238 | 0.605 | 100.00 | 238 | 1 | 1 | 0.6047 | 1 |

| | | | | | | | | | |
|---|---|---|---|---|---|---|---|---|---|
| NODE_623_length_238_cov_0.913043 | 1.2647 | 238 | 0.6471 | 100.00 | 238 | 1 | 1 | 0.6379 | 1 |
| NODE_624_length_238_cov_0.913043 | 1.2647 | 238 | 0.6471 | 100.00 | 238 | 2 | 0 | 0.6645 | 1 |
| NODE_625_length_238_cov_0.894410 | 1.2563 | 238 | 0.7311 | 100.00 | 238 | 2 | 0 | 0.7342 | 1 |
| NODE_626_length_238_cov_0.888199 | 1.5714 | 238 | 0.4832 | 100.00 | 238 | 2 | 1 | 0.4866 | 1 |
| NODE_627_length_238_cov_0.850932 | 1.2605 | 238 | 0.605 | 100.00 | 238 | 0 | 2 | 0.5933 | 1 |
| NODE_628_length_238_cov_0.838509 | 1.2605 | 238 | 0.6681 | 100.00 | 238 | 1 | 1 | 0.6633 | 1 |
| NODE_629_length_238_cov_0.763975 | 1.1639 | 238 | 0.6807 | 100.00 | 238 | 1 | 1 | 0.6957 | 1 |
| NODE_630_length_237_cov_1.368750 | 1.903 | 237 | 0.5738 | 100.00 | 237 | 0 | 3 | 0.5787 | 2 |
| NODE_631_length_237_cov_1.106250 | 1.8017 | 237 | 0.6456 | 100.00 | 237 | 1 | 2 | 0.6464 | 2 |
| NODE_632_length_237_cov_0.968750 | 1.903 | 237 | 0.6203 | 100.00 | 237 | 1 | 2 | 0.5987 | 2 |
| NODE_633_length_237_cov_0.918750 | 1.27 | 237 | 0.6835 | 100.00 | 237 | 1 | 1 | 0.6811 | 1 |
| NODE_634_length_237_cov_0.912500 | 1.2658 | 237 | 0.6709 | 100.00 | 237 | 1 | 1 | 0.68 | 1 |
| NODE_635_length_237_cov_0.912500 | 1.2658 | 237 | 0.5823 | 100.00 | 237 | 1 | 1 | 0.5967 | 1 |
| NODE_636_length_237_cov_0.906250 | 1.2616 | 237 | 0.4473 | 100.00 | 237 | 1 | 1 | 0.4415 | 1 |
| NODE_637_length_237_cov_0.906250 | 1.7342 | 237 | 0.6709 | 100.00 | 237 | 2 | 2 | 0.6496 | 1 |
| NODE_638_length_237_cov_0.875000 | 1.2743 | 237 | 0.6962 | 100.00 | 237 | 1 | 1 | 0.7053 | 1 |
| NODE_639_length_237_cov_0.731250 | 1.2616 | 237 | 0.7046 | 100.00 | 237 | 2 | 0 | 0.7143 | 1 |
| NODE_640_length_236_cov_2.754717 | 3.8178 | 236 | 0.6483 | 100.00 | 236 | 2 | 4 | 0.6204 | 4 |
| NODE_641_length_236_cov_1.817610 | 2.5339 | 236 | 0.7246 | 100.00 | 236 | 2 | 2 | 0.7224 | 2 |
| NODE_642_length_236_cov_1.515723 | 2.5508 | 236 | 0.6992 | 100.00 | 236 | 2 | 2 | 0.701 | 2 |
| NODE_643_length_236_cov_1.364780 | 2.5339 | 236 | 0.7076 | 100.00 | 236 | 2 | 2 | 0.7057 | 2 |
| NODE_644_length_236_cov_0.924528 | 1.2754 | 236 | 0.6737 | 100.00 | 236 | 1 | 1 | 0.6744 | 1 |
| NODE_645_length_236_cov_0.924528 | 1.2754 | 236 | 0.589 | 100.00 | 236 | 1 | 1 | 0.6179 | 1 |
| NODE_646_length_236_cov_0.918239 | 1.9068 | 236 | 0.7161 | 100.00 | 236 | 2 | 1 | 0.7311 | 2 |
| NODE_647_length_236_cov_0.918239 | 1.2797 | 236 | 0.6314 | 100.00 | 236 | 1 | 1 | 0.6325 | 1 |
| NODE_648_length_236_cov_0.918239 | 1.2712 | 236 | 0.7373 | 100.00 | 236 | 1 | 1 | 0.73 | 1 |
| NODE_649_length_236_cov_0.918239 | 1.2712 | 236 | 0.6949 | 100.00 | 236 | 1 | 1 | 0.69 | 1 |
| NODE_650_length_236_cov_0.918239 | 1.2712 | 236 | 0.5593 | 100.00 | 236 | 1 | 1 | 0.58 | 1 |
| NODE_651_length_236_cov_0.911950 | 1.2669 | 236 | 0.6737 | 100.00 | 236 | 1 | 1 | 0.6622 | 1 |
| NODE_652_length_236_cov_0.899371 | 1.2585 | 236 | 0.6568 | 100.00 | 236 | 0 | 2 | 0.6412 | 1 |
| NODE_653_length_236_cov_0.805031 | 1.2754 | 236 | 0.5508 | 100.00 | 236 | 1 | 1 | 0.5382 | 1 |
| NODE_654_length_236_cov_0.761006 | 1.2712 | 236 | 0.5508 | 100.00 | 236 | 2 | 0 | 0.56 | 1 |
| NODE_655_length_235_cov_1.493671 | 2.8511 | 235 | 0.6426 | 100.00 | 235 | 2 | 3 | 0.6388 | 2 |
| NODE_656_length_235_cov_1.386076 | 2.5617 | 235 | 0.4638 | 100.00 | 235 | 2 | 2 | 0.4319 | 3 |
| NODE_657_length_235_cov_1.379747 | 2.4213 | 235 | 0.5191 | 100.00 | 235 | 2 | 2 | 0.5132 | 2 |
| NODE_658_length_235_cov_1.107595 | 1.9149 | 235 | 0.7447 | 100.00 | 235 | 2 | 1 | 0.7444 | 2 |
| NODE_659_length_235_cov_0.981013 | 1.9277 | 235 | 0.7702 | 100.00 | 235 | 2 | 1 | 0.7638 | 2 |
| NODE_660_length_235_cov_0.930380 | 1.2809 | 235 | 0.7106 | 100.00 | 235 | 1 | 1 | 0.701 | 1 |
| NODE_661_length_235_cov_0.930380 | 1.9106 | 235 | 0.5787 | 100.00 | 235 | 1 | 2 | 0.5857 | 2 |

| | | | | | | | | | |
|---|---|---|---|---|---|---|---|---|---|
| NODE_662_length_235_cov_0.930380 | 1.2851 | 235 | 0.617 | 100.00 | 235 | 1 | 1 | 0.6225 | 1 |
| NODE_663_length_235_cov_0.892405 | 1.2553 | 235 | 0.6809 | 100.00 | 235 | 1 | 1 | 0.6611 | 1 |
| NODE_664_length_235_cov_0.873418 | 1.2766 | 235 | 0.6638 | 100.00 | 235 | 1 | 1 | 0.66 | 1 |
| NODE_665_length_234_cov_1.821656 | 2.5769 | 234 | 0.5983 | 100.00 | 234 | 2 | 2 | 0.6103 | 3 |
| NODE_666_length_234_cov_1.401274 | 1.9274 | 234 | 0.7564 | 100.00 | 234 | 2 | 1 | 0.7472 | 2 |
| NODE_667_length_234_cov_1.388535 | 1.9188 | 234 | 0.7051 | 100.00 | 234 | 1 | 2 | 0.6949 | 2 |
| NODE_668_length_234_cov_1.375796 | 1.9188 | 234 | 0.6368 | 100.00 | 234 | 2 | 1 | 0.6281 | 2 |
| NODE_669_length_234_cov_1.108280 | 1.7308 | 234 | 0.688 | 100.00 | 234 | 2 | 1 | 0.6716 | 2 |
| NODE_670_length_234_cov_1.063694 | 2.0299 | 234 | 0.6667 | 100.00 | 234 | 2 | 2 | 0.6568 | 1 |
| NODE_671_length_234_cov_1.038217 | 1.6838 | 234 | 0.6496 | 100.00 | 234 | 2 | 1 | 0.632 | 2 |
| NODE_672_length_234_cov_1.019108 | 1.9316 | 234 | 0.7179 | 100.00 | 234 | 2 | 1 | 0.7235 | 2 |
| NODE_673_length_234_cov_0.936306 | 1.2863 | 234 | 0.7051 | 100.00 | 234 | 2 | 0 | 0.7043 | 1 |
| NODE_674_length_234_cov_0.929936 | 1.2821 | 234 | 0.7479 | 100.00 | 234 | 2 | 0 | 0.7633 | 1 |
| NODE_675_length_234_cov_0.910828 | 1.2692 | 234 | 0.6538 | 100.00 | 234 | 1 | 1 | 0.6611 | 1 |
| NODE_676_length_234_cov_0.834395 | 1.2821 | 234 | 0.6581 | 100.00 | 234 | 1 | 1 | 0.6433 | 1 |
| NODE_677_length_234_cov_0.789809 | 1.9231 | 234 | 0.7393 | 100.00 | 234 | 1 | 2 | 0.7378 | 2 |
| NODE_678_length_234_cov_0.707006 | 1.1624 | 234 | 0.6197 | 100.00 | 234 | 1 | 1 | 0.6379 | 1 |
| NODE_679_length_233_cov_1.435897 | 2.5923 | 233 | 0.4464 | 100.00 | 233 | 2 | 2 | 0.4487 | 2 |
| NODE_680_length_233_cov_1.378205 | 1.9313 | 233 | 0.6352 | 100.00 | 233 | 2 | 1 | 0.6311 | 2 |
| NODE_681_length_233_cov_1.358974 | 1.9356 | 233 | 0.7082 | 100.00 | 233 | 1 | 2 | 0.7058 | 2 |
| NODE_682_length_233_cov_1.012821 | 1.9313 | 233 | 0.515 | 100.00 | 233 | 2 | 1 | 0.5178 | 2 |
| NODE_683_length_233_cov_1.000000 | 1.9399 | 233 | 0.5966 | 100.00 | 233 | 1 | 2 | 0.5907 | 2 |
| NODE_684_length_233_cov_0.948718 | 1.9142 | 233 | 0.7082 | 100.00 | 233 | 2 | 1 | 0.7152 | 2 |
| NODE_685_length_233_cov_0.942308 | 1.2918 | 233 | 0.6009 | 100.00 | 233 | 1 | 1 | 0.6179 | 1 |
| NODE_686_length_233_cov_0.935897 | 1.2918 | 233 | 0.6824 | 100.00 | 233 | 1 | 1 | 0.6944 | 1 |
| NODE_687_length_233_cov_0.929487 | 1.2833 | 233 | 0.6524 | 100.00 | 233 | 1 | 1 | 0.6589 | 1 |
| NODE_688_length_233_cov_0.743590 | 1.2918 | 233 | 0.6438 | 76.82 | 179 | 2 | 0 | 0.6146 | 2 |
| NODE_689_length_232_cov_1.374194 | 1.9483 | 232 | 0.7112 | 100.00 | 232 | 1 | 2 | 0.6969 | 2 |
| NODE_690_length_232_cov_0.948387 | 1.2974 | 232 | 0.6164 | 100.00 | 232 | 1 | 1 | 0.6113 | 1 |
| NODE_691_length_232_cov_0.948387 | 1.931 | 232 | 0.625 | 100.00 | 232 | 1 | 2 | 0.6295 | 2 |
| NODE_692_length_232_cov_0.864516 | 1.2414 | 232 | 0.7414 | 100.00 | 232 | 1 | 1 | 0.7425 | 1 |
| NODE_693_length_232_cov_0.851613 | 1.5647 | 232 | 0.5172 | 100.00 | 232 | 2 | 1 | 0.5124 | 1 |
| NODE_694_length_232_cov_0.761290 | 1.2974 | 232 | 0.6379 | 100.00 | 232 | 2 | 0 | 0.6312 | 1 |
| NODE_695_length_232_cov_0.625806 | 1.7198 | 232 | 0.6897 | 100.00 | 232 | 3 | 1 | 0.6943 | 2 |
| NODE_696_length_231_cov_2.811688 | 3.8788 | 231 | 0.6623 | 100.00 | 231 | 3 | 3 | 0.6652 | 3 |
| NODE_697_length_231_cov_1.428571 | 2.368 | 231 | 0.632 | 100.00 | 231 | 2 | 2 | 0.6317 | 2 |
| NODE_698_length_231_cov_1.383117 | 2.5931 | 231 | 0.6926 | 100.00 | 231 | 2 | 2 | 0.6828 | 3 |
| NODE_699_length_231_cov_0.954545 | 1.303 | 231 | 0.6537 | 100.00 | 231 | 1 | 1 | 0.6645 | 1 |
| NODE_700_length_231_cov_0.954545 | 1.303 | 231 | 0.684 | 100.00 | 231 | 0 | 2 | 0.7043 | 1 |

| | | | | | | | | | |
|---|---|---|---|---|---|---|---|---|---|
| NODE_701_length_231_cov_0.948052 | 1.2987 | 231 | 0.6667 | 100.00 | 231 | 1 | 1 | 0.6667 | 1 |
| NODE_702_length_231_cov_0.948052 | 1.303 | 231 | 0.6234 | 91.34 | 211 | 1 | 1 | 0.6246 | 1 |
| NODE_703_length_231_cov_0.948052 | 1.2987 | 231 | 0.6407 | 100.00 | 231 | 1 | 1 | 0.6367 | 1 |
| NODE_704_length_231_cov_0.948052 | 1.71 | 231 | 0.671 | 100.00 | 231 | 2 | 1 | 0.6541 | 2 |
| NODE_705_length_231_cov_0.883117 | 1.3074 | 231 | 0.7229 | 100.00 | 231 | 1 | 1 | 0.7285 | 1 |
| NODE_706_length_231_cov_0.798701 | 1.1991 | 231 | 0.6407 | 100.00 | 231 | 1 | 1 | 0.6379 | 1 |
| NODE_707_length_231_cov_0.727273 | 1.303 | 231 | 0.7186 | 100.00 | 231 | 2 | 0 | 0.7243 | 1 |
| NODE_708_length_231_cov_0.662338 | 1.2987 | 231 | 0.632 | 100.00 | 231 | 1 | 1 | 0.6233 | 1 |
| NODE_709_length_230_cov_1.411765 | 1.9522 | 230 | 0.7217 | 100.00 | 230 | 2 | 1 | 0.7238 | 2 |
| NODE_710_length_230_cov_1.320261 | 2.2174 | 230 | 0.5174 | 100.00 | 230 | 2 | 2 | 0.5216 | 2 |
| NODE_711_length_230_cov_1.150327 | 2.5304 | 230 | 0.6174 | 100.00 | 230 | 3 | 3 | 0.6254 | 3 |
| NODE_712_length_230_cov_0.986928 | 1.6652 | 230 | 0.7 | 100.00 | 230 | 2 | 1 | 0.7102 | 2 |
| NODE_713_length_230_cov_0.967320 | 1.313 | 230 | 0.7217 | 100.00 | 230 | 1 | 1 | 0.7252 | 1 |
| NODE_714_length_230_cov_0.960784 | 1.9304 | 230 | 0.6957 | 100.00 | 230 | 2 | 2 | 0.704 | 2 |
| NODE_715_length_230_cov_0.960784 | 1.3087 | 230 | 0.6087 | 100.00 | 230 | 1 | 1 | 0.598 | 1 |
| NODE_716_length_230_cov_0.954248 | 1.3043 | 230 | 0.5435 | 100.00 | 230 | 1 | 1 | 0.5567 | 1 |
| NODE_717_length_230_cov_0.954248 | 1.3043 | 230 | 0.6565 | 100.00 | 230 | 1 | 1 | 0.6567 | 1 |
| NODE_718_length_230_cov_0.947712 | 2.6087 | 230 | 0.687 | 100.00 | 230 | 2 | 2 | 0.6817 | 2 |
| NODE_719_length_230_cov_0.810458 | 1.3087 | 230 | 0.6565 | 100.00 | 230 | 1 | 1 | 0.6512 | 1 |
| NODE_720_length_230_cov_0.803922 | 1.313 | 230 | 0.7 | 100.00 | 230 | 1 | 1 | 0.6987 | 1 |
| NODE_721_length_230_cov_0.803922 | 1.313 | 230 | 0.7348 | 100.00 | 230 | 1 | 1 | 0.7318 | 1 |
| NODE_722_length_230_cov_0.673203 | 1.8087 | 230 | 0.6348 | 100.00 | 230 | 1 | 2 | 0.6133 | 2 |
| NODE_723_length_230_cov_0.660131 | 1.313 | 230 | 0.6435 | 100.00 | 230 | 2 | 0 | 0.6391 | 1 |
| NODE_724_length_229_cov_2.052632 | 3.5895 | 229 | 0.6987 | 100.00 | 229 | 2 | 4 | 0.6694 | 4 |
| NODE_725_length_229_cov_1.440789 | 1.9694 | 229 | 0.6987 | 100.00 | 229 | 2 | 1 | 0.7162 | 2 |
| NODE_726_length_229_cov_1.434211 | 3.441 | 229 | 0.5066 | 100.00 | 229 | 3 | 3 | 0.4962 | 2 |
| NODE_727_length_229_cov_0.973684 | 1.3188 | 229 | 0.4847 | 100.00 | 229 | 1 | 1 | 0.4768 | 1 |
| NODE_728_length_229_cov_0.967105 | 1.3144 | 229 | 0.7074 | 100.00 | 229 | 0 | 2 | 0.6944 | 1 |
| NODE_729_length_229_cov_0.967105 | 1.3144 | 229 | 0.6943 | 100.00 | 229 | 1 | 1 | 0.7043 | 1 |
| NODE_730_length_229_cov_0.960526 | 1.31 | 229 | 0.6507 | 100.00 | 229 | 1 | 1 | 0.64 | 1 |
| NODE_731_length_229_cov_0.960526 | 1.31 | 229 | 0.6507 | 100.00 | 229 | 1 | 1 | 0.6333 | 1 |
| NODE_732_length_229_cov_0.960526 | 1.3144 | 229 | 0.4847 | 100.00 | 229 | 1 | 1 | 0.5183 | 1 |
| NODE_733_length_229_cov_0.960526 | 1.3144 | 229 | 0.559 | 100.00 | 229 | 1 | 1 | 0.5581 | 1 |
| NODE_734_length_229_cov_0.960526 | 1.3144 | 229 | 0.6114 | 100.00 | 229 | 1 | 1 | 0.6246 | 1 |
| NODE_735_length_229_cov_0.960526 | 1.31 | 229 | 0.7118 | 100.00 | 229 | 1 | 1 | 0.7167 | 1 |
| NODE_736_length_229_cov_0.960526 | 1.3144 | 229 | 0.6376 | 100.00 | 229 | 1 | 1 | 0.6478 | 1 |
| NODE_737_length_229_cov_0.960526 | 1.31 | 229 | 0.7249 | 100.00 | 229 | 1 | 1 | 0.7233 | 1 |
| NODE_738_length_229_cov_0.875000 | 1.31 | 229 | 0.7424 | 100.00 | 229 | 1 | 1 | 0.7367 | 1 |
| NODE_739_length_229_cov_0.828947 | 1.5721 | 229 | 0.6987 | 100.00 | 229 | 2 | 1 | 0.6944 | 1 |

| | | | | | | | | | |
|---|---|---|---|---|---|---|---|---|---|
| NODE_740_length_229_cov_0.828947 | 1.3144 | 229 | 0.7249 | 100.00 | 229 | 1 | 1 | 0.7276 | 1 |
| NODE_741_length_229_cov_0.782895 | 1.3144 | 229 | 0.7249 | 100.00 | 229 | 1 | 1 | 0.7209 | 1 |
| NODE_742_length_229_cov_0.717105 | 1.31 | 229 | 0.6507 | 100.00 | 229 | 2 | 0 | 0.65 | 1 |
| NODE_743_length_78_cov_9798.000000 | 0 | 78 | 1 | 0.00 | 0 | 0 | 0 | 0 | 0 |

| Std_Dev |
|---|
| 1901.25 |
| 7.21 |
| 2.43 |
| 2.31 |
| 1.61 |
| 1.25 |
| 8.72 |
| 2.7 |
| 1.55 |
| 0.89 |
| 1.37 |
| 0.62 |
| 1.68 |
| 0.94 |
| 0.69 |
| 1.31 |
| 0.95 |
| 0.82 |
| 1.32 |
| 0.91 |
| 1.17 |
| 1.55 |
| 0.72 |
| 0.81 |
| 0.5 |
| 0.52 |
| 0.83 |
| 0.76 |
| 1.21 |
| 0.84 |
| 1.07 |
| 0.43 |
| 1.33 |
| 1.05 |
| 0.97 |
| 0.51 |
| 0.61 |

| |
|---|
| 1.12 |
| 0.77 |
| 0.45 |
| 0.45 |
| 0.68 |
| 1.34 |
| 0.66 |
| 0.84 |
| 0.63 |
| 0.82 |
| 0.8 |
| 1.69 |
| 0.69 |
| 0.81 |
| 0.88 |
| 0.5 |
| 1.96 |
| 0.77 |
| 0.5 |
| 0.49 |
| 0.55 |
| 1.01 |
| 1.63 |
| 1.2 |
| 2.27 |
| 1.09 |
| 0.5 |
| 2.1 |
| 1.52 |
| 0.98 |
| 0.77 |
| 0.62 |
| 0.96 |
| 0.98 |
| 0.89 |
| 0.35 |
| 1.95 |
| 1.1 |
| 0.8 |

| |
|---|
| 1.26 |
| 0.7 |
| 1.96 |
| 1.1 |
| 0.93 |
| 0.64 |
| 0.37 |
| 0.54 |
| 1.04 |
| 0.97 |
| 0.65 |
| 0.65 |
| 0.66 |
| 1.07 |
| 0.4 |
| 1.2 |
| 1.49 |
| 0.81 |
| 0.73 |
| 1.06 |
| 0.64 |
| 1.16 |
| 1.09 |
| 0.41 |
| 0.49 |
| 0.42 |
| 0.8 |
| 0.65 |
| 0.42 |
| 0.98 |
| 0.42 |
| 0.84 |
| 0.75 |
| 0.43 |
| 0.62 |
| 0.43 |
| 0.83 |
| 0.66 |
| 1.23 |

| |
|---|
| 0.87 |
| 0.44 |
| 1.13 |
| 1.76 |
| 0.92 |
| 0.66 |
| 0.8 |
| 1.74 |
| 1.16 |
| 1.02 |
| 0.46 |
| 0.65 |
| 0.89 |
| 0.75 |
| 0.61 |
| 1.37 |
| 1.67 |
| 0.4 |
| 0.99 |
| 1.11 |
| 0.64 |
| 0.85 |
| 0.89 |
| 0.81 |
| 0.78 |
| 0.47 |
| 0.47 |
| 0.47 |
| 0.73 |
| 0.48 |
| 0.95 |
| 0.94 |
| 0.75 |
| 0.7 |
| 0.7 |
| 0.97 |
| 0.82 |
| 0.48 |
| 0.48 |

1.21
0.58
0.66
0.72
0.48
0.81
0.85
0.68
0.49
1.2
0.9
0.56
0.98
0.69
0.9
0.49
0.75
1.88
0.49
0.49
0.73
1.08
1.11
0.88
0.54
1.22
0.5
0.83
0.5
1.01
1.45
0.52
0.5
0.6
0.5
0.55
0.74
0.88
0.59

| |
|---|
| 0.91 |
| 1.29 |
| 1.36 |
| 0.5 |
| 0.84 |
| 0.98 |
| 0.5 |
| 0.5 |
| 0.58 |
| 0.64 |
| 1.15 |
| 0.53 |
| 0.79 |
| 0.77 |
| 2.47 |
| 0.79 |
| 0.74 |
| 0.52 |
| 0.51 |
| 0.66 |
| 0.51 |
| 0.87 |
| 1.55 |
| 0.74 |
| 0.51 |
| 0.83 |
| 1.36 |
| 0.51 |
| 0.51 |
| 0.68 |
| 0.51 |
| 0.77 |
| 1.12 |
| 0.78 |
| 0.51 |
| 0.84 |
| 1.33 |
| 1.01 |
| 0.54 |

| |
|---|
| 0.8 |
| 0.53 |
| 0.48 |
| 0.94 |
| 0.4 |
| 0.54 |
| 0.56 |
| 0.55 |
| 2.18 |
| 1.4 |
| 0.56 |
| 0.76 |
| 0.56 |
| 0.5 |
| 0.66 |
| 1.17 |
| 0.58 |
| 0.57 |
| 0.51 |
| 0.57 |
| 0.91 |
| 0.86 |
| 1.43 |
| 1.26 |
| 0.58 |
| 0.57 |
| 0.59 |
| 0.58 |
| 1.51 |
| 0.8 |
| 0.87 |
| 0.59 |
| 0.58 |
| 0.49 |
| 1.04 |
| 0.58 |
| 0.6 |
| 0.47 |
| 0.76 |

| |
|---|
| 0.59 |
| 0.59 |
| 0.73 |
| 1.03 |
| 0.88 |
| 0.6 |
| 0.59 |
| 0.49 |
| 0.84 |
| 0.87 |
| 0.81 |
| 0.61 |
| 0.86 |
| 0.27 |
| 0.52 |
| 1.02 |
| 0.61 |
| 0.27 |
| 0.93 |
| 0.63 |
| 1.06 |
| 0.27 |
| 1.35 |
| 0.64 |
| 0.28 |
| 0.28 |
| 0.28 |
| 0.64 |
| 1.18 |
| 0.63 |
| 0.29 |
| 0.29 |
| 0.28 |
| 0.88 |
| 0.64 |
| 0.83 |
| 0.78 |
| 0.9 |
| 0.29 |

0.29
0.29
0.28
0.29
0.83
0.28
0.61
1.1
1.13
0.3
0.65
0.3
0.3
0.3
0.3
0.3
0.3
0.29
0.66
0.65
0.31
0.59
0.31
0.31
0.31
1
0.32
0.51
0.31
0.31
0.32
0.92
0.65
0.68
0.67
0.67
0.32
0.32
0.32

| |
|---|
| 0.99 |
| 0.68 |
| 0.8 |
| 0.33 |
| 0.33 |
| 0.32 |
| 0.33 |
| 0.32 |
| 0.32 |
| 0.32 |
| 2 |
| 1.09 |
| 0.68 |
| 0.68 |
| 1.47 |
| 0.68 |
| 1.12 |
| 0.69 |
| 0.68 |
| 0.67 |
| 0.68 |
| 0.34 |
| 0.34 |
| 0.34 |
| 0.34 |
| 0.34 |
| 0.69 |
| 0.33 |
| 0.69 |
| 0.34 |
| 0.34 |
| 0.34 |
| 0.34 |
| 0.34 |
| 0.68 |
| 0.69 |
| 0.51 |
| 1.54 |
| 0.84 |

| |
|---|
| 0.6 |
| 0.75 |
| 1.15 |
| 0.34 |
| 0.34 |
| 1.89 |
| 1.01 |
| 0.8 |
| 0.7 |
| 0.5 |
| 0.35 |
| 0.34 |
| 0.34 |
| 0.31 |
| 0.7 |
| 0.69 |
| 0.36 |
| 0.36 |
| 0.35 |
| 0.35 |
| 0.35 |
| 1.02 |
| 0.58 |
| 0.36 |
| 0.36 |
| 0.36 |
| 0.35 |
| 1.03 |
| 0.36 |
| 0.71 |
| 0.94 |
| 0.72 |
| 0.72 |
| 0.56 |
| 0.37 |
| 0.37 |
| 0.36 |
| 0.36 |
| 0.36 |

0.36
1.62
1.03
0.72
0.37
0.37
0.37
1.01
0.37
0.37
0.54
0.36
0.37
0.37
0.37
0.72
0.73
0.55
0.72
0.37
0.37
0.37
1.57
1.18
0.64
0.38
0.38
0.73
0.47
0.38
0.37
0.38
1.02
1.17
0.82
0.72
0.49
0.74
0.73

| |
|---|
| 0.79 |
| 0.38 |
| 0.32 |
| 0.37 |
| 1.51 |
| 0.73 |
| 0.39 |
| 0.39 |
| 0.39 |
| 0.38 |
| 0.38 |
| 0.39 |
| 0.7 |
| 0.7 |
| 0.65 |
| 0.73 |
| 0.39 |
| 0.38 |
| 0.38 |
| 0.39 |
| 0.39 |
| 1.96 |
| 0.89 |
| 1.21 |
| 0.75 |
| 0.74 |
| 0.74 |
| 0.39 |
| 0.39 |
| 0.74 |
| 0.75 |
| 0.4 |
| 0.75 |
| 0.39 |
| 0.72 |
| 0.41 |
| 0.4 |
| 0.4 |
| 0.4 |

2.79
0.76
1.27
0.41
0.41
0.81
0.4
0.67
0.76
0.81
1.18
0.41
0.69
0.32
0.4
0.4
0.67
0.41
0.41
0.41
0.41
1.57
0.75
0.79
0.42
0.42
0.42
0.74
0.41
0.4
0.66
1.2
0.42
2.11
1.43
0.78
0.77
0.42
1.07

| |
|---|
| 0.76 |
| 0.41 |
| 0.42 |
| 0.42 |
| 2.08 |
| 0.83 |
| 1.15 |
| 2.21 |
| 0.72 |
| 0.76 |
| 0.43 |
| 0.42 |
| 0.42 |
| 0.42 |
| 0.42 |
| 0.42 |
| 0.66 |
| 1.26 |
| 0.43 |
| 0.43 |
| 0.43 |
| 0.42 |
| 2.3 |
| 0.77 |
| 1.11 |
| 0.43 |
| 1.03 |
| 0.43 |
| 0.42 |
| 0.78 |
| 0.78 |
| 0.43 |
| 0.43 |
| 0.43 |
| 0.74 |
| 0.43 |
| 1.8 |
| 1.65 |
| 0.44 |

| |
|---|
| 0.44 |
| 0.44 |
| 0.43 |
| 0.43 |
| 0.43 |
| 0.43 |
| 0.43 |
| 0.43 |
| 0.43 |
| 0.44 |
| 1.6 |
| 0.88 |
| 1.12 |
| 0.78 |
| 0.44 |
| 0.44 |
| 0.44 |
| 0.29 |
| 1.69 |
| 1.25 |
| 1.56 |
| 0.79 |
| 1.23 |
| 0.44 |
| 0.79 |
| 0.44 |
| 0.31 |
| 0.79 |
| 1 |
| 0.45 |
| 0.44 |
| 0.44 |
| 0.44 |
| 0.41 |
| 1.2 |
| 0.44 |
| 0.89 |
| 1.51 |
| 0.45 |

| |
|---|
| 0.45 |
| 0.45 |
| 0.44 |
| 0.67 |
| 0.45 |
| 0.45 |
| 0.38 |
| 0.8 |
| 0.78 |
| 0.8 |
| 0.45 |
| 0.45 |
| 0.45 |
| 0.45 |
| 0.98 |
| 0.45 |
| 0.45 |
| 1.91 |
| 0.9 |
| 0.91 |
| 0.9 |
| 0.45 |
| 0.45 |
| 0.8 |
| 0.46 |
| 0.45 |
| 0.45 |
| 0.45 |
| 0.45 |
| 0.44 |
| 0.45 |
| 0.45 |
| 1.68 |
| 1.19 |
| 0.97 |
| 0.8 |
| 0.81 |
| 0.46 |
| 0.81 |

| |
|---|
| 0.46 |
| 0.44 |
| 0.45 |
| 1.2 |
| 0.81 |
| 0.8 |
| 0.81 |
| 0.76 |
| 1.28 |
| 0.74 |
| 0.81 |
| 0.46 |
| 0.46 |
| 0.45 |
| 0.46 |
| 0.81 |
| 0.38 |
| 0.93 |
| 0.81 |
| 0.81 |
| 0.81 |
| 0.81 |
| 0.81 |
| 0.46 |
| 0.46 |
| 0.46 |
| 0.82 |
| 0.82 |
| 0.46 |
| 0.81 |
| 0.43 |
| 0.69 |
| 0.46 |
| 0.79 |
| 1.37 |
| 0.73 |
| 1.23 |
| 0.47 |
| 0.47 |

0.46
0.62
0.46
0.58
0.47
0.41
0.47
0.46
0.81
0.64
0.92
0.74
0.47
0.83
0.47
0.47
0.47
0.93
0.47
0.47
0.47
0.71
0.47
1.48
0.81
1.82
0.47
0.47
0.47
0.47
0.47
0.47
0.47
0.47
0.47
0.47
0.47
0.47
0.69

0.47
0.47
0.47
0
