## Supplemental Tables 3 to 5 for "A new archaeal virus that suppresses the transcription of host immunity genes"

| strain | Genotype | Source / Reference |
| --- | --- | --- |
| 48N | WT isolate | (1) |
| DS_2_ | WT *Haloferax volcanii* | (2) |
| UG613 | 48N *∆pyrE2* | This study |
| UG595 | 48N *∆pyrE2* *∆HLSV1_integrase* | This study |
| UG685 | 48N *∆pyrE2 ∆HLSV1* | This study |
| UG469 | *ᐃtrpA*, *HVO_0894::pyrE* | This study |

**Supplementary Table T3: Strains used in this study.**

**Supplementary Table T4: Oligonucleotides used in this study**

| Primer | sequence (5’-3’) | properties | |
| --- | --- | --- | --- |
| IT748 | CGGGTTCGGACTCGCGCTCGG | | genotype confirmation for the 48N  *ᐃpyrE2* strain, forward primer |
| IT749 | CGGGGCGTTCGAGGTCCAGCG | | genotype confirmation for the 48N  *ᐃpyrE2* strain, reverse primer |
| IT5 | CACCGAGGACGAACTCGAA | | HLSV1 upstream forward primer,  for generating 48N HLSV1 ᐃintegrase |
| IT6 | CCGATTCAGCCTTAATGGGTATGGGACCGCCCGGATTTGAACCGGGGTCACG | | HLSV1 upstream reverse primer,  for generating 48N HLSV1 ᐃintegrase |
| IT7 | TCAAATCCGGGCGGTCCCATACCCATTAAGGCTGAATCGGCCGTTCTTGC | | HLSV1 downstream forward primer,  for generating 48N HLSV1 ᐃintegrase |
| IT8 | CACCGAGTGAGTTCGGGC | | HLSV1 downstream reverse primer,  for generating 48N HLSV1 ᐃintegrase |
| IT1 | CACCGAGGACGAACTCGAA | | HLSV1 upstream forward primer,  for generating 48N ᐃHLSV1 |
| IT2 | CCGATTCAGCCTTAATGGGTATGGGACCGCCCGGATTTGAACCGGGGTCACG | | HLSV1 upstream reverse primer,  for generating 48N ᐃHLSV1 |
| IT3 | TCAAATCCGGGCGGTCCCATACCCATTAAGGCTGAATCGGCCGTTCTTGC | | HLSV1 downstream forward primer,  for generating 48N ᐃHLSV1 |
| IT4 | CACCGAGTGAGTTCGGGC | | HLSV1 downstream reverse primer,  for generating 48N ᐃHLSV1 |
| IT24 | CCTCCTCGAAGCGATAACAG | | virus 48N ddPCR Fw-Amplify  48N virus for ddPCR |
| IT25 | CGAGTTTCTCTCGGGTGTTC | | virus 48N ddPCR Rv-Amplify  48N virus for ddPCR |
| IT556 | CGCGGGAACGACTTTCGAC | | HLSV1 integrated form  -down forward 1000bp |
| IT557 | CCGTCCAGTGGGACGTTATC | | HLSV1 integrated form  -down reverse 1000bp |
| IT558 | AATCAGATGAACGTCGCCCT | | HLSV1 integrated form  -up forward 1000bp |
| IT559 | CGACGATGCCGTGTGTCTAC | | HLSV1 integrated form  -up reverse 1000bp |
| IT560 | GTCGATCACGTGGCATCAG | | HLSV1 forward circular  - 500bp |
| IT561 | ACGAAGGGGAGGTCCGTC | | HLSV1 reverse circular  - 500bp |
| IT270 | GGGTCGACGGAAACGTTGAT | | Forward- used for new  CRISPR spacers acquisitions  in CRISPR array C and D  of *Hfx. volcanii* lab strain |
| IT271 | AATTGGACCCCGGCTTCG | | Reverse- used for new  CRISPR spacers acquisitions  in CRISPR array D  of *Hfx. volcanii* lab strain |
| IT272 | TGTGATTCGATACGCGACAC | | Reverse- used for new  CRISPR spacers CRISPR acquisitions  in CRISPR array C of  *Hfx. volcanii* lab strain |
| IT548 | CCGTACTCAGACCACGACA | | Forward- used for new  CRISPR spacers acquisitions  in CRISPR array A of 48N |
| IT549 | CGTAGTCACCCCTCAGAGAGT | | Reverse- used for new  CRISPR spacers acquisitions  in CRISPR array A of 48N |
| IT550 | GACAATTCGCTCGGTCACG | | Forward- used for new  CRISPR spacers acquisitions  in CRISPR array B of 48N |
| IT551 | TGATTTCGGGACGGTTTCAG | | Reverse- used for new  CRISPR spacers acquisitions  in CRISPR array B of 48N |
| IT552 | GGCTTCGACGGGGATTGTC | | Forward- used for new  CRISPR spacers acquisitions  in CRISPR array C of 48N |
| IT553 | GGGTCGACGGAAACACTCTT | | Reverse- used for new  CRISPR spacers acquisitions  in CRISPR array C of 48N |

**Supplementary Table T5: plasmids used in this study**

| Plasmid | Description | Source / Reference | |
| --- | --- | --- | --- |
| pGB68 | containing the halobacterial novobiocin resistance gene *gyrB* and flanking sequences of *pyrE2* | | (3) |
| pIS71 | pop in-pop out vector used to delete the HLSV1 integrase gene (stock name UG587) | | This study |
| pIS76 | pop in-pop out vector used to delete the entire HLSV1 from 48N genome (stock name UG584) | | This study |
| pWL102 | *Escherichia coli*-*Haloferax volcanii* shuttle vector. confers resistance to ampicillin and mevinolin | | (4) |

1.	**Shalev Y, Soucy SM, Papke RT, Gogarten JP, Eichler J, Gophna U.** 2018. Comparative Analysis of Surface Layer Glycoproteins and Genes Involved in Protein Glycosylation in the Genus Haloferax. *Genes (Basel)* **9**.

2.	**Hartman AL, Norais C, Badger JH, Delmas S, Haldenby S, Madupu R, Robinson J, Khouri H, Ren Q, Lowe TM, Maupin-Furlow J, Pohlschroder M, Daniels C, Pfeiffer F, Allers T, Eisen JA.** 2010. The complete genome sequence of Haloferax volcanii DS2, a model archaeon. *PLoS One* **5:**e9605.

3.	**Bitan-Banin G, Ortenberg R, Mevarech M.** 2003. Development of a gene knockout system for the halophilic archaeon Haloferax volcanii by use of the pyrE gene. *J Bacteriol* **185:**772-778.

4.	**Lam WL, Doolittle WF.** 1989. Shuttle vectors for the archaebacterium Halobacterium volcanii. *Proc Natl Acad Sci U S A* **86:**5478-5482.
