## Supplemental Figure S1 for "A new archaeal virus that suppresses the transcription of host immunity genes"

### 48N virus is a replicating circular virus

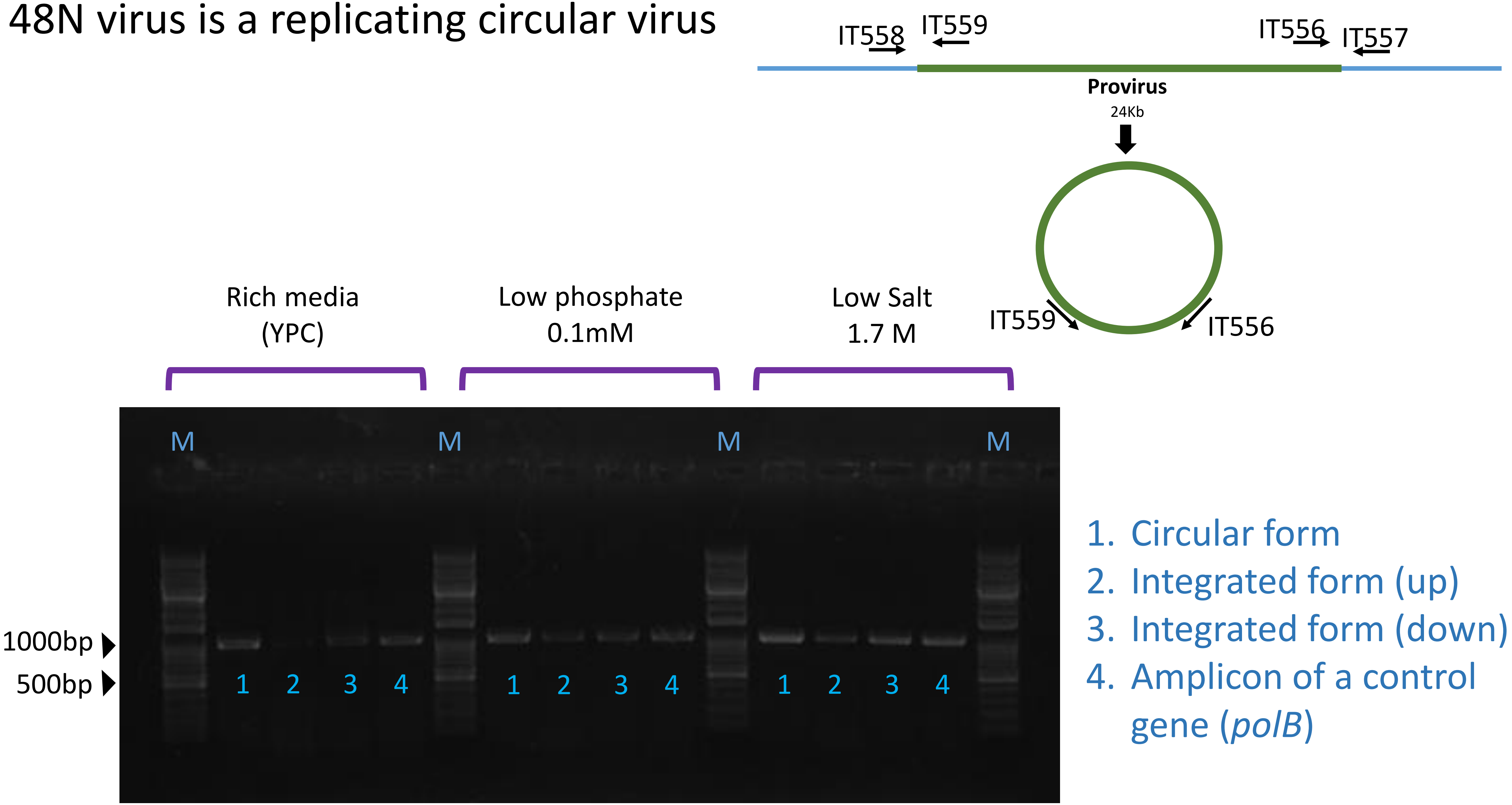


**Supplementary figure S1. Evidence for both replicating and integrated forms of LSV-48N in cells.** Agarose gel electrophoresis of PCR amplicons obtained from liquid cultures of 48N cells grown under different conditions.
