## Supplemental Figure S2 for "A new archaeal virus that suppresses the transcription of host immunity genes"

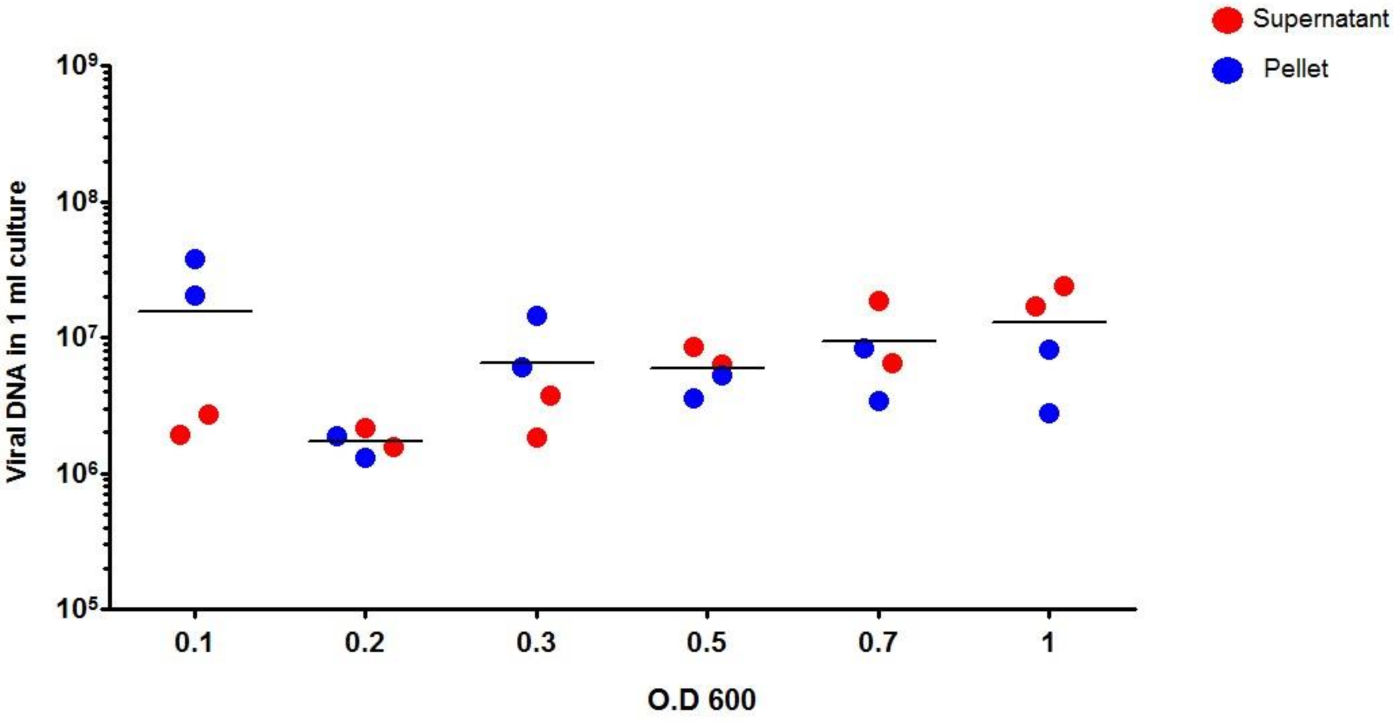


**Supplementary Figure S2.** Digital droplet PCR quantification of LSV-48N genome copies along the growth curve (increasing OD) taken from a culture of 48N grown on rich (YPC) medium.
