## Supplemental Figure S3 for "A new archaeal virus that suppresses the transcription of host immunity genes"

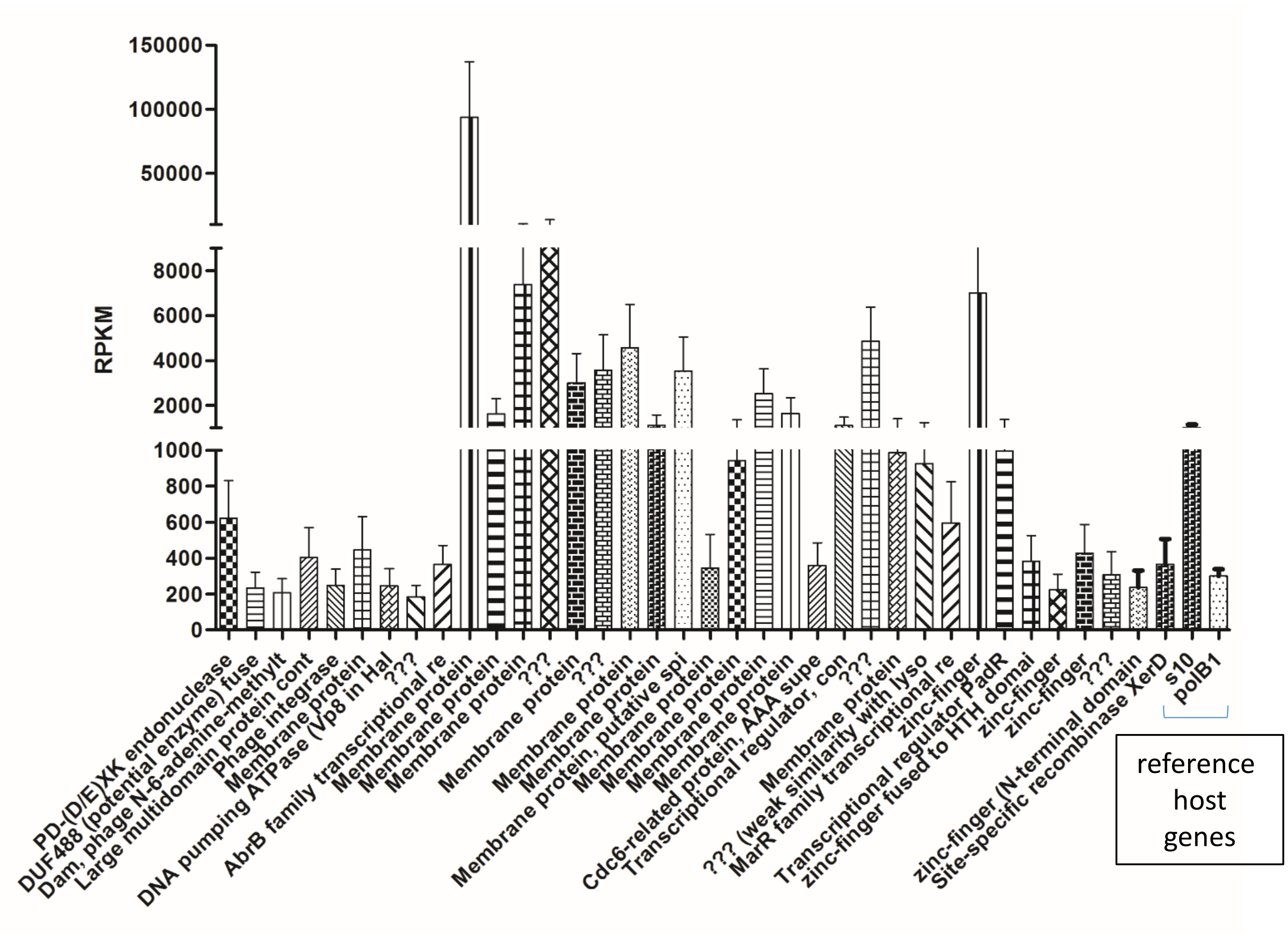


**Supplementary Figure S3.** Mean gene expression levels of the LSV-48N genes in a virus-infected 48N culture based on three 3 independent RNA-Seq experiments. A highly-expressed host gene and a moderately-expressed host gene are provided as a reference. Genes of unknown function are marked as “???”.
