## Supplementary figures and images for "A new archaeal virus that suppresses the transcription of host immunity genes"

### Supplemental Figure S4

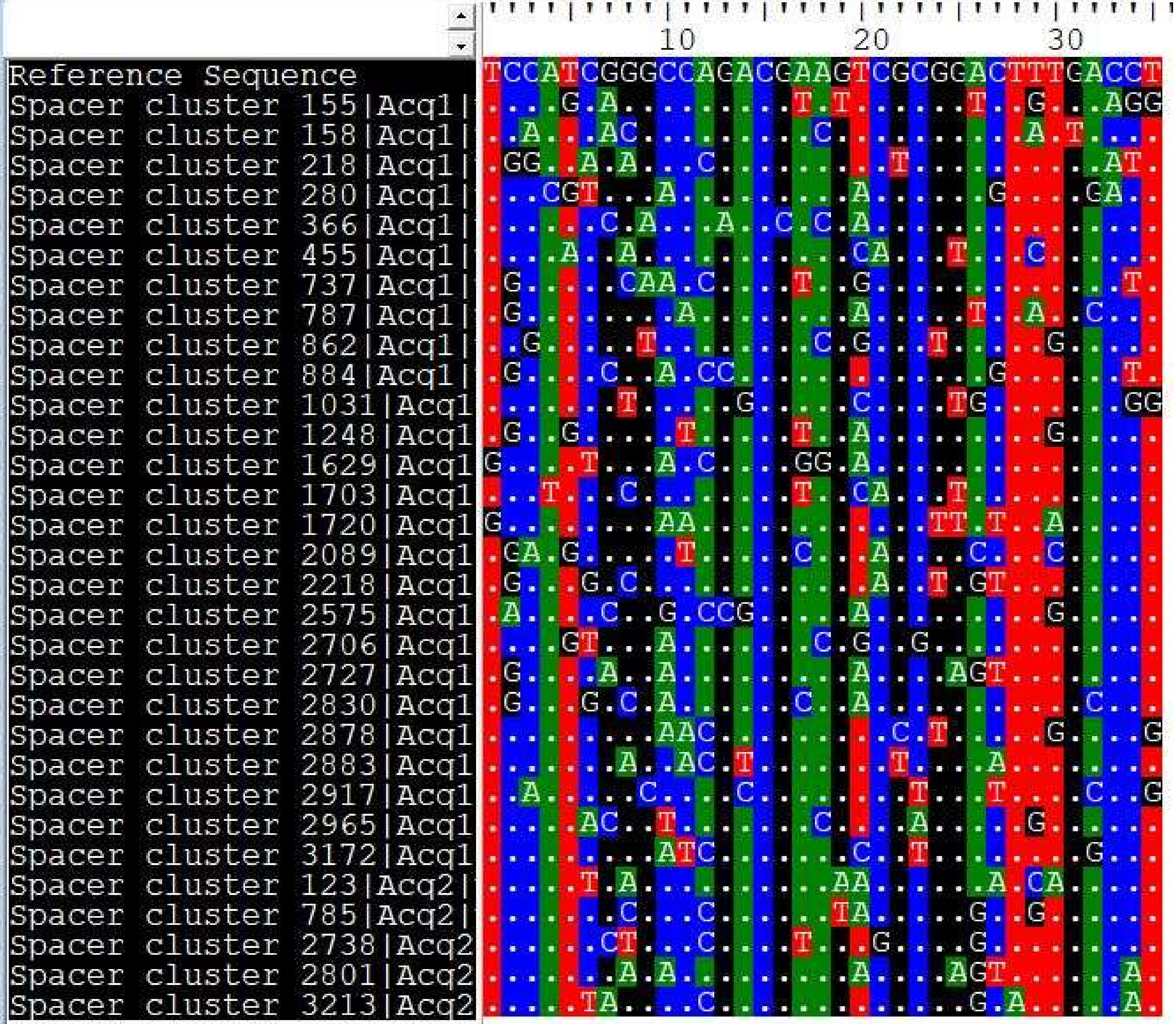
